# Multi-Layer Autocatalytic Feedback Enables Integral Control Amidst Resource Competition and Across Scales

**DOI:** 10.1101/2024.08.22.609155

**Authors:** Armin M. Zand, Stanislav Anastassov, Timothy Frei, Mustafa Khammash

## Abstract

Integral feedback control strategies have proven effective in regulating protein expression in unpredictable cellular environments. These strategies, grounded in model-based designs and control theory, have advanced synthetic biology applications. Autocatalytic integral feed-back controllers, utilizing positive autoregulation for integral action, are particularly promising due to their similarity to natural behaviors like self-replication and positive feedback seen across biological scales. However, their effectiveness is often hindered by resource competition and context-dependent couplings. This study addresses these challenges with a multi-layer feedback strategy, enabling population-level integral feedback and multicellular integrators. We provide a generalized mathematical framework for modeling resource competition in complex genetic networks, supporting the design of intracellular control circuits. Our controller motif demonstrated precise regulation in tasks ranging from gene expression control to population growth in multi-strain communities. We also explore a variant capable of ratiometric control, proving its effectiveness in managing gene ratios and co-culture compositions in engineered microbial ecosystems. These findings offer a versatile approach to achieving robust adaptation and homeostasis from subcellular to multicellular scales.

## Introduction

Engineered genetic devices that regulate intracellular processes hold great promise for applications in biomanufacturing and biomedicine [1–3]. One obstacle to practical applications of synthetic gene circuits is the lack of reliability and predictability, regarding whether these circuits would behave as planned robustly amidst contingencies like altered conditions or uncontrolled environments. This lack of reliability and robustness is partly due to higher degrees of uncertainty and emerging complexities associated with engineered living systems [2–5]. Inspired both by control engineering applications [2, 6] and natural phenomena [3, 4], biomolecular feedback control mechanisms that provide robustness to uncertainty and disturbance rejection have been proposed and tested [7–16]. Due to their desirable adaptation properties, genetic integral feedback control (IFC) designs have been of particular interest.

Genetic IFC has been realized in bacteria [13, 17] and, more recently, in mammalian cells [18–20]. These studies showed that IFC in a closed-loop circuit, under ideal conditions, can equip the controlled output of interest with a property known as robust perfect adaptation (RPA) [3, 4]. RPA guarantees that the target species is tightly regulated about a set-point even in the event of persistent disturbances. This has potential significance in application areas such as bacterial growth-rate control [13, 21], where the regulation of target populations needs to be robust against, for example, temperature or pH variations in the culture media. A lot of recent theoretical studies have paved the way for the characterization of diverse bio-controllers that can achieve RPA, uncovering the design principles that govern their homeostatic behavior, alongside their theoretical properties and potential limitations [22–32].

Autocatalysis, or, in simple terms, ”repeatedly making more of itself,” underpins many fundamental natural processes and intrinsic behaviors of living systems across scales [33–38], from the molecular to cellular and ecosystem levels, and beyond. Epitomized by *self-replication* and *self-reproduction* [33–35], which represent nature’s grand machinery evolved to ensure the continued generation of species, the manifestation of autocatalytic systems, autoregulation, and positive feedback span from ecology [33] to cognitive systems [38], development [37], medicine [39], evolution and the origins of life [33–35], chemistry [35, 36], social behaviors [33], and synthetic and systems biology [40, 41]. This inspiration has driven us to delve deeper into the use of autocatalysis and positive autoregulation for achieving integral control, motivating the scope of our current research. We begin by considering the simplest (minimal) formulation of the autocatalytic IFC motif [32, 41–44]. This minimal formalism, often recognized as the standard model in the literature, is characterized by a single species that utilizes positive autoregulation [41] to effectuate integral action. By leveraging mathematical modeling, we set to initially reevaluate the potential limitations of such a control paradigm, especially in terms of its practical implementation within a growing cell or cell-free system.

Regardless of controller type used, its genetic implementation must be done with great care, as biological parts can affect one another directly or indirectly, leading to unintended interactions that affect regulatory performance [45]. For instance, shared cellular resources can indirectly couple otherwise orthogonal expression systems [46–51]. The modeling of dynamical couplings resulting from resource competition in genetic circuits, with a particular focus on protein translation in bacterial cells, has sparked lots of interest lately [17, 48, 49, 51–56]. So far, the role of context dependence and competition for transcriptional and translational resources in the design of genetic controllers has not received sufficient attention.

In particular, two or more genes competing for the same resources may alter the intended controller topology and lead to degraded performance. Interestingly, this embraces control paradigms that rely on autocatalysis to achieve integral feedback, as autocatalytic production frequently manifests in competitive forms. For instance, consider the plasmid replication in bacteria [57]. While necessary, the presence of a single plasmid copy is not sufficient to ensure replication. Although often omitted for simplicity, this process involves DNA polymerases, Rep proteins, and various other enzymes, which may be limited in quantity and shared between other replicons within a cell. Such shared dependencies and limited availability of necessary resources result in indirect couplings and lead to the emergence of competitive autocatalytic terms. These fundamentally arise from the temporal occupation of limited resources and their subsequent slow release (or consumption). In fact, we show that the minimal representation of autocatalysis-based IFCs loses the capability to retain RPA when resource couplings are present.

This fact is discussed after we provide a mathematical framework for modeling intracellular resource competition in complex genetic circuits. The approach we adopt follows the methodology described in [53], which involves treating the resources as common enzymes shared between different substrates occupying them, and inhibiting their availability in a mutually exclusive manner [58, 59]. Similar resource-limited modeling approaches were previously also described in other publications [48, 49, 51, 59–61]. Intracellular reaction networks are known for their complexity and high dimensionality. While simplified models offer a basic understanding of their functionality in reduced spaces, there is always a drive to refine and expand the models to enhance accuracy and deepen understanding. Our resource-aware modeling framework expands beyond resource-limited zeroth-order and *unimolecular* reactions to include resource-limited *bimolecular* reactions. This expansion allows us to investigate the impact of scarce resources on chemical reactions in greater detail, providing valuable insights and useful tools for the remainder of the article. The presented framework addresses various types of competitive reactions up to the bimolecular level. This includes both catalytic and conversion types of production reactions, such as when gene transcription is limited by a shared pool of RNA polymerases, as well as competitive degradation and sequestration reactions, like when the degradation of proteins is catalyzed by a shared protease. More detailed derivations and further extensions, complemented by biological insights and examples related to the framework and aimed at facilitating its integration, are provided in Supplementary Material.

Multi-layer systems and feedback redundancy, widely used across disciplines such as control theory, network science, robotics, physics, and systems biology, are utilized to distribute tasks, coordinate subsystems, and enhance system robustness and performance, among others (see [62–65] and references therein). Biological examples include the bacterial heat shock response system [66] and the human glucose regulatory system, which involves multiple pancreatic hormones and tissue-level feedback layers [67]. The phenomena and properties observed in multi-layer dynamical processes are often uniquely tied to their layered structure [63], which may not become evident through analyzing the sub-layers in isolation. Recent experimental study [64] has shown that layering two feedback loops together in an engineered circuit in *E. coli* can outperform its single-layered versions by better managing the tradeoff between robustness to uncertainties and control performance.

Motivated by the loss of RPA in the minimal autocatalytic controller due to resource competition, we propose a multi-layer control strategy that still relies on autocatalysis to establish the integral feedback but successfully retains RPA even if the scarcity of shared resources is taken into account. This is achieved through an interplay between an additionally introduced positive autoregulation loop and the original loop, with resource competition serving as the sole means of communication between the two. In Multi-layer Autocatalytic Biomolecular Controllers: An Effective Solution, we provide a more detailed explanation of this new core motif, which we name layered autocatalytic integral feedback. Further, we evaluate the performance of such a control mechanism by applying it to an embedded gene-expression control task, in which a single limiting resource pool is shared between the protein of interest to be regulated and some other genetic modules present in isogenic populations. Although we initially introduce the motif tailored for intracellular reactions and suited for genetic networks in single-strain systems, but this introduced biocontrol topology is versatile in nature and is adaptable for different situations including altered resource competition dynamics.

Multicellular implementations of functional biocircuits has recently garnered attention for its capacity to distribute the computational workload and metabolic demand of complex tasks across a large population of cooperating individuals [68–75]. This involves assigning specific functional roles to different populations in a microbial consortium, allowing each population to specialize in different tasks and, ultimately, improving the efficacy of the overall process [69, 70, 76–78]. In Multicellular Realization of the Layered Autocatalytic Integrator Motif, we explore a potential multicellular realization of our layered autocatalytic integrator in an engineered community and show that the natural growth dynamics of two or more populations competing for common nutrients, modeled by Lotka-Volterra competition models for example, can be exploited to realize the motif. We capitalize on this versatility in implementation, alternating between the intracellular and multicellular facets and potentials of our control mechanism throughout the article and through provided illustrative examples. The co-cultured community species could represent microbial species grown together in a bioreactor or could be different cell types differentiated from the same stem cell line, for instance. In general, they could be any living, self-replicating entities having *ecosystem-level* interactions, although in this article, we primarily consider co-cultured microbial species.

Following this idea, we consider population growth regulation examples, which are well-known problems in biotechnology and bioprocesses [13, 21, 77, 78]. Our proposed control mechanism distributes the tasks of computing, integrating, and storing the tracking error, as well as correcting the control action based on this error, among two cell populations of a multi-strain microbial system. The dynamics brought about by these two community species, acting in concert as the synthetic controller under consideration, construct a *population-level* integral feedback loop across the consortium.

Ratios and ratiometric calculations play a key role in biological systems and possess considerable relevance in the field of medical science, from sugar utilization networks in yeast [79], to BMP signaling pathways in mammalian cells [80], to human immunology [81] and psychological disorders [82]. For instance, the ATP:ADP ratio signifies the free energy released from the ATP hydrolysis process, an essential quantity for various intracellular reactions [83]. As another example, the ratio of regulatory T cells (Treg) to effector T helper cells is implicated in the development of specific autoimmune diseases [81]. Moreover, there is evidence associating the ratio between the proinflammatory T helper type 17 cells and Treg with the incidence of acute rejection in liver transplant recipients [84].

In the field of synthetic biology, researchers have recently been exploring approaches for the design of engineered genetic devices for achieving robust ratiometric control [69, 74, 78, 85–91]. Recent implementations of ratiometric control involves the deployment of a high-throughput optogenetic platform to achieve robust co-culture composition control in a two-strain engineered *E. coli* community [87], and leveraging a synthetic network involving two proteins to perform ratiometric computation in bacteria [86].

In Ratiometric Control using a Layered Autocatalytic Integral Feedback Strategy, we introduce a suitably adapted version of our layered autocatalytic controller that can achieve precise ratiometric control under the constraint of finite, limiting resources. We showcase the controller’s effectiveness through two different application examples, one at the intracellular level and the other at the population level. The former focuses on embedded regulation of gene expression ratios between two different genes co-expressed within the same host, where sensing and action take place at the transcriptional level and involve protein-protein interactions. Meanwhile, the latter demonstrates the application of our controller in co-culture composition control of multi-strain engineered communities.

## Results

### A Mathematical Framework to Model Intracellular Resource Competition

This section aims to lay down a generalized mathematical framework for dealing with the effect of competition for limited resources in a chemical reaction network (CRN) of interest. We shall assume that this CRN, which could potentially be a complex, high-dimensional network of various species interacting through multiple reactions, is given and that it consists of *m* distinct reaction channels ℛ_*j*_, some of which are production reactions. The production of biomolecular species often relies on free resources, those that are available to bind with reactants in order to initiate or facilitate the production of the final product species. Within a cell, the count of these resources may be limited and simultaneously shared among multiple production reactions. For example, the translation of a gene and protein synthesis is usually modulated by the availability of a large group of required resources, such as ribosomes, transfer RNAs, and elongation factors, among others. For ease of use, we often categorize these into a single resource pool. Typically, this translational resource pool is accessed simultaneously by multiple genes, implying that its availability is intricately tied to the dynamics of all genes sharing it.

A parallel can be drawn with certain degradation reactions. For example, during a post-translational proteolysis process (refer, e.g., to the ClpXP-based protease sharing used in [92]), a protease can serve as a degradation catalyst for multiple proteins, being shared among them and available in limited numbers. Sequestration reactions might similarly necessitate the involvement of some shared species to occur. In this section, our primary focus will be on *production* reactions influenced by resources shared across different substrates. However, in Supplementary Material, Section S1.6, we provide a straightforward extension to the resource-limited degradation/sequestration types of intracellular reactions.

Moving forward, let us discriminate between the reactions that are affected by a resource and those that are not. We assume *q* out of *m* reactions are resource-limited, among which *q*_0_ are zeroth-order (also called birth events or constant inflows), *q*_1_ unimolecular, and the rest, *q*_2_, are bimolecular reactions. We label these 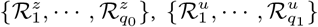, and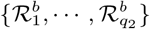, respectively. In the sequel, we assume deterministic models and mass-action kinetics whenever describing intracellular reactions. Throughout, the lower-case letters represent the concentrations of the corresponding species denoted by bold, capital letters.

See Figure 1 for an overview. In this section, we consider resources as general conceptual intracellular species without explicit specification. This way, we maintain our framework’s generality, making it adaptable across genetic circuits of varying scales and components, in which some reaction rates are limited by a shared species. Resources may either represent distinct biochemical species or a compartment comprising various species as a group. The specific resources involved would, apparently, depend on the context, organism, and the biological setting under consideration. Examples could include, among others, transcriptional resources, such as RNA polymerases and transcription factors; translational resources, including ribosomes, aminoacyl tRNAs, and initiation and elongation factors; transmembrane carrier proteins; shared ubiquitin-loaded enzymes; and molecular chaperones.

**Figure 1:**
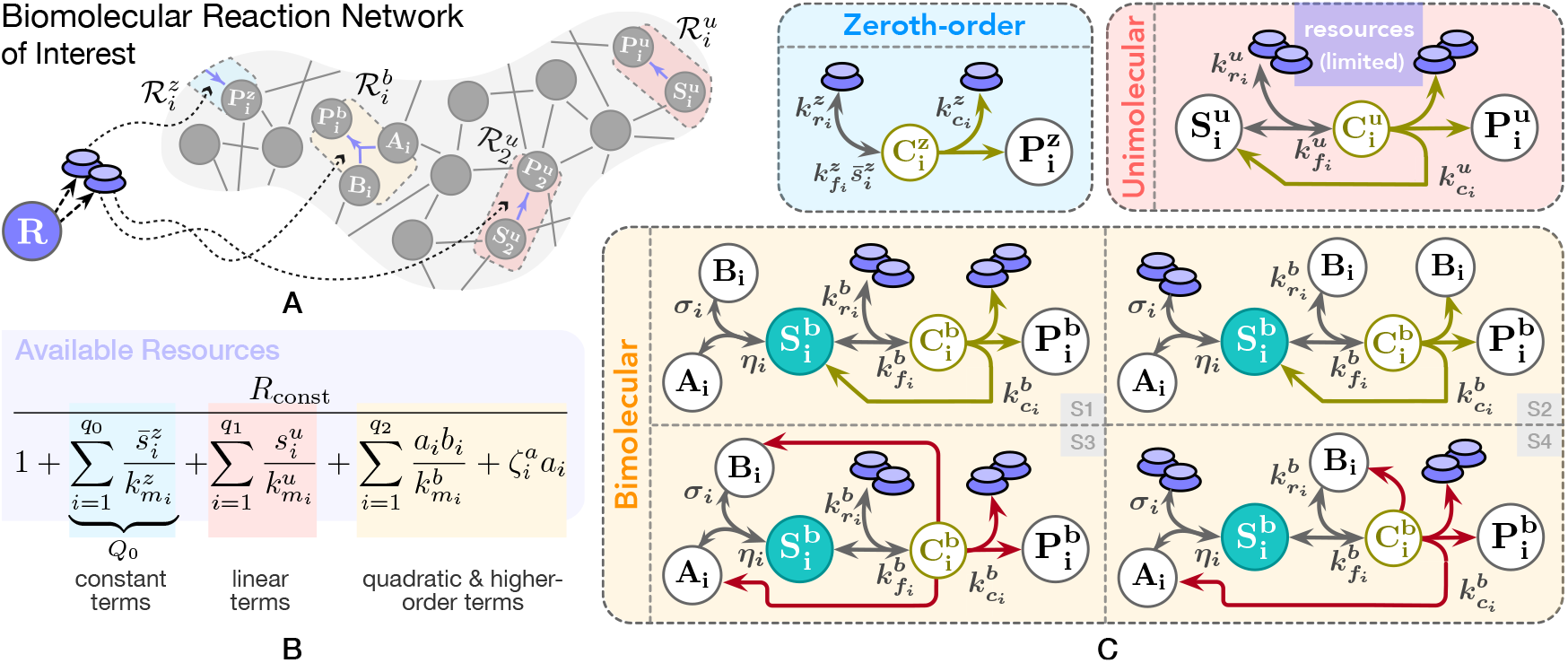
A framework for modeling competition for limited pools of intracellular resources in a given genetic network. (**A**) Each reaction in this network is classified as either resource-limited or not. Resource-limited reactions require the formation of intermediary active complexes between freely available resources (denoted as **R**) and substrate species 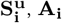, and **B**_i_), which then enable the production of product species (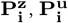 and 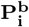). These intermediates, denoted as 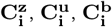 and 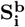, are assumed to be at quasi-steady state, allowing the free resource **R** to be expressed based on substrate concentrations. The cartoon in (**C**) illustrates this process for zeroth-order, unimolecular, and bimolecular catalytic production reactions (four bimolecular scenarios are considered; additional scenarios are discussed in Supplementary Material, Section S1.3). Catalytic production implies substrates act as enzymatic catalysts alongside **R**, enabling their recovery post-reaction. Under certain assumptions (see A1-A3 in the main text), the concentration of the available resources, *r*(*t*), can be approximated by a fraction as in (**B**), where the numerator represents total resource capacity and the denominator consists of substrate-dependent terms, each corresponding to a competing reaction. The extensions of this framework to include resource-limited conversion reactions, as well as degradation/sequestration reactions and reactions involving two different shared resource pools, are detailed in Supplementary Material, Sections S1.4, S1.6 and S1.7.

We shall model the freely available shared resources by an additional species, **R**, whose concentration is affected by all the species that compete for it. Thus, reactions that depend on the availability of free resources will be affected by the concentrations of species which are competing for that resource. For clarity, this article assumes a single limiting resource pool throughout, unless specified otherwise. However, our framework naturally extends to scenarios involving multiple resource pools, provided that no resource-limited reaction draws from more than one pool simultaneously. Extensions to cases with resource-limited reactions constrained by two different resource pools (up to the unimolecular level) are available in Supplementary Material, Section S1.7.

Let *r*(*t*) ∈ ℝ _≥0_ denote the amount of free resources or, equivalently, the concentration of the free resource **R** at time *t*. We take that the pathways from substrates to the product of the reactions 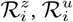, and 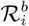 for every *i* conform to the generic forms

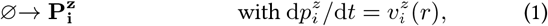

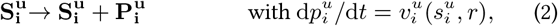

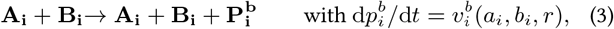

respectively, wherein the effect of resources has not yet been explicitly treated at the reaction level but is taken into account as a limiting factor that affects the production rates. By adopting such representative reaction forms for 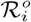 where *o* ∈ {*z, u, b*}, we have indeed restricted the CRN under consideration to resource-limited reactions only of *catalytic* production types, for which reactants (and resources) act as enzymatic catalysts. See Supplementary Material, Section S1.1 for detailed reaction network representations of these three types of resource-limited catalytic production reactions, involving the reaction-level explicit treatment of **R**. Figure 1 provides an illustration of their associated network species, interactions, intermediates, and reaction rate constants. Four different bimolecular scenarios of resource-limited reactions are considered. In Supplementary Material, Section S1.3, we extend the framework to encompass two additional bimolecular scenarios. Though may not covering every possible configuration, the core building-block reactions considered in this framework, from the zeroth-order to all these scenarios of resource-limited bimolecular reactions, enable the modeling of genetic networks with greater detail and permit the explicit treatment of various (often omitted) components, e.g., auxiliary proteins, accessory factors, non-coding RNAs, and more.

Our main goal in this section is to derive, under suitable assumptions, approximate formulae for describing each production rate—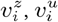, and 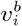—based solely on the concentrations of the substrates 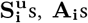, or **B**_i_s that all compete for **R** (detailed derivation is provided in Supplementary Material, Section S1). In reality, the production of a new species typically involves consuming energy, the consumption of additional substrates, or may require potentially irreversible transformation of some existing biomolecules. Here we are assuming that the depletion of such demanded material is minute. Following this, any additional substrates which may be necessary for the production of 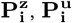, or 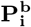 but exist in surplus and not rate-limiting, is neglected in the reactant sides of (1)-(3) for simplicity. Generic examples of such excluded substrates include nucleotides involved in gene transcription and amino acids in gene translation processes, which one may assume that they are plentiful when the cells grow in the exponential phase (not nutrient-starved) and that their depletion during the time interval of interest is negligible, thereby not affecting the production rate 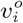.

Figure 2 presents an example of a resource-limited bimolecular reaction, specifically examining the theta-type plasmid replication mechanism in gram-negative bacteria. In addition to the original plasmid slated for replication, this self-replication process pivots on the availability of two other key components to take place: the plasmid replication (Rep) protein and a pool of shared resources **R**. The former acts as the substrate **B**_i_, and the latter encompasses DNA polymerase complexes along with certain other proteins.

**Figure 2:**
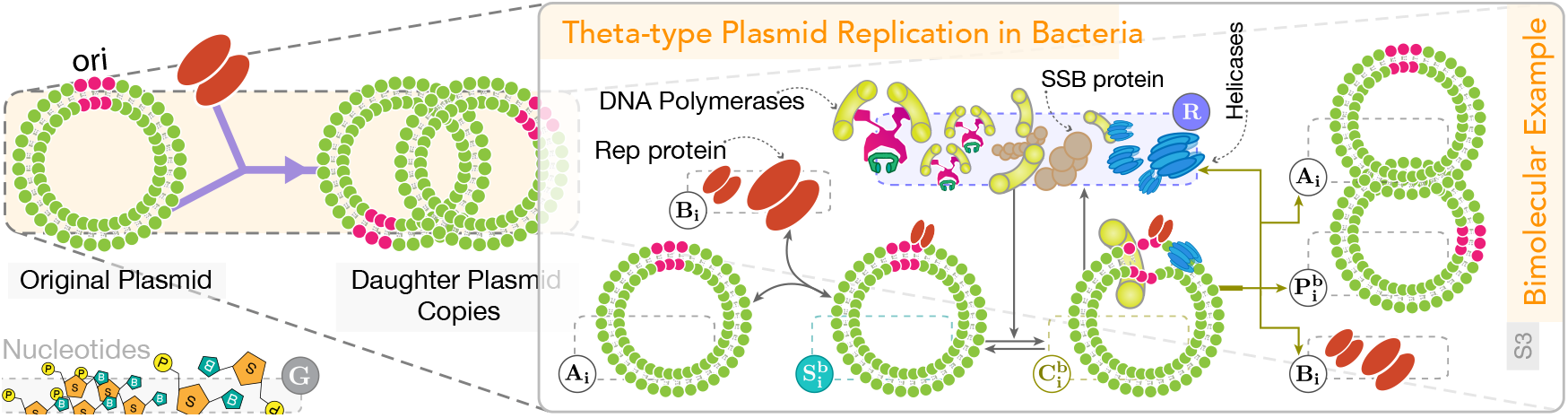
Bacterial plasmid replication framed as a resource-limited bimolecular reaction. This figure illustrates a simplified plasmid replication mechanism in gram-negative bacteria, depicted as a bimolecular autocatalytic reaction constrained by the availability of free resources, collectively represented as **R**, including DNA polymerases, DNA helicases, and DNA ligases, among others. The replication process involves two key substrates: the original plasmid and the Rep protein, the latter acting as a primer for regulating plasmid copy numbers in certain bacteria [57]. By modeling the Rep protein separately as a rate-limiting substrate species, we enable a more detailed mathematical analysis of its role in the competitive formulation of plasmid production framed above. This formulation is particularly relevant in scenarios where, for instance, the synthesis rate of the Rep protein (or its mutants) is upregulated to tune the plasmid replication rate [93]. Other substrates, like nucleotides, are considered abundant and do not limit production rates; these are classified as species **G**. Additional biological insights and examples are provided in Supplementary Material, Section S2.

More detailed examples can be found in Supplementary Material, Section S2. In what follows, we focus only on resource-limited reactions of the catalytic type. Accordingly, we avoid explicitly accounting for the dilution of intermediate complexes due to cellular growth and division, as this would lead to the gradual loss of reactant species (see Section S1.5). This dilution effect might already be negligible in certain contexts, for example, if the cells were cultured in cell lines where cell growth is known to be comparably slow, in cell-free systems, or if all other reaction rates involved in the dynamics of the intermediate complexes are comparably fast to the dilution rate. Separately we account for the effect of diluting intermediates in Supplementary Material, Section S1.5. The modeling of resource-limited *conversion* reactions, in which one or more of the reactants are gradually depleted over time, is outlined in Supplementary Material, Section S1.4. Note that the treatment of resource-limited conversion reactions and resource-limited degradation/sequestration reactions is nearly identical. Therefore, in Supplementary Material, Section S1.6, we derive the latter from the former by induction.

We focus on situations where the amount of available resources is limited, yet the total quantity of resources remains conserved. In these situations, some resources are engaged by the complexes 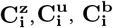, or the 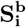 complexes in the second and fourth scenarios, while the remainder are freely available to bind. We imply that the resources remain occupied by the complex 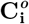 until the product 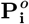 is formed, after which they are released and become available again.

Let us make the following assumptions on the resource-limited reactions considered hereafter.

**Assumptions**. *The following conditions hold for every resource-limited reaction in the considered CRN:*

A1 *For every i, the resource-binding complexes* 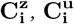, *and* 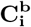 *together with the intermediate complexes* 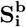 *are all at dynamic equilibrium. This means they rapidly equilibrate to their steady-state values, given changes in the concentration of their constituents*.
A2 *Each bimolecular reaction* 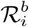*of the third scenario satisfies* 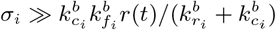.
A3 *Each bimolecular reaction* 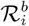 *of the fourth scenario satisfies* 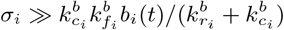.

We would like to remark that the above assumptions are often met in biomolecular resource-substrate interactions, as the last production step, involving the rate constant 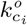, is usually regarded as the slowest step during which the product 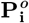is being gradually created until the occupied resource molecules leave the reacting system and become available again. The first assumption is in particular the case if the reactions forming 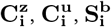, and 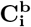 are instances of binding/unbinding reactions that occur on a much faster time scale than the time scale at which the concentrations of the substrates 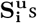, **A**_i_s, and **B**_i_s evolve.

Under the main assumptions above, we find that 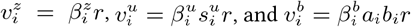, where

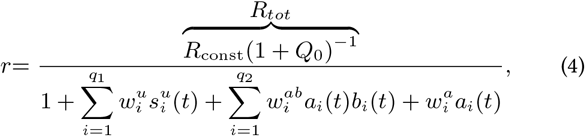

and 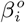 are some positive constants. *R*_const_ is defined as a positive constant, reflecting our initial assumption of conserved total resources. This framework can be extended to accommodate resource-limited reactions with non-static total resource levels—such as those depleting over time—by redefining *R*_const_ as a time-dependent state variable, *R*_const_(*t*). For instance, if the species **R** under study is defined as being some shared consumable for which there tends to be no regeneration system running, e.g. amino acids or nucleotides in cell-free systems. Needless to say, convoluted nonlinearities can arise in the formulation of *r* and 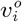 if substrates like 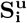 or **B**_**i**_ are treated as quasi-steady-state solutions of an upstream subnetwork.

Rather, the quadratic terms in the denominator of (4) arise from the structural patterns and reaction-level arrangements of the bimolecular scenarios, and may develop into higher-order nonlinear forms in extended scenarios (see Supplementary Material, Section S1.3).

The non-negative scalar *Q*_0_ in (4) indicates the overall contribution of zeroth-order reactions 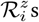 to *r*(*t*). This scaling factor, which by definition remains constant over the time interval of interest, together with the new constant *R*_*tot*_ ∈ ℝ _*>*0_, which may be thought of as the *effective* total capacity of the resource pool, are employed in the way represented by (4) so as to reduce the dimensionality of model parameter space. The constants 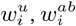, and 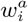 are defined to simplify the notation. They are lumped scalars in terms of the involved reactions’ rate constants, inversely scaled by a factor of 1 + *Q*_0_. We will refer to them as the *competition gains*. They are positive scalars by definition, the only exception is that 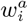 is defined to be zero for the bimolecular reactions of the first and third scenarios. These competition gains should be regarded as independent, unknown constants in the system model, with values that are generally distinct from one another and different from 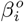.

### The Minimal Autocatalytic Integral Feedback Motif Cannot Mitigate the Effect of Resource Sharing

One of the simplest biochemical reaction network motifs for realizing integral feedback and achieving RPA in closed-loop control is through autocatalytic feedback. In its minimal form, autocatalytic feedback comprises a single controller species that features positive autoregulation and is catalytically inhibited by the regulated output of interest. We provide the schematic illustration of this motif in Figure 3 and a brief discussion in Supplementary Note 1.

**Figure 3:**
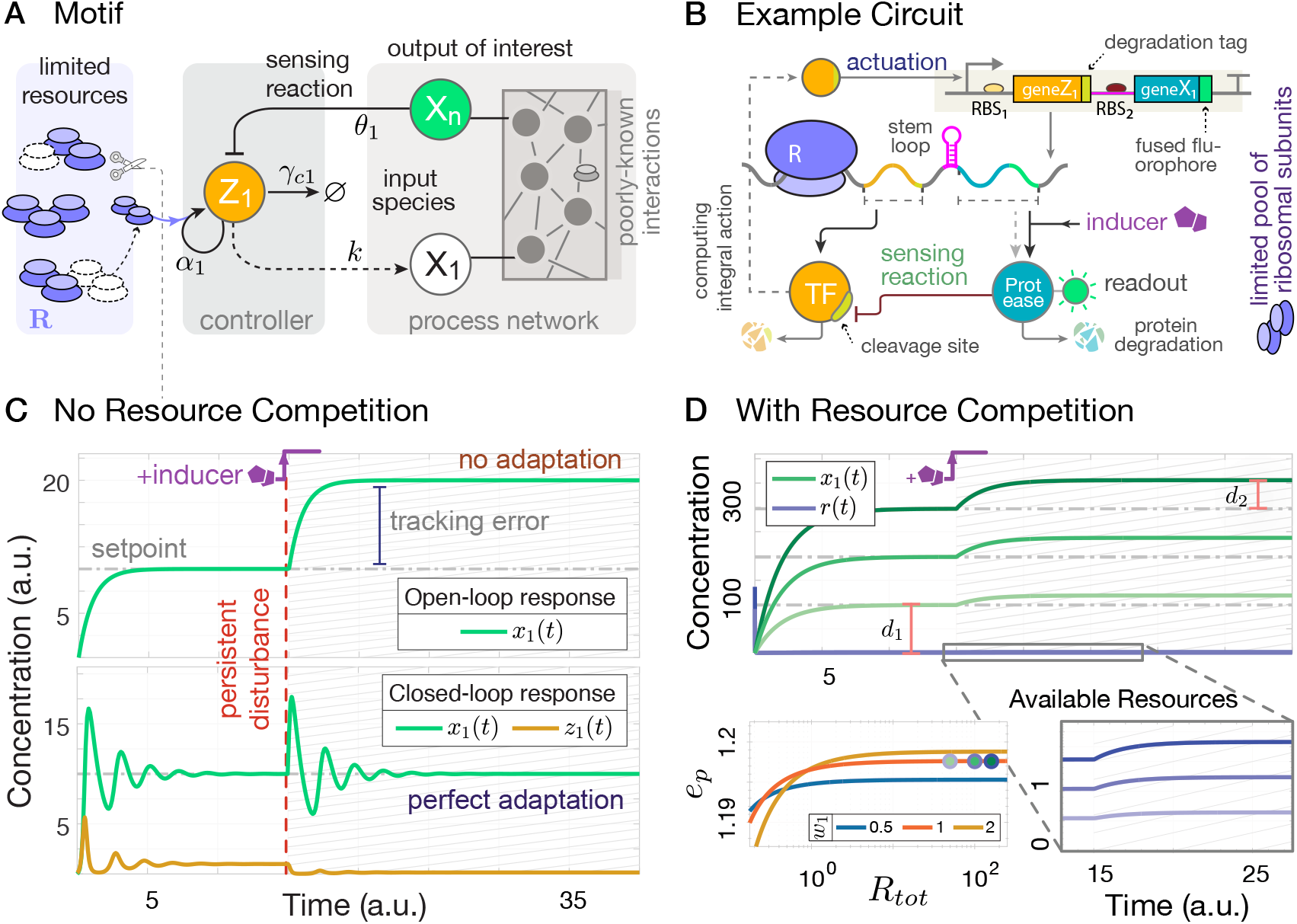
The minimal autocatalytic IFC mechanism no longer achieves RPA in a resource-limited setting. (**A**) Schematic representation of this minimal autocatalytic control scheme controlling a poorly-known reaction network (a.k.a process network). (**B**) An example biomolecular circuit following the control structure in (**A**). The figure depicts a simplified single-species protein expression model. An activating transcription factor fused to a degradation tag forms the protein complex **Z**_1_. The species **X**_1_ is a complex formed by a fluorescent protein, serving as the readout, combined with a fused protease. The synthesis rates of **Z**_1_ and **X**_1_ can be independently tuned via ribosome binding sites (RBS_1,2_). This promoter-selective transcriptional activation directly transfers the control signal, though in scenarios where resource competition is prominent, the controller may also indirectly influence the process through shared resources. (**C**) Sample closed-loop response of the example circuit in (**B**), obtained from the dynamic model (S70) in Supplementary Note 1 with some preset parameters, where the controller does not compete for resources. The degradation tag in **Z**_1_ recognizes **X**_1_’s protease, modulating **Z**_1_’s degradation rate through a second-order reaction. This controlled circuit achieves RPA against external disturbances, such as an abrupt increase in **X**_1_ expression induced by chemical inducers at time *t* = 15. (**D**) Sample closed-loop response of the circuit in (**B**) under resource-limited conditions: The production of **Z**_1_ now depends on the availability of limited resources **R**. As shown for three different values of *R*_*tot*_, the system exhibits imperfect adaptation after a 20% increase in *k*^*\**^ at *t* = 15. The error *e*_*p*_ := (*d*_1_ + *d*_2_)*/d*_1_ measures the normalized deviation of the steady-state value post-perturbation. See Supplementary Note 6 for further details and parameter values.

Under certain standard assumptions, such as set-point admis-sibility and closed-loop stability, this control circuit realizes a (constrained) IFC mechanism in non-competing scenarios when the loop is closed. As noted in Supplementary Note 1, the resulting closed-loop dynamic model in a competitive scenario follows

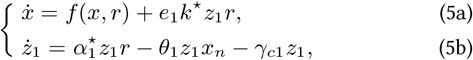

where

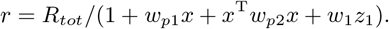

The aggregate competition gains 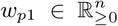 and 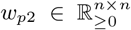 are associated, respectively, with the linear and quadratic terms that arise from the competition for the shared resources **R**, solely from the process side (let us assume that the bimolecular cases, if any, fall under the first scenario).

Note the notation superscript “*\**” used to distinguish between the rate constants of resource-limited reactions—the rate constants 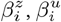 and 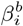 noted in the previous section—and regular (resource-unlimited) ones. This way we explicitly mark the rate constants that have undergone alterations after accounting for a resource-limited setting a nd i mplicitly e mphasize that these a ltered r ate constants may have totally different values from the ones before (in the previous, resource-unlimited setting). Consider these altered rate constants, when scaled by the factor *R*_*tot*_, to be equivalent to their corresponding resource-unlimited rate constants. In this arrangement, the dynamics within a resource-limited setting would, mathematically speaking, approach those of the resource-unlimited counterpart in the limit the competition gains go to zero.

According to the above model, the steady-state 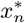, assuming closed-loop stability, now depends on the process parameters, implying that RPA is no longer ensured. This is also confirmed by the simulation results in Figure 3. We note that the concentrations and time are presented in arbitrary units throughout this article, unless the units are explicitly mentioned. The example genetic circuit provided in Figure 3B follows a simplified (single-species) protein-expression model as its process network, with 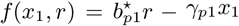 describing its internal dynamics. Here, the resource species is taken to be translational resources, mainly ribosomes produced from the cell’s nucleolus with total number of them being assumed to be conserved (over the time interval of interest). The closed-loop circuit is composed of two distinct species, **X**_1_ and **Z**_1_. The latter is comprised of a transcription factor (TF) fused to a degradation tag. **X**_1_’s gene recruits **Z**_1_ as an activating TF. This gene is inserted in an operon to be co-expressed with the sequence representing **Z**_1_.

Perturbations are applied to the actuation gain and are introduced via the addition of an external chemical inducer to the cell culture medium. The stem loop within the polycistronic mRNA strand is specific to this inducer, exhibiting high affinities that enable the translation of **X**_1_ according to the present inducer’s level. For that purpose, this stem-loop construct could include, for example, an RNA aptamer sensor (see [94] for an example). The leaky expression rate of **X**_1_, or equivalently the basal rate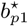, can be modulated by adjusting the tightness of the RNA aptamer. As shown in Figure 3D, increasing the number of total resources alone does not help with restoring the integral action, whereas doing so in combination with decreasing the competition gain does.

### Multi-layer Autocatalytic Biomolecular Controllers: An Effective Solution

As previously demonstrated, the minimal autocatalytic IFC mechanism fails to achieve RPA in the presence of competition for one or more resource pools. This is true even if the process under control does not compete with the controller species and when only **Z**_1_ limits resource availability. Reported in Supplementary Note 2, we systematically explore various variants of the autocatalytic feedback motifs, initially identifying those capable of RPA in the presence of competition for shared resources. These variants typically include an auxiliary autocatalytic feedback layer in addition to the main control layer that features the minimal realization of the autocatalytic IFC. The additional layer may interact directly or indirectly with the rest, the latter through resource coupling. Indirect interactions suggest that the layer is isolated from the main control loop unless it is engaged in the resource competition.

Our search primarily focuses on these indirect cases. The basic idea is to add an auxiliary controller species **Z**_2_, whose main task is to regulate the available resources via *buffering* them. The specific design of its dynamics dictates how the buffering mechanism functions. Particularly, we consider three types of buffering mechanisms: no buffering, *passive* buffering, and *active* buffering. Among the considered variants in Supplementary Note 2, we have selected one to highlight in this section as our proposed solution, based on three key criteria: its ability to dynamically balance resource availability, termed as active buffering; its operating range not being constrained by the controller parameters; and its architectural simplicity, which minimizes additional implementation complexities while fulfilling the first two criteria. This select multi-layer autocatalytic feedback motif successfully overcomes the limitations of the minimal auto-catalytic controller and is capable of achieving RPA amidst resource couplings. In what follows, we present this novel autocatalytic integrator in detail and discuss how it functionally operates, all tailored to intracellular schemes.

With this controller motif closing the loop, Figure 4A shows the schematic of the resulting control system. We provide in Supplementary Note 1 the associated CRN representation and dynamic model, the latter obtained by incorporating our resource-aware framework. As noted therein, in the absence of resource competition, our proposed solution does not offer any improvement compared to the minimal autocatalytic integral feedback. Taking into account the effect of resource competition, however, arranges for the following model for the closed-loop system

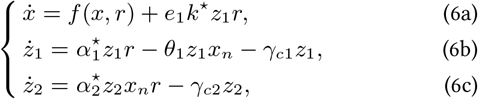

wherein

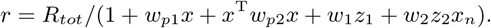

**Figure 4:**
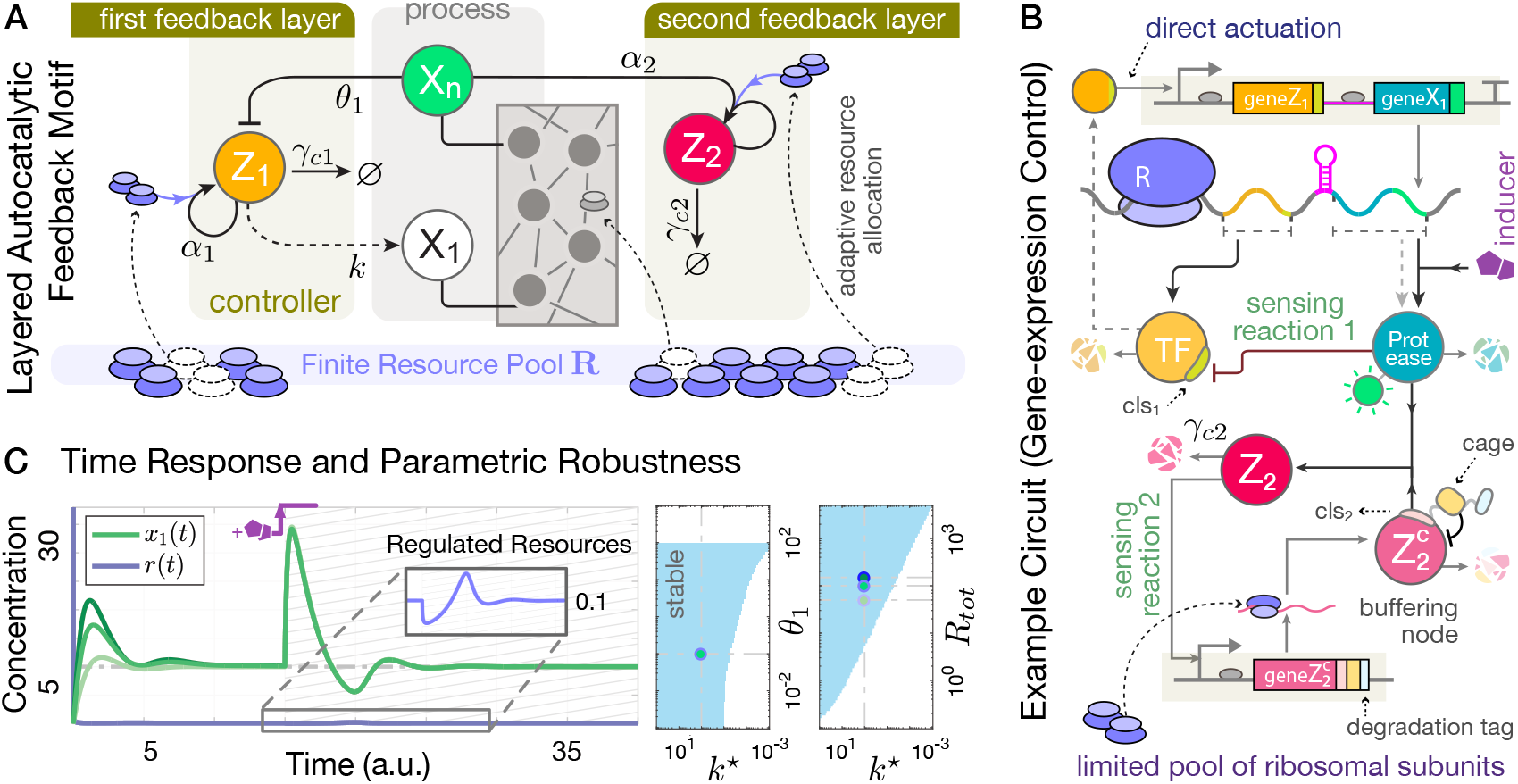
The proposed autocatalysis-based biomolecular controller restores RPA in the presence of resource competition. (**A**) Circuit illustrating the proposed multi-layer autocatalytic IFC strategy. The second layer functions as an implicit feedback loop, where the species **Z**_2_ influences **X**_n_ by buffering shared resources. By actively releasing or occupying resources as needed, **Z**_2_ affects the production of **Z**_1_, thereby modulating the overall abundance of **X**_n_. (**B**) The protein-expression control example adapted from Figure 3B, where the layered autocatalytic controller compensates for resource competition. The process circuitry and species **Z**_1_ remain unchanged, while a new species, 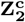, is introduced. 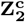 consists of a transcription factor (TF), a cleavage site (cls) for protease degradation by **X**_1_, protein cages, and a separate degradation tag targeting a specific protease (not shown). The protein caging process inhibits the functionality of the cargo TF until it is cleaved, allowing for controlled release and activation. The 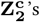 promoter recognizes the released TF (**Z**_2_) as an activator. This mechanism represents a higher-order implementation of the layered autocatalytic motif considered in (**A**). See Supplementary Note 3 for additional information. Even with transcriptional resource competition, the controller maintains its function and exhibits RPA. Further details and conditions ensuring this higher-order model aligns with the reduced model in (6b)-(6c) are provided in Supplementary Note 3. (**C**) Simulation results: At *t* = 15, a disturbance increases the actuation gain *k*^*\**^ from 1 to 50. The controller effectively regulates both the available resources and the output species. The model is based on (8) with 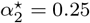 and *γ*_*c*2_ = 0.25, with other parameters as in Figure 3 (*w*_1_ = 1, *w*_2_ = 1, *R*_*tot*_ = 100). Initial conditions are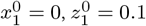, and 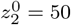. The right panel in (**C**) shows local stability regions from numerical evaluations in two different two-dimensional parametric spaces.

The introduction of intracellular resource competition among controller species, as symbolically depicted in Figure 4 using color-coded arrows, leads to the indirect couplings between the species **Z**_1_, **Z**_2_, and the rest of the reaction network. The role of the additional autocatalytic loop is to leverage these couplings to reinstate the lost RPA property. To see this, we use our resource-aware model (6), to shed light on the functionality of the additional controller species **Z**_2_. Solving (6c)-(6b) for equilibria gives 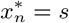 with

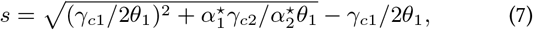

as the only equilibrium point whose value is solely dependent on the controller parameters alone. Hence, so long as the stability of this *desired* equilibrium is preserved, one might expect to observe RPA at the output level *x*_*n*_(*t*). Note that, from the special form of dynamics in (6b) and (6c), it implicitly entails the concentrations of **Z**_1_ and **Z**_2_ to be bounded away from their absorbing state at zero. This condition is a prerequisite for the desired equilibrium satisfying the equality in (7) to be reachable from its neighborhood, and we will always implicitly assume it within the time interval of interest—including the initial conditions—and whenever speaking of the stability of this equilibrium point.

Indeed, given the closed-loop stability of such a desired equilibrium, the steady-state value of *x*_*n*_(*t*) will be maintained at 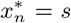 specified by (7) despite constant disturbances in the regulated process or resource-related parameters, including the competition gains and *R*_*tot*_. This holds true even if the explicit formula for *r* is altered. That said, the expression derived in (7) remains unchanged if we were to consider different dynamics for resource couplings, including scenarios with non-static total resources, where *R*_*tot*_ depletes over time, for example. Of course, the existence of a feasible equilibrium point and its stability must be checked for, regardless.

According to our model, the extra autocatalytic loop shaped by **Z**_2_ acts as a buffer that compensates for the effects of resource competition. The coupling between **Z**_1_ and **Z**_2_ through the shared resources realizes two interconnected (constrained) integral feedback loops that work in tandem to maintain the available resource *r*(*t*) at a constant value. It is this resource coupling, therefore, that equips the controller to robustly regulate the species **X**_n_ in the face of process uncertainties and unpredictable tolerances in resource availability. This is illustrated by the numerical results presented in Figure 4C. It is worth mentioning that the controller presented here represents a core, fundamental autocatalytic motif for achieving RPA in competitive settings, for which higher-order implementations are certainly conceivable. A class of such is explored in Supplementary Note 3, an example is given in Figure 4B.

Of note, our controller can automatically manage the available resources while regulating the gene of interest, without the need to introduce extra reactions or dedicate additional controllers. The value at which these available resources are regulated is inversely proportional to the set-point 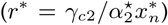. The fact that (7) is encoded via several controller parameters allows flexibility for separate tuning of *r*^*^ and *x*^*^. Indeed, one can vary the ratio 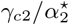 to tune *r*^*^, while keeping the set-point fixed by fine-tuning 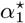 such that 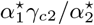 remains unchanged. This parallel regulation may be considered advantageous in certain applications. E.g., in [95] a dedicated integral controller is employed to robustify the ribosome availability as a strategy to control genetic burden. This means that a decrease in available ribosomes, triggered by the activation of disturbance genes, would be offset by the controller, thereby preventing changes in the expression of heterologous genes of interest.

We opted for a resource-limited bimolecular formulation over a unimolecular one to deal with the competitive autocatalysis of **Z**_2_. This choice avoids globally regulating the resources to a fixed value (a *passive* buffering mode), instead allowing for dynamic regulation of available resources according to the control objective, thus providing greater flexibility rather than confining **R** to a preset value. Enabled by the *active* buffering role that the species **Z**_2_ plays, such a dynamic resource allocation strategy can ensure that when high levels of **X**_n_ are not needed (in low set-point regimes), more resources remain available for other circuits, minimizing interruptions to the functionality of the cell’s other (endogenous) genetic modules that also draw from these resources.

Conversely, when high production of **X**_n_ is a need, the controller tends to prioritize **X**_n_’s production, allocating more resources to it. Although this may inevitably limit the expression rates of other circuits, the strategy still offers more flexibility compared to a fixed regulation of **R**, which often lacks easy tuning options. Additionally, if any of the interfering genetic modules suddenly demand more **R** for functioning, the controller would release buffered resources to mitigate disturbances in resource availability. This response is effective as long as the spike in demand remains within the buffering capacity of the controller. Notice that for this integral feedback topology to work, the degradation term *γ*_*c*2_ essentially needs to be non-zero. In intracellular scenarios, there are different approaches to address this concern. For example, choosing **Z**_2_ in a way that it can sequester another auxiliary protein available in excess, or using proteases to catalytically target **Z**_2_ at a controlled rate. These are effective if **Z**_2_ is intended to represent a protein. In the latter, assuming the protease is highly specific to **Z**_2_, there is no competition for it, and sufficient **Z**_2_ is present to operate in a saturated regime, a first-order degradation rate law with a constant *γ*_*c*2_ will remain a valid approximation.

### Embedded Gene-Expression Control in the Presence of External Resource Loads: A Case Study

In this section, we consider a closed-loop circuit where the proposed controller in Multi-layer Autocatalytic Biomolecular Controllers: An Effective Solution acts on a gene-expression plant in the presence of *L* other different genetic modules (*L* ∈ ℕ _≥1_). The modules, each driving the expression of a different co-expressed protein, are referred to as *external resource loads*, which rely on the same resource pool and are (indirectly) connected to the target protein (only) through resource sharing. The controller circuitry, the target protein, and these genetic modules present in the competition are assumed to be all in an isogenic cell population, implying an embedded gene regulation task.

The closed-loop dynamic model of this control system can be expressed by the equations

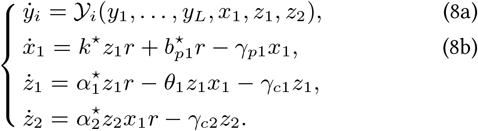

Note its controller dynamics, which are kept the same as those specified in (6b)-(6c). Here,

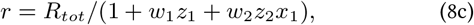

*i* ∈ {1, …, *L*}, and each 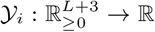 is a continuous function describing the internal dynamics of the *i*th module. The dynamics of the species **X**_1_, representing the target protein to be regulated, are modeled by a simple birth-death process whose birth reaction is resource limited and competes with **Z**_1_ and **Z**_2_ for the resources **R**. The genetic (load) modules are coupled to our controlled circuit only through shared resources. The only assumption we impose on the modules is that *∂*𝒴_*i*_*/∂r* is constant over time for every *i*, or, equivalently, they incorporate resource-limited reactions only of zeroth-order type. In such a scenario, the overall effect of modules on the dynamic evolution of controlled circuit reflects only through the lumped parameter *Q*_0_. This general context is particularly fitting for showcasing the dynamical couplings that arise when a target synthetically inserted gene, encoding for **X**_1_, competes for translational resources with the cell’s endogenous genes (including housekeeping genes, for instance) or with other heterologous genes present within the same cellular environment. The control objective is to, by acting on the transcriptional level, robustly steer the concentration of **X**_1_ to a desired set-point *s*, specified by (7), while rejecting (constant) disturbances that may even arise now from changes solely in the other modules’ internal dynamics.

The controller is expected to reject constant disturbances on the scaling parameter *Q*_0_ as long as the stability of the desired equilibrium is preserved. Recall that step-like changes on *Q*_0_ is equivalent to instant scaling of the competition gains *w*_1_ and *w*_2_ and also the parameter *R*_*tot*_. A sample circuit is illustrated by Figure 5, where each genetic module is assumed to follow a birth-death process 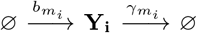 whose basal expression rate is resource-dependent, i.e. 𝒴 _*i*_ is simply taken as 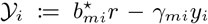. Assuming that all the mRNA components are at quasi-steady state and only considering for translational resource couplings, the dynamics arising from this sample circuit will follow the dynamic model in (8). These considerations may be especially relevant in exponentially growing bacteria, where the competition for translational resources, rather than transcriptional resources, is stipulated to play a more dominant role in gene expression [17, 48, 56, 95].

**Figure 5:**
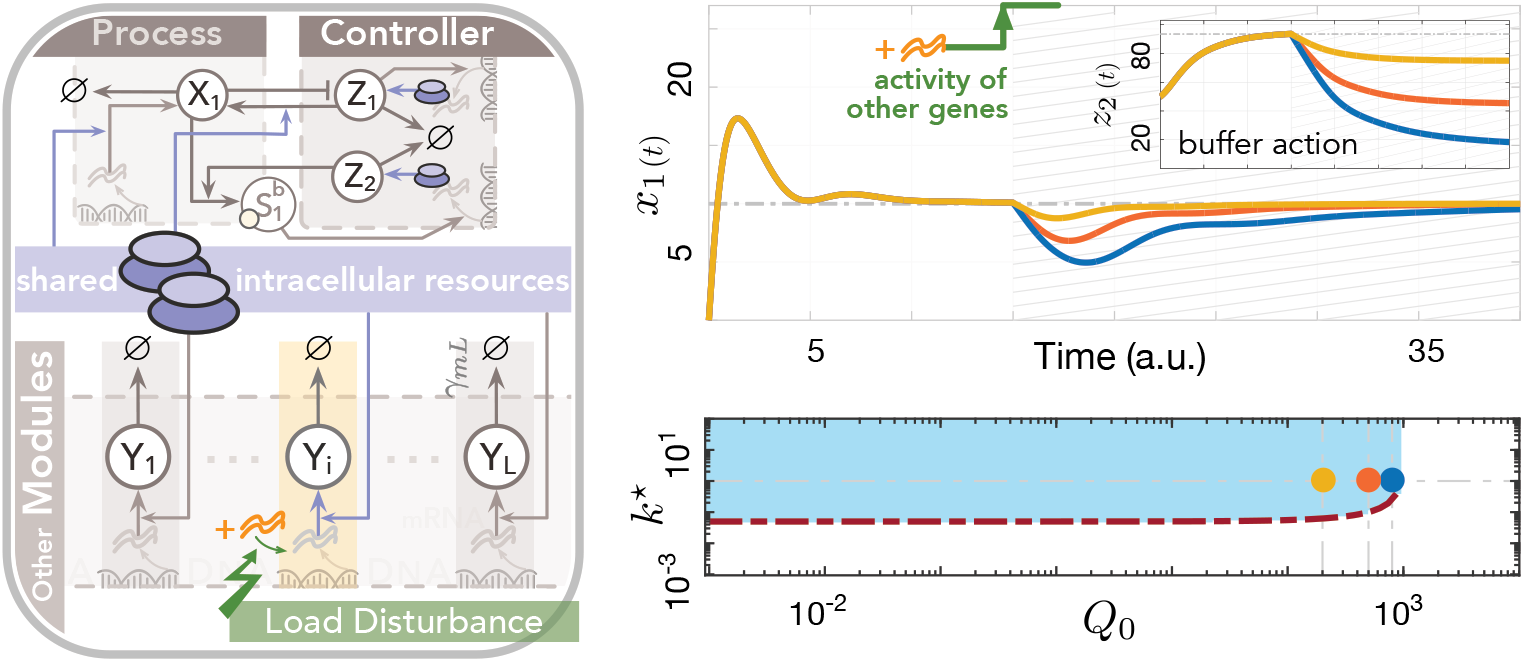
Embedded gene-expression control in the presence of shared translational resources using a multi-layer autocatalytic feedback strategy. The proposed layered autocatalytic feedback motif, embedded within the same cell population as the process it regulates, ensures precise control of a synthetic gene’s expression within a specified range. This process involves not only the synthetic gene but also other genetic modules within the cell that either directly interact with the gene or indirectly influence its expression by sharing translational resources, labeled as species **R**. A dynamic system representing this closed-loop circuit follows the model in (8) with 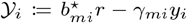 for each *i* ∈ {1, …, *L*}. Increasing the actuation gain *k*^*\**^, as long as the set-point remains admissible, helps reject resource-load disturbances, such as a sudden increase in downstream gene transcription. These increases are modeled as step-like changes in the resource parameter *Q*_0_. Note that a sudden, step-like change in *Q*_0_ within the time interval of interest does not affect resource settings or the reference set-point in (7). The pre-perturbed nominal value for *Q*_0_ is 10^−3^, with other parameters consistent with Figure 4C. The two conditions 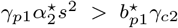 and *R*_*tot*_ *> r*^*^ + *w*_1_*γ*_*p*1_*s/k*^*\**^ ensure the desired equilibrium (*q*_*d*_) is feasible. The boundary determined by the latter condition, 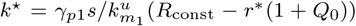, is shown by a dash-dotted red line. The light-blue area indicates the locally stable parametric region, calculated from linear perturbation analysis of the closed-loop system.

This dynamic model can apparently admit multiple equilibrium states. Our equilibrium of interest is the one in which *x*_1_, *z*_1_, and *z*_2_ are all strictly positive. We aim to enforce this desired equilibrium to be a stable fixed point of the closed-loop system. Let us denote it by *q*_*d*_. We shall assume, as a given prerequisite prior to defining a control regularization task, that the set-point to be tracked is chosen such that is admissible by applying the intended control inputs. Thereby, it is reachable starting from some region of attraction in the positive orthant 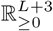, if the closed-loop stability is given.

As proved in Supplementary Note 4, this positive equilibrium is locally (asymptotically) stable provided the only condition 2*γ*_*c*2_ ≤ *γ*_*p*1_ is met. Thus, as long as the equilibrium remains positive, fixing *γ*_*c*2_ small enough ensures the stability of the closed-loop system independently from *Q*_0_ and the exact value of the set-point. From a practical point of view, one may find it interesting when considering bounded parametric uncertainties on the controlled plant. As one can meet the control objective (RPA) and achieve the robust stability by only tuning one single design parameter (here *γ*_*c*2_), while the target set-point could already be freely tweaked by varying other design parameters. Of course, that the admissibility of this reference set-point still needs to be taken into consideration separately. Shaded in light blue, the stability regions are shown in Figure 4B and Figure 5 for a range of uncertainties on the model parameters and where *γ*_*c*2_ is fixed such that *γ*_*c*2_ = *γ*_*p*1_*/*2. As can be seen, increasing *k*^*\**^ alone helps keep maintaining the positiveness of the desired fixed point so long as *r*^*^ is set less than *R*_*tot*_.

### Multicellular Realization of the Layered Autocatalytic Integrator Motif

Inspired by the concept of the layered autocatalytic IFC introduced in Multi-layer Autocatalytic Biomolecular Controllers: An Effective Solution, here we harness some social interactions between two strains of a competitive microbial ecosystem to suggest an experimental realization of the proposed controller’s dynamics, given by (6b)-6c). We treat these two strains, called **N**_1_ and **N**_2_, as two different species, whose viable cell density will be expressed by non-negative values *N*_1_(*t*) and *N*_2_(*t*). We take that the intrinsic growth of each **N**_1_ and **N**_2_ population follows logistic growth [21] in the absence of the other. Let both species be co-cultured in the same medium and be **R** auxotrophs; that is, they must draw from a (limited) resource pool **R** present in the medium—either consuming or temporarily occupying it—to reproduce themselves. Essential for self-reproduction, this resource is a growth-limiting substance, meaning its availability in the growth medium determines, to a large extent, the growth rate of both **N**_1_ and **N**_2_. We mainly consider this shared pool to consist of common nutrients, such as sources of carbon, certain amino acids, or specific fatty acids, present in the medium, but it could also represent other environmental factors— such as energy, light, some gases, or even a shared space—or a combination of them, depending on the context.

The two strains will therefore compete with each other for the *consumption* of **R** to grow, resulting in population-level competitive dynamics between **N**_1_ and **N**_2_. This we will exploit to construct the necessary dynamics realizing (6b)-6c). At the core of the idea, we will basically leverage the fact that the population individuals grow naturally, divide and double regularly, and use it as a means for resembling the autoregulation parts of (6). The growing community is supposed to be in a well-mixed medium, which might undergo continuous dilutions with tunable rates under certain setups. For an experimental setting, one may find the microchemostat setup in [96] or [70] relevant, which enable for long-term monitoring of the synthetic community under study for over hundreds of hours.

In this section, we consider a more general setup where the integral control action, realized by **N**_1_ and **N**_2_, applies to an *n*-species poorly-known process network. Our controller applied to microbial population regulation tasks will be later discussed as a special case study. Note that the nature of considered process species here can be highly heterogeneous, representing population counts, cell types, or concentration of biomolecules, for instance. We generally allow this process to include *m* different populations (0 ≤ *m* ≤ *n*), labeled by **N**_i+2_ and *i* ∈ {1, · · ·, *m*}. The rest of species can be, for example, target proteins being expressed within each or some of the populations **N**_i_, including **N**_1_ and **N**_2_. Note, the populations in the controller side may have indirect interaction and become coupled with those in the process side, for example, through cross-feeding and mutualism, or through competition for the same resource **R**. We will keep the convention of using the italic letters *N*_*i*_(*t*) to denote the respective cell density of population **N**_i_ for every *i*.

The necessary cell-cell communication channels between different populations will be established by means of orthogonal quorum sensing systems [70, 71, 75, 78, 91]. The genes responsible for synthesizing such diffusible signaling molecules could be engineered to activate a signaling pathway, for instance, leading to the upregulation of specific growth factors, whereby accelerating the growth of certain microbial species. Further, the quorum sensing pathways could be engineered to confine cross-feeding of specific metabolites among microbial species. This provides flexibility to synthetically assign varied social roles within an engineered community.

Leveraging this flexibility, we define specific guidelines for intercellular communications through which our controller strains exchange information with the process network to establish sensing and actuation. Similar to the previous sections, we label the output species, whose concentration needs to be regulated, by **X**_n_. It can represent the extracellular concentration of a small signaling molecule secreted into the medium, or it can represent the average copy number of an intracellular species (if in high-copy regimes) within one of the cell populations. In the former case, any individual population or a mix of them within the consortium might act as a sender of **X**_n_. Just like the core motif in Multi-layer Autocatalytic Biomolecular Controllers: An Effective Solution, the internal links between **N**_1_ and **N**_2_—essential for their synergistic collaboration in achieving integral control—naturally arise from resource couplings, eliminating the need for arranging additional communication channels or enforcing social roles for their direct contact.

Even in a nutrient-rich mono-culture setup, with abundant growth factors in the media and no superior competitor to invoke the competitive exclusion principle, the growth of microbial species would still eventually level off when the population reaches the environment’s carrying capacity or when waste products accumulate to inhibitory levels. Various models have been developed to understand the complex dynamics of microbial communities and their emerging behaviors at the ecological level. Let us rely on Lotka-Volterra models to account for the interspecies interactions arising from competition in engineered consortia [70, 91, 97] and to describe the qualitative behavior of our synthetic ecosystem under study. We propose the following closed-loop dynamic model

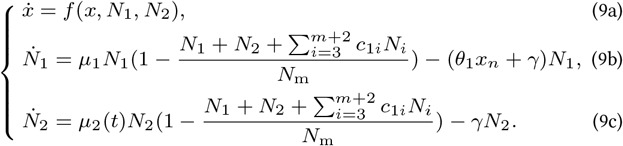

Refer to Figure 6 for a closed-loop circuit illustration. In what follows, we discuss the details and assumptions relevant to this model and present an example (where the function *f* takes a specific form). To enable the realization of the layered autocatalytic strategy, **X**_n_ needs to have two (opposing) interactions with the community species **N**_1_ and **N**_2_. Note that these two interactions may not necessarily be direct, but rather mediated through comparably fast indirect pathways. First, **X**_n_ is required to inhibit the population count of **N**_1_. This can happen, for example, if **X**_n_ induces cell death through a signaling cascade. Second, the intrinsic growth rate of **N**_2_, which we specified in the above model by *µ*_2_(*t*) : ℝ_≥0_ → ℝ_≥0_, should be induceable by **X**_n_: a *growth-stimulating* interaction. As such, in **N**_2_ a similar signaling cascade may express essential genes that increase cellular fitness and, hence, growth.

**Figure 6:**
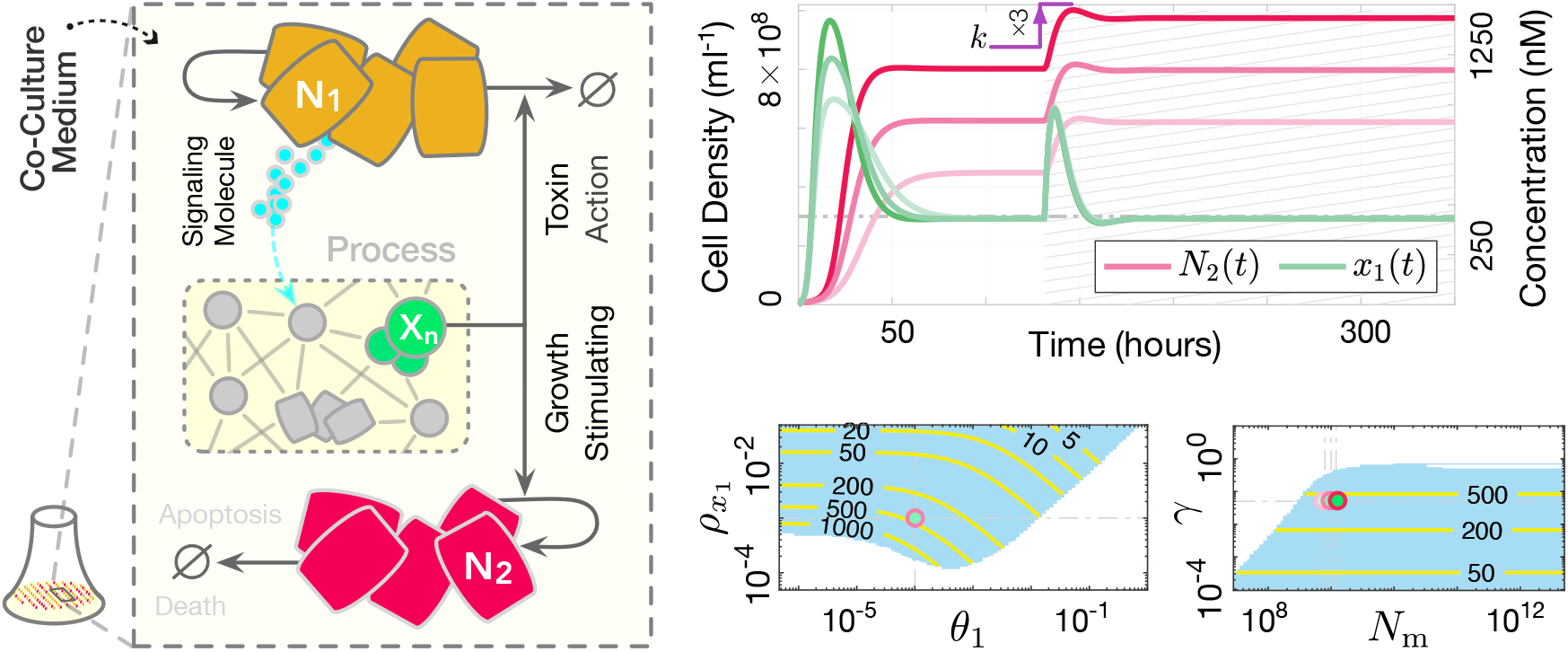
Potential population-level biological realization of the proposed controller. Each species **N**_i_ represents a distinct cell population, such as differentiated cell types from stem cell lines or diverse microbial species in an engineered ecosystem. The controlled process can include populations beyond **N**_1_ and **N**_2_, provided the resulting feedback system does not lead to the exclusion of either **N**_1_ or **N**_2_. Left: Closed-loop circuit illustration of (9) operating in a batch-mode culture setup. The co-culture medium contains shared resources **R**, essential for the growth of both **N**_1_ and **N**_2_ strains, with limited availability leading to direct competition (e.g., common nutrients). The species within the “process” box represent the regulated process network, which can be highly heterogeneous, including intracellular biomolecules, diffusible substances, cell populations, etc. The species **X**_n_, the output of interest to be regulated, exerts opposing effects on the controller populations **N**_1_ and **N**_2_. A dynamic balance between the death rate of **N**_1_ and the doubling rate of **N**_2_ steers the concentration of **X**_n_ toward a predetermined steady-state value, assuming a stabilizing actuation reaction is engineered. The integral feedback is achieved through the participation of both **N**_1_ and **N**_2_ cells at the population level, effectively realizing a multicellular integrator. Right: Sample trajectories for *n* = 1 and *f* := *kN*_1_ − *γ*_*p*1_*x*_1_, along with stability analysis results over uncertain intervals of 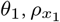, the dilution term *γ*, and the carrying capacity *N*_m_. The (solid, yellow) contour lines in the stable (light-blue) regions show the corresponding steady-state values for *x*_1_(*t*). Numerical values and parameters are provided in Supplementary Note 6.

Here **X**_n_ is seen as a growth inducer for **N**_2_. It can be engineered to, for example, upregulate the synthesis of certain recombinant growth factors. These factors can, in turn, stimulate the proliferation of individual **N**_2_ cells, enhance their nutrient uptake, or promote cell division and metabolic activities—all contributing to an enhanced growth rate for **N**_2_ cell populations. Alternatively, it could be that the induction of **X**_n_ inhibits the production of certain growth inhibitory proteins in **N**_2_, thereby accelerating the growth of **N**_2_ cells through mitigating growth burden. In these general examples, the carrying capacity of the environment for **N**_2_—the maximum number of **N**_2_ it can sustain in isolation—may remain unchanged, as no new metabolite or additional nutrients have been introduced to the growth medium. For simplicity, we model this growth induction using first-order kinetics. It follows 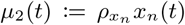 with 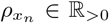 associated reaction rate constant. The implications of using Hill functions instead to address saturation effects on this term can be found in Supplementary Note 5.

As per the works [70, 71, 77, 90, 91, 97, 98], we assume that the populations **N**_1_ and **N**_2_ are two strains of the same microbial species and that, given the chosen resource **R** and despite potential differences in intrinsic growth rates in isolation, the environment’s carrying capacity for both **N**_1_ and **N**_2_ is identical. Thus, the average per capita inhibiting effect that **N**_2_ has on the growth of **N**_1_, and vice versa, is equal to the average self-inhibiting effect that either of populations has on its own growth. Additionally, the per capita inhibition effect of competitors (other populations existing in the co-culture) on **N**_1_ is considered the same as their effect on **N**_2_, and conversely, the per capita effect that **N**_1_ has on the other populations is considered the same as the effect that **N**_2_ has on them. We note that deviating from these assumptions could limit the controller’s capacity to achieve perfect adaptation, potentially resulting in near-perfect adaptation instead.

In the above model, 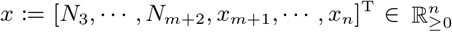 is the augmented state variable representing the process species, 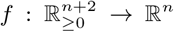, and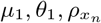, *N*_m_ are positive constants. The dilution rate *γ* can be zero. *N*_m_ is the carrying capacity of mixed microbial populations for **N**_1_ and **N**_2_. Some of the process species in *x* are indeed the process populations *N*_3_ to *N*_*m*+2_, if any. The Lotka-Volterra competition coefficients in (9b)-(9c), denoted by *c*_*ji*_, quantify the average per capita inhibiting effect that the population *N*_*i*_ has on the growth of population *N*_*j*_, relative to the effect that *N*_*j*_ has on its own growth as a result of consuming resources (let every *c*_*jj*_ be set to unity). If no other population relies on the resources **R** for their growth, all the constants *c*_1*i*_ will be zero.

Comparing (9) to (6), this system realizes the proposed motif.

The strictly positive equilibrium entails

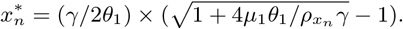

This reference signal (the set-point) is hard-wired into various characteristics of the controller populations—including parameters associated with their growth—which offers additional degrees of freedom and allows for better accommodation in the design of an optimal experimental setup. Closing the loop using an appropriate way of actuating on the plant may ensure the stability of this equilibrium. Assuming that the consortium comprises solely the populations **N**_1_ and **N**_2_, let us briefly examine one sample closed-loop circuit. Let the process network consist of a single species, and let its internal dynamics be described by *f* (*x*_1_, *N*_1_, *N*_2_) := *kN*_1_ − *γ*_*p*1_*x*_1_, with *γ*_*p*1_, *k* positive scalars. Here, **N**_1_ carries the actuation signal to the process network. If the species **X**_1_ is produced as a direct result of the growth and division in **N**_1_, a biological interpretation of it could be diffusible quorum sensing molecules constitutively expressed from some synthase gene in **N**_1_, such as the autoinducer acyl-homoserinelactone (AHL) found in gram-negative bacteria. Alternatively, **X**_1_ could be regarded as a specific gene in **N**_1_ cells that is acted upon by an AHL synthesized from **N**_1_ with fast dynamics. The concentration of the species **X**_1_ will be regulated according to the population-level characteristics of the circuit, despite uncertainties on the competition parameters, *γ*_*p*1_, and *k*. We provide our numerical simulations in Figure 6. Note that, wherever available (including in the remaining sections), the nominal values for parameters are taken close to reported experimental ones [21, 70, 96].

### Population-Level Controller for Microbial Population Control: An Application Example

Here, we turn to an application example of the introduced multi-cellular integrator, where the controller strains aim to regulate the population of another strain. In particular, we consider three competing microbial strains in the same culture. Via secretion of AHL molecules, two of them communicate with and act on the third one (target strain) to enable IFC. The control objective is to maintain the population of the target strain (**N**_3_) at a level proportional to an inducible set-point, robust to its growth profile variations.

Now, let us keep the core parts of the controller represented by (9b)-9c) intact and consider another special form for the process dynamics in (9a). Let us add in another strain to the consortium, say **N**_3_, and take that the plant under control consists only of three species: **N**_3_ and two orthogonal AHL molecules **A**_1_ and **A**_3_. Assume that the individuals of population **N**_3_ also participate in the competition for the common nutrients. Let **N**_3_ (**N**_1_) constitutively secrete the AHL **A**_3_ (**A**_1_) and think of it as the target species the controller aims to regulate (as a growth-stimulating molecule for **N**_3_ by which the controller actuates on the plant). A symbolic illustration of such a three-strain synthetically engineered microbial community can be found in Figure 7, whose governing dynamics may be written as

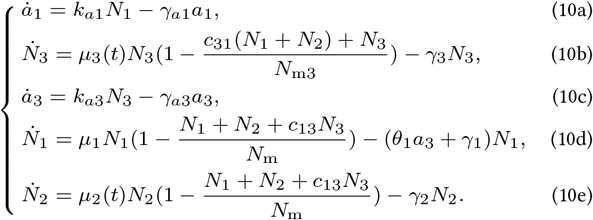

**Figure 7:**
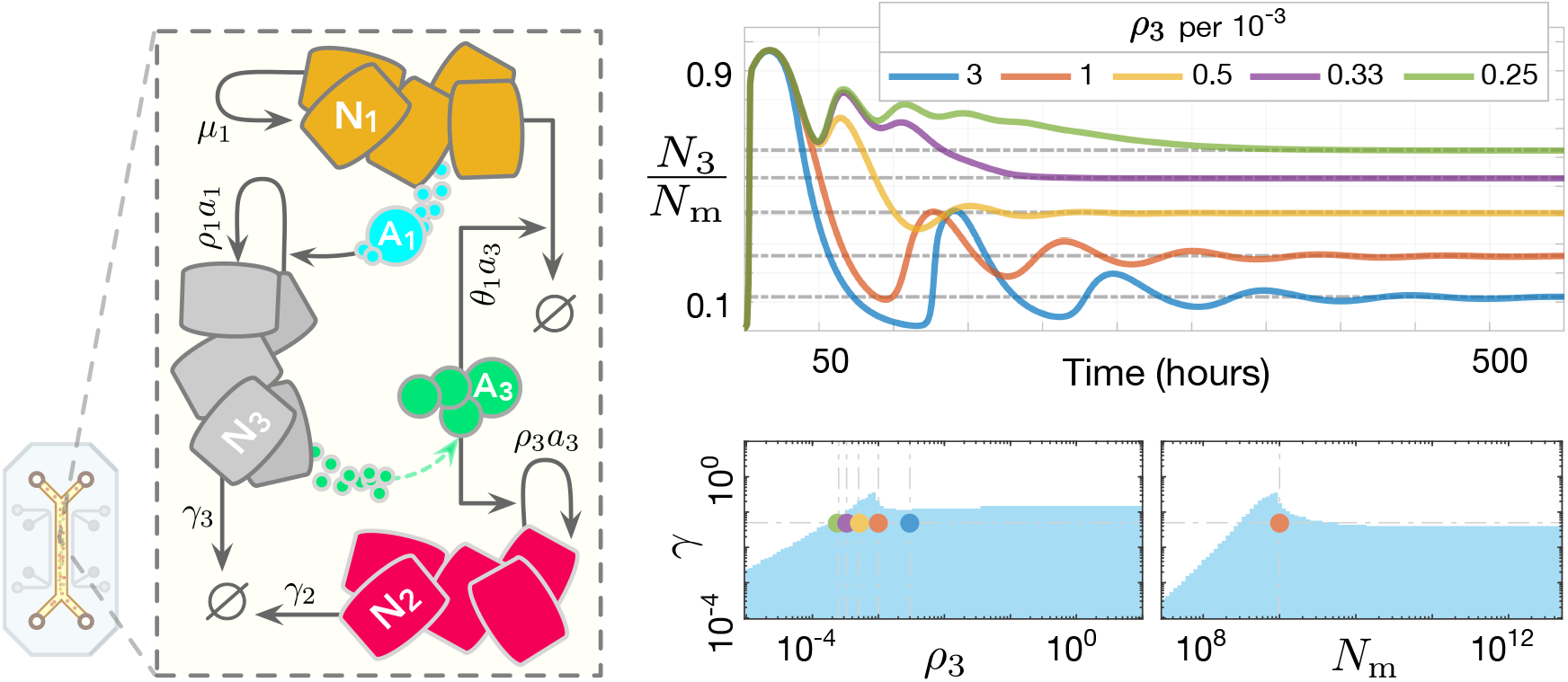
Microbial population control using a layered autocatalytic integral feedback mechanism. The left panel illustrates an example of the general network from Figure 6, where the regulated species, **A**_3_, is a quorum-sensing molecule secreted by another cell population, **N**_3_, in a co-culture system. This system could be established in a microchemostat bioreactor, allowing for long-term observation of the engineered ecosystem. There is natural competition between the two controller strains due to the competitive exclusion principle, with each strain vying for resources and potentially outcompeting the other. The opposing effects of **A**_3_ on **N**_1_ and **N**_2_ exacerbate this issue, as increased **A**_3_ reduces **N**_1_ while promoting **N**_2_, risking the exclusion of **N**_1_. However, the system can be stabilized through properly engineered feedback mechanisms. For example, engineering **N**_1_ to positively influence **A**_3_ by growth-stimulating interactions with **N**_3_ could mitigate the initial surge in **A**_3_, stabilizing the ecosystem. The numerical results support feedback stabilization across various set-point values, assessing the stability of the closed-loop circuit (10) under bounded parametric uncertainties. The right panel shows the temporal evolution of normalized **N**_3_ for different set-points. See Supplementary Note 6 for parameter values.

The carrying capacity of the co-culture system for the species **N**_3_ is denoted by *N*_m3_, which might in general differ from that for populations **N**_1_ and **N**_2_. In line with works such as [16, 21, 70, 90, 91, 96], we model the dynamics of the AHL-mediated quorum sensing systems considered in this article, whose autoinducers are assumed to diffuse freely across the plasma membrane and, as a result of it, have uniform concentrations both inside and outside the cells, by first-order reactions as in (10a) or (10c). Simply assuming that the changes on the concentration of an autoinducer in the extracellular volume, which serves as a signal indicator for all the populations sensing it, can be approximated by a birth-death process whose death rate is fixed while its birth rate is linearly dependent on the density of the source cells producing the autoinducer.

Similar to the general control setup modeled in (9), we take again mass-action kinetics with first-order linear dynamics to model the growth-stimulating interactions. Hence, for the intrinsic growth rates of the cells **N**_2_ and **N**_3_ we have *µ*_2_(*t*) := *ρ*_3_*a*_3_ and *µ*_3_(*t*) := *ρ*_1_*a*_1_. The desired equilibrium here follows

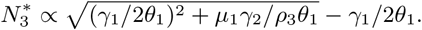

Observe that this steady state is not sensitive to **N**_3_’s parameters in (10b), nor is it sensitive to the competition constants *c*_13_, *c*_31_, and *N*_m3_. Thus, as long as the coexistence of species at stationary phase is maintained—the closed-loop stability is preserved—one would expect that the precise population control of strain **N**_3_ will be achieved regardless of tolerable disturbances on it. Such disturbances can arise from various sources, including antibiotic-induced depletion, the upregulation of toxin genes, or shifts in environmental conditions, for example. For a set of parameters where the competition coefficients are set to unity, *N*_m3_ = *N*_m_, and where all the dilution terms *γ*_*i*_s are set identical to the value *γ*, numerical simulations presented in Figure 7 assess the stability of the closed-loop system (10) to give some insight on the parameter ranges in which such a circuit is functional. We observe that the introduced feedback system effectively regulates the density of the targeted strain, ensuring stable co-existence of the microbial species within the consortium across a comparatively wide range of set-points (induced by variations in *ρ*_3_ and *γ*) and in the presence of parametric uncertainties.

### Ratiometric Control using a Layered Autocatalytic Integral Feedback Strategy

Let us formally define a ratiometric regulator problem as one involving the design of a controller to manage a system with two outputs subject to an unknown disturbance input, given a commanded reference signal. The controller’s goal is to act on the controlled plant in a way that the relative values between the two outputs of interest are steered towards this (constant) reference signal. The controller aims to maintain the desired ratio between the two outputs, regardless of their absolute magnitudes or initial values, particularly when disturbances are present.

In this section, we modify the core controller motif introduced in Multi-layer Autocatalytic Biomolecular Controllers: An Effective Solution to adapt it for resource-aware ratiometric control tasks. The primary control objective is to regulate the ratio between two distinct species of the process at steady state. These two process species we shall denote by **X**_1_ and **X**_2_. We define 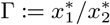 to capture the output of interest to be regulated. Here, 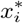 represents the concentration of species **X**_i_ at steady state, where the transient response is sufficiently damped to be considered negligible. We will think of Γ as an implicit input signal to the controller, which determines the reference set-point. The goal is to apply a stabilizing control input to the process network aimed at controlling the ratio level Γ arbitrary close to a predetermined, adjustable value, throughout an extended period of observation. This is to be achieved despite potential persistent (step-like) disturbances affecting process parameters and structure, or (sudden) global changes in cellular resource availability triggered by external loads or downstream demands.

Reported in Supplementary Note 5, one possible control law capable of achieving such a control objective can be biochemically realized by the controller motif represented in Figure 8A. The associated closed-loop dynamic model may be written as follows

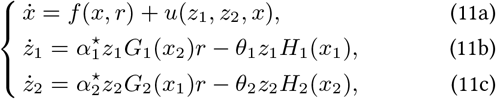

where the nonlinear maps *H*_*i*_ and *G*_*i*_ are to capture possible saturation effects. We assume negligible constant controller degradation/dilution. The function *u* in (11a) is defined to represent the stabilizing control input(s), which shall be regarded not as given but rather as a design problem. As with the previous sections, the way the controller acts on the plant is case-specific, to be determined by the designer. See Supplementary Note 5 for details.

**Figure 8:**
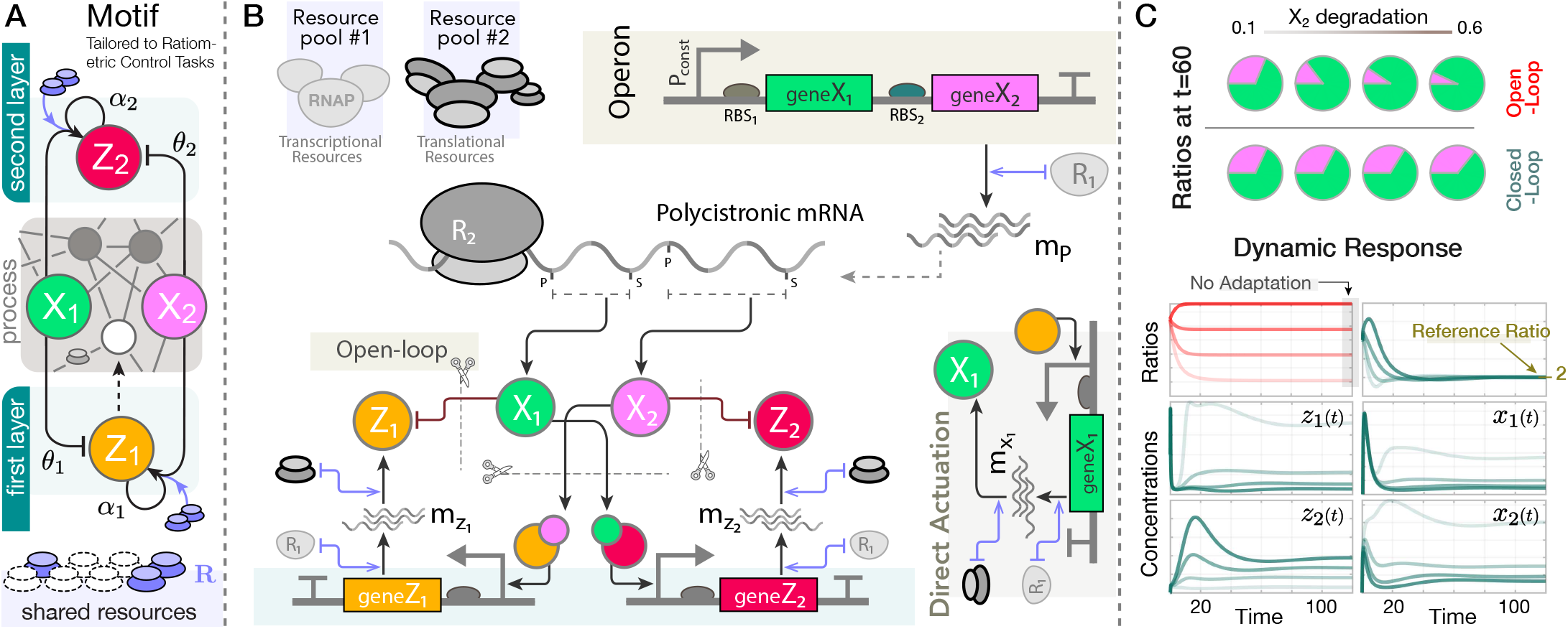
Ratiometric control using a layered autocatalytic integral feedback strategy. (**A**) illustrates the modified motif, where the controller now senses two process species, **X**_1_ and **X**_2_. The control objective is to steer the concentration ratio of these species to a prescribed reference, Γ. (**B**) Embedded ratio control in prokaryotes: Schematic of the circuit discussed in Robust Gene Expression Ratio Control in a Resource-Limited Setting. Refer to (S94) in Supplementary Note 6 for the closed-loop mathematical model. The system co-expresses two genes as its open-loop circuit, with two additional genes driving the controller, all within the same host cell. The operon gene is constitutively expressed. Indirect couplings due to competition for transcriptional and translational resources are managed by two separate resource pools, **R**_1_ and **R**_2_. There is direct actuation from **Z**_1_ to **X**_1_. (**C**) Time response trajectories of the circuit for four different values of 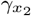(a.u.). The computed ratios, *x*_1_(*t*)*/x*_2_(*t*), from the open-loop circuit (red lines) are not robust against parameter variations, while the controlled plant (teal lines) shows robustness. In the open-loop setting, direct transcriptional actuation on **X**_1_ was treated as a zeroth-order activation input, allowing fine-tuning of the ratio. Structural disruptions in (**A**) or significant controller degradation/dilution may lead to *imperfect* ratiometric control (see Supplementary Note 5). Numerical values and further details are available in Supplementary Note 6.

For any strictly positive steady-state response of this closed-loop system (if it exists), the following relation between steady states of the species **X**_1_ and **X**_2_ holds true

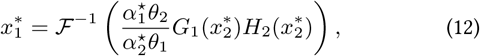

where ℱ (*ρ*) := *G*_2_(*ρ*)*H*_1_(*ρ*) and ℱ^−1^ refers to its inverse function, which exists and is well-defined on the positive axis. Note that the relation expressed by equation (12) actually maps the two steady states to each other through a smooth manifold (to be referred to as the *ratio manifold*), which is uniquely identifiable solely based on the controller’s parameters and structure. This mapping is generally nonlinear, but becomes a line, thereby enabling *perfect* ratiometric control, if we consider linear forms for all functions *H*_*i*_ and *G*_*i*_. By solving from (12) for the cases of purely linear mappings with *G*_*i*_(*ρ*) := *ρ* and *H*_*i*_(*ρ*) := *ρ*, the ratio Γ can be easily obtained as

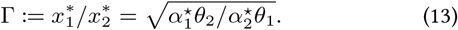

Assuming the robust stability of the closed-loop system given (bounded) uncertainties on the process network, this result signifies the achievement of *ratiometric adaptation* perfectly and robustly in response to stimuli affecting the controlled plant. Furthermore, in this case, the controller automatically manages for the parallel regulation of **R**, with the steady-state expression for it given by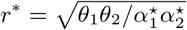.

Note that the expression for Γ will change if either of the assumptions—no saturation or no controller dilution—is violated. Deviating from either assumption, however, merely shifts the mapping from 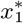 to 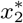 from a purely linear to a nonlinear subspace. Nonetheless, the relationship between the two steady states will be still determined exclusively by the design parameters and remains independent of the process under control, as well as the disturbances that do not affect (12) and the dilution terms. This implies an *imperfect* ratiometric control, emphasizing the still existence of a dynamic controller effort aimed at compensating for ratio changes induced by disturbances, even if factors such as controller saturation and dilution were to be taken into account. The impact of saturation and constant controller degradation/dilution on the ratio manifold is further discussed in Supplementary Note 5. It is worth mentioning that the same argument applies to the concentration regulation problems discussed in previous sections, as one can directly relate the steady-state ratio results of this section to those scenarios by simply considering the control input *x*_2_(*t*) as constant.

Of note, the long-term concentrations of the species **X**_1_ and **X**_2_ do not necessarily need to approach an isolated single *point* in space to maintain the controller dynamics at zero. In fact, the absorbing attractor of the system after transient phase could be limit cycles or other (non-isolated) periodic orbits, while the ratio between **X**_1_ and **X**_2_ counts still remains tightly regulated. Within the scope of this article, however, we primarily focus on point attractors for the closed-loop system. In the remainder, we will explore two different application examples.

### Robust Gene Expression Ratio Control in a Resource-Limited Setting

In the first application example, we address the problem of regulating gene expression ratios between two distinct genes co-expressed within the same host. Envision these genes being incorporated into an operon for simultaneous transcription, a typical experimental approach for controlling expression ratios in prokaryotes despite being an open-loop control strategy. In this scenario, we will have the following production reactions

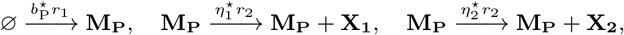

which are assumed to be resource limited, where the species **R**_1_ and **R**_2_ are to represent the available transcriptional and translational resources, respectively. Here, the species **M**_P_, **X**_1_, and **X**_2_ represent the resulting polysictronic mRNA strand and the two proteins of interest, respectively. Every species **X** is subject to constant degradation with the corresponding rate *γ*_*x*_. Note that the effect of transcriptional competition on the steady-state ratio 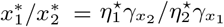 is eliminated thanks to the fact that the mRNA numbers are exactly the same for both of the genes. This might help with a better regulation of the ratios between **X**_1_ and **X**_2_, in a broad sense, but it remains vulnerable to variations in the parameters 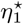 and 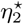, as well as uncertainties in the degradation rates 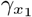 and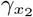. The parameters 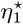 and 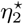, representing the expression rates of **X**_1_ and **X**_2_ from the mRNA strand, respectively, depend on the choice of the ribosome binding sites (RBS) associated to their corresponding genes. Also, the degradation of proteins could be modulated by synthetically introducing degradation tags and use of constitutive proteases, for instance. For fixed choices of RBS_1_ and RBS_2_, this operon-based approach to control the ratio between **X**_1_ and **X**_2_ usually requires fine-tuning of the degradation rates, which is not an efficient way of encoding a reference ratio. It thereby demands for an alternative strategy, such as the use of negative feedback and closed-loop control, for a better performance.

In light of this, we integrate our controller into the circuit described above, which we shall treat as the regulated process network. This enables for a robust ratio control scheme that is resilient with regard to unknown factors and disturbances which concern **M**_P_, **X**_1_, or **X**_2_. A diagram of the resulting closed-loop circuit is depicted in Figure 8B, wherein the controller species are assumed to be participating in the competition for both transcriptional and translational resources. In contrast to the previous sections that focused on a single limiting resource pool, we chose a more detailed model to investigate the controller’s performance in the presence of multiple limiting cellular resources. The closed-loop dynamic model is given in Supplementary Note 6. It is noted therein how solving for the ratio 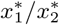 maps to the formula given by (13). According to this note, the controller holds 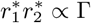 at steady state. This implies that the robustness brought about by the negative feedback ensues from the controller’s effort to establish and maintain a relationship between the product of the two available resources (**R**_1_ and **R**_2_) and the reference ratio (Γ), even in the presence of disturbances.

Simulation results are included in Figure 8C to support the findings. Compared to an open-loop control approach, which suffers from lack of robustness, these results highlight our controller’s ability to manage intracellular ratios robust, to a large extent, against undesired factors and disturbances. These include variations in the expression levels of the two controlled genes at the translation stage or differences in their degradation rates. As examples, the former can be a cause of using ribosome binding sites of varying strengths, while the latter may result from using different proteins for **X**_1_ and **X**_2_ with varying relative thermodynamic stability, or from the fusion of degradation domains (degrons) to either protein.

### Co-culture Composition Control in an Engineered Multi-Strain Microbial Consortium

The application example we consider here adopts a multicellular approach to implement the layered autocatalytic controller in (11) and demonstrates its application in the dynamic co-culture control of a multi-strain engineered community. We build this case study upon the population control circuit (10) and expand it to incorporate the additional components necessary for resembling the dynamics of (11). Let us begin by adding a new microbial species to the consortium, which relies on the same limiting substance **R** to grow and self-reproduce. Consider it as the species **X**_2_ treated in (11) and refer to it by **N**_4_. The goal is to achieve robust tuning of the cell density ratio levels between the two populations **N**_3_ and **N**_4_ when grown together, irrespective of their own growth profiles. We further introduce two new orthogonal signaling molecules, **A**_2_ and **A**_4_, and assume them to be (constitutively) secreted from **N**_2_ and **N**_4_, respectively. With these eight species in the reaction network, we wire the population-level interactions between them—either of growth-stimulating or toxin types of interactions—according to the circuit sketched in the pictorial representation Figure 9.

**Figure 9:**
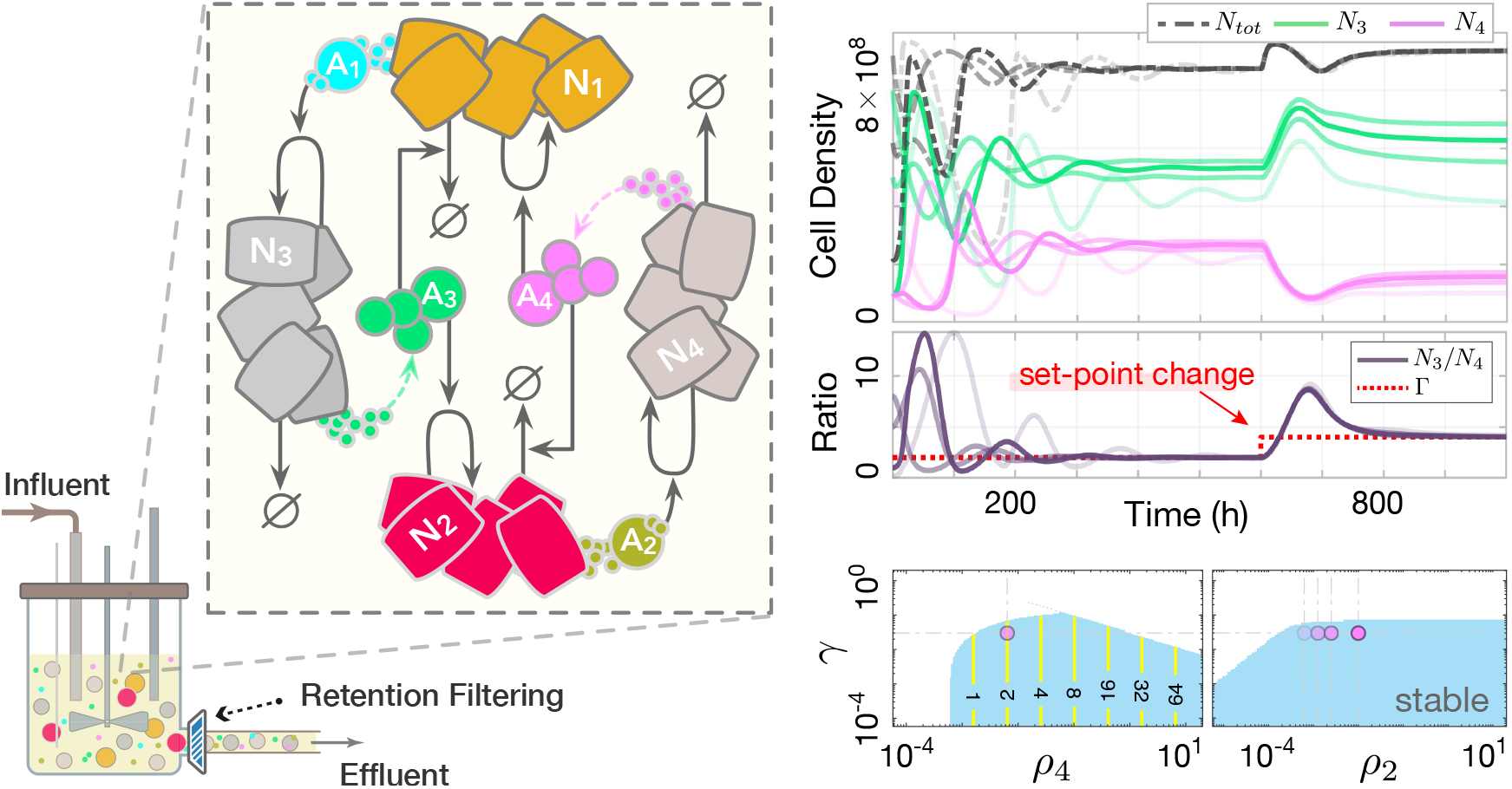
Multicellular implementation of a layered autocatalytic IFC mechanism enables community composition control in an engineered microbial consortium. Left: Circuit schematic of the example from Co-culture Composition Control in an Engineered Multi-Strain Microbial Consortium. See (S95) in Supplementary Note 6 for the closed-loop dynamic model. The bioreactor cartoon suggests a continuous-culture setup for co-culturing the cells in the proposed circuit. This bioreactor setup, especially with a retentive filter to prevent washout of controller cells, can achieve perfect ratiometric control. Right: Set-point tracking response for four different initial conditions and parameters. In each case, the process parameter *ρ*_2_ and initial composition are varied. The total population (dash-dotted lines), 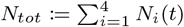, represents the sum of all cell densities. At time *t* = 600 h, *ρ*_4_ is suddenly increased fourfold, changing the reference ratio Γ from 2 to 4. Populations **N**_3_ and **N**_4_ adjust their levels to match the new Γ. The evolution of the cell density ratio *N*_3_(*t*)*/N*_4_(*t*) is shown in a separate graph. The bottom plots present numerical results from local stability assessments, considering a wide range of uncertainties in process parameters *γ* and *ρ*_2_, and the controller parameter *ρ*_4_. Variations in *ρ*_4_ shift the set-point Γ, indicated by the yellow contour lines. Numerical values and parameters are detailed in Supplementary Note 6.

Consistent with the previous sections, we keep using *µ*_*i*_ to describe the intrinsic growth rate of the microbial species **N**_i_. Let us confine the way the controller strains act on the process (actuation) to population-level interactions of growth-stimulating types such that *µ*_*i*_ = *ρ*_*i*−2_*a*_*i*−2_ for *i* ∈ {3, 4}. For the sensing reactions, introduce *µ*_1_ = *ρ*_4_*a*_4_ and *µ*_2_ = *ρ*_3_*a*_3_. A model for this closed-loop control system is given in Supplementary Note 6. For a set of parameters, we also specify therein the sufficient conditions for the admissibility of a given reference ratio. This control system holds

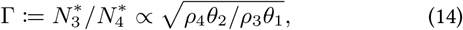

if the stability of its unique, (strictly) positive equilibrium is given. Thanks to the several independent parameters encoding the commanded ratio, there is enough flexibility to make any given reference ratio Γ admissible: by choosing sufficiently large values for *ρ*_3_ or *ρ*_4_, and a slow enough rate of dilution. See the inequalities (S96).

Note, our proposed population-level ratiometric regulator offers a universal solution for controlling the ratios between two, even entirely different, microbial species of interest, such as one being *E. coli* and the other *B. subtilis*. But, this holds as long as the introduced interspecies interactions lead to a stabilized community dynamics; in other words, none of the four microbial species present in the consortium go extinct over long periods of observation. Thus, we consider bounded uncertainties on the model parameters to numerically examine the robustness of the circuit when the growth profiles of the controlled species vary significantly. The results are provided in Figure 9, which assess the dynamic response and stability of the considered multicellular circuit and identify a working range of achievable strain ratios for a set of nominal parameters.

In our model, the process populations are subject to constant dilutions at a rate of *γ*, whereas the controller strains are not. This is to enable *perfect* composition control, deviating from which may result instead in *imperfect* ratiometric adaptation (as discussed in Ratiometric Control using a Layered Autocatalytic Integral Feedback Strategy and Supplementary Note 5). As far as developing a feasible experimental setup appropriate for the resulting circuit above is concerned, multiple scenarios warrant consideration. One such scenario involves cultivating the strains in a fed-batch mode while incorporating constitutive killer genes into both **N**_3_ and **N**_4_ cells to account for the constant degradation rate, *γ*. Alternatively, if the controller cells are larger in size than the process cells, one can still choose continuous culture and resort to a chemostat setting with a dilution rate of *γ*, provided that a selective retention device is employed that guarantees the retention of controller cells. This could be done by use of specific permeable membranes, or, potentially, some capillary fibers, that function in a way that allows the **N**_3_ and **N**_4_ cells to cross over and become diluted, while retaining the microbial species **N**_1_ and **N**_2_ within the growth medium. Alternatively, it could be through employing special cross-flow filtration mechanisms that recirculate the larger cells back to the medium while subjecting the process cells to the desired dilution rate, similar to those used in long-term perfusion cell cultures.

## Discussion

Biomolecular integral feedback controllers play an important role in synthetic biology, as they could be leveraged to regulate various vital biological processes. Several controller motifs have been discovered that can realize integral feedback and ensure robust perfect adaptation (RPA). Such RPA-achieving controllers can find widespread applications within synthetic biology such as population control and ratio control. A subset of these employs positive autoregulation of controller species to establish integral action, which can be found favorable in certain design cases since self-replication and self-regulatory networks are ubiquitous in nature across scales. Concerning precise gene regulation within rapidly growing organisms like *E. coli*, the inherent robustness of autocatalytic IFCs against exponential cell growth and constant dilution factors—compared to non-autocatalytic controllers [29–31]—highlights their potential for achieving better performance at the molecular and cellular scales.

However, resource competition presents a significant challenge to these autocatalytic controllers, as it can impact their ability to maintain RPA. This limitation restricts the use of autocatalytic control loops in developing novel synthetic circuit designs, particularly those where positive autoregulation can simplify implementation. This encompasses the biocircuit designs that aim to leverage multi-cellular interacting systems, division of labor, or a combination of intracellular and interspecific social interactions between different populations in order to introduce various innovative functionalities. To address this challenge, we initially developed a mathematical framework that accounts for the effect of competition for scarce, shared resources. Biological systems are known for their complexity, characterized by intricate interactions within a high-dimensional network of species. This complexity underscores the importance of developing computational and analytical approaches that are capable of accounting for them. Our resource-aware framework lays the foundational building blocks for modeling resource competition within an intricate web of intracellular reactions, extending to the bimolecular levels. Besides its simplicity in use, the introduced framework is versatile and expandable, enabling more detailed modeling of resource-limited chemical reaction networks (CRNs).

Various bimolecular scenarios were investigated, along with expansions of the framework. However, the scope of cases examined may not yet cover all possible scenarios of intracellular resource-limited reactions. We leave the door open for future developments to include more scenarios. Efforts in this direction, potentially building upon the core reactions considered in this study, could arm the framework to encompass, for example, reactions of higher dimensions—including those limited by three or more shared resources—or other bimolecular scenarios involving as intermediary steps some intramolecular reactions or irreversible cleavages.

Next, we specifically demonstrated that the minimal realization of the autocatalytic integral feedback control (IFC) scheme fails to achieve RPA in the presence of resource competition. Resource competing scenarios offer a more realistic context to analyze the performance of such controllers claiming superlinear growth rates, given that the mere existence of a self-reproducing species often falls short of guaranteeing its effective replication. Typically, successful self-reproduction hinges on the availability of certain resources, which introduce self-inhibition if limited and dynamically couple competing entities sharing them. Motivated to resolve the observed limitation, we introduced a multi-layer autocatalytic control strategy. This approach, named layered autocatalytic IFC, successfully restored the RPA property amidst competition for shared resources. The restoration was achieved by introducing another autocatalytic feedback layer, which draws from the same resource pool as the original loop. The newly introduced species functions as an adaptive resource allocator. It buffers available resources, releasing them as needed later on. In a case study, we showcased the effectiveness of our controller in maintaining RPA during embedded gene-expression control tasks, despite translational resource competition.

More broadly, the coupling between the controller species does not have to represent common resources; it could be any rate-limiting shared inhibitor. Competitive resource couplings might just simply be naturally emerging instances. The fact that our proposed biocontroller does not rely on the exact dynamics of resource couplings to establish IFC makes it more versatile and applicable to environments involving different types of resource competition and across different scales. Leveraging this, we pursued the implementation of our control approach using one of perhaps the most prevalent types of autocatalytic production in nature, that is, the replication of chromosomal DNA, which typically manifests through population dynamics as a result of the cell cycle and cell division. We considered population dynamics emerging from a microbial consortium and the ecological interactions exhibited by its members. Indeed, whenever that population-level dynamics are meant to be used for realizing a synthetic controller, one would need to deal with the emergence of competitive autocatalytic terms. This is due to the intrinsic nature of dividing cells that consume nutrients, often shared with co-existing rivals, to self-reproduce. We utilized generalized Lotka-Volterra models to capture potential competitive interactions within the engineered consortia studied in our work. These models have been demonstrated through several experimental studies to successfully capture the dynamics of microbial competition, both in engineered consortia [70] and in the human gut microbiota [99].

A multicellular realization of our controller motif was proposed and investigated in the presence of competition for limited resources, e.g., common nutrients in a synthetically engineered consortium. The integrator achieves the integral action by actually establishing a dynamic balance between the death and doubling rates of certain co-cultured community species. Here, two strains of the same microbial species jointly create IFC loops at the population level when co-cultured together. In particular, we examined the performance of the resulting controller in a population control task. An attractive feature for biotechnological applications is that the system has to be characterized and optimized only once for controlling the strain of interest. The optimized controller parameters do not require further tuning even if the target strain or its growth profile change afterwards, provided the stability of the closed-loop circuit and, therefore, the long-term stable coexistence of the controller microbial species is preserved.

The introduced motif exemplifies one of the simplest architectures to achieving integral feedback control at the multicellular scale, where the integral action is cooperatively brought about by the contributions from at least two different cell populations. The outcome of the contest between these two microbial species, fueled by both their competition for resources and their engineered interactions with the process species, ultimately contributes to the robust regulation of output levels. Essentially, in this setup, it is the individual cells and their natural reproductive systems that serve as agents executing the controller functions. Given a nutrient-rich environment, these systems are naturally optimized to function effectively at higher densities, if necessary. This might be seen as an advantage when compared to embedded genetic implementations, offering a potential alternative that mitigates the genetic burden and slowed growth often associated with the overexpression of controller genes in embedded settings.

Lastly, we demonstrated how the introduced layered autocatalytic IFC mechanism can be adapted to address ratiometric control tasks. Given two distinct process species, a ratiometric control problem was defined to regulate the steady-state concentration ratio between them. We introduced the modifications necessary to enable the controller for this purpose. The efficacy of the resulting control mechanism was explored through two application examples. First, we incorporated our controller embedded within the same cell as the process to enable robust ratio control between two co-expressed genes. Second, we built upon our core population-level controller and introduced the required cell-cell communications through quorum sensing molecules to enable robust tuning of the co-culture composition, that is, controlling the cell density ratio levels between two other populations when grown together with the controller strains, irrespective of their own growth profiles.

Representing the signaling molecules of our circuits through general yet abstract notation as CRN species provides greater design flexibility. This allows them to represent quorum sensing, analytes within a microenvironment, chemotactic cytokines, or even morphogenes in a developmental process. Further research to expand this versatility might reveal a range of other adaptations of the multicellular biocontrollers discussed in this work, customized for specific applications in fields such as cell fate control, synthetic developmental biology, and bacteria-based cancer immunotherapy. By potentially moving beyond ecological models to incorporate genome-scale mechanistic models—aimed at finely integrating the dynamic coupling between intracellular genetic factors, growth, and cellular metabolism—we believe that such explorations into our multi-layer biocontrollers could set the stage for their promising practical applications in biotechnology and bioprocesses.

## Acknowledgements

The authors would like to thank Dr. Stephanie Aoki, Dr. Maurice Filo, and Dr. Joaquin Gutierrez Mena for their insightful discussions, as well as Dr. Maurice Filo for their valuable feedback on the manuscript and proofreading it.

