## Supplementary Material for "Multi-Layer Autocatalytic Feedback Enables Integral Control Amidst Resource Competition and Across Scales"

**This supplementary document contains:**

- Supplementary text (including Supplementary Notes 1 to 6)
- Figures S1 to S9
- Tables S1 to S3

---

### Outline

In this Supplementary Material of the manuscript titled "*Multi-Layer Autocatalytic Feedback Enables Integral Control Amidst Resource Competition and Across Scales*," we first provide in Section S1 the extended derivations of the mathematical framework introduced for modeling intracellular resource competition. More in the same section, we explore two other bimolecular cases of resource-limited production reactions (the fifth and sixth scenarios) and then discuss how to extrapolate the framework to accommodate different other types of resource-limited reactions. In Section S2, we present several relevant biological examples regarding the framework, aimed at providing further insights into how to conceptually incorporate the introduced framework into the modeling of resource-limited biological processes. The supplementary notes follow, concluding this document.

### S1 Extended Derivations of the Resource-Competition Modeling Framework

In this section, we provide the derivation of the framework introduced in A Mathematical Framework to Model Intracellular Resource Competition of the main manuscript. We do so after giving an overview of the reaction network representations for each type of considered resource-limited reaction. This overview includes a detailed treatment of the interactions between shared resources and substrates, covering every involved elementary reaction. We initially focus on a given chemical reaction network (CRN) composed exclusively of resource-limited catalytic production reactions, in which resource-limited bimolecular reactions of first four scenarios are included. We further cover two other scenarios of bimolecular catalytic production reactions in Section S1.3. The remaining subsections will consider extensions of the framework to different other scenarios of intracellular resource competition, including resource-limited conversion reactions, degradation/sequestration reactions, and reactions limited by the availability of two different resource pools.

#### S1.1 Reaction Network Representations

For the resource-limited zeroth-order (catalytic) production reaction $\mathcal{R}_i^z$, we consider that it follows the set of reactions

$$\mathbf{R} \underset{k_{r_i}^z}{\overset{k_{f_i}^z \bar{s}_i^z}{\rightleftharpoons}} \mathbf{C_i^z} \xrightarrow{k_{c_i}^z} \mathbf{R} + \mathbf{P_i^z}, \tag{S1}$$

where $\mathbf{C_i^z}$ is an intermediate complex formed at a rate proportional to the available free resources, $\mathbf{P_i^z}$ is the product species, and where the inflow rate $\bar{s}_i^z \in \mathbb{R}_{>0}$ is assumed to remain constant over time. This type of resource-limited reaction might be useful, for example, when explicitly modeling for the expression of housekeeping genes or when considering the genetic burden imposed by the synthesis of a constituitively-expressed gene on that of a target synthetic gene. An illustration of this network of reactions can be found in Figure 1.

We model each resource-limited unimolecular catalytic production reaction $\mathcal{R}_i^u$ as incorporating a substrate, referred to as $\mathbf{S_i^u}$, that binds to the resource $\mathbf{R}$ and forms an intermediate complex $\mathbf{C_i^u}$. This (active) complex can then be converted into a product $\mathbf{P_i^u}$ while releasing the resource $\mathbf{R}$ and the substrate $\mathbf{S_i^u}$ as well. In chemical reaction notation, we write

$$\mathbf{S_i^u} + \mathbf{R} \underset{k_{r_i}^u}{\overset{k_{f_i}^u}{\rightleftharpoons}} \mathbf{C_i^u} \xrightarrow{k_{c_i}^u} \mathbf{S_i^u} + \mathbf{R} + \mathbf{P_i^u}. \tag{S2}$$

For a resource-limited bimolecular catalytic production reaction $\mathcal{R}_i^b$, where three species $\mathbf{A_i}$, $\mathbf{B_i}$, and $\mathbf{R}$ limit the production of $\mathbf{P_i^b}$ simultaneously, we shall assume that it takes place according to one of the following four different scenarios, denoted respectively by S1-S4,

$$\text{S1} \begin{cases} \mathbf{A_i} + \mathbf{B_i} \underset{\sigma_i}{\overset{\eta_i}{\rightleftharpoons}} \mathbf{S_i^b}, \\ \mathbf{S_i^b} + \mathbf{R} \underset{k_{r_i}^b}{\overset{k_{f_i}^b}{\rightleftharpoons}} \mathbf{C_i^b} \xrightarrow{k_{c_i}^b} \mathbf{S_i^b} + \mathbf{R} + \mathbf{P_i^b}, \end{cases} \tag{S3}$$

$$\text{S2} \begin{cases} \mathbf{A_i} + \mathbf{R} \underset{\sigma_i}{\overset{\eta_i}{\rightleftharpoons}} \mathbf{S_i^b}, \\ \mathbf{S_i^b} + \mathbf{B_i} \underset{k_{r_i}^b}{\overset{k_{f_i}^b}{\rightleftharpoons}} \mathbf{C_i^b} \xrightarrow{k_{c_i}^b} \mathbf{S_i^b} + \mathbf{B_i} + \mathbf{P_i^b}, \end{cases} \tag{S4}$$

$$\text{S3}\begin{cases} \mathbf{A_i}+\mathbf{B_i} \underset{\sigma_i}{\overset{\eta_i}{\rightleftharpoons}} \mathbf{S_i^b}, \\ \mathbf{S_i^b}+\mathbf{R} \underset{k^b_{r_i}}{\overset{k^b_{f_i}}{\rightleftharpoons}} \mathbf{C_i^b} \xrightarrow{k^b_{c_i}} \mathbf{A_i}+\mathbf{B_i}+\mathbf{R}+\mathbf{P_i^b}, \end{cases} \tag{S5}$$

and

$$\text{S4}\begin{cases} \mathbf{A_i}+\mathbf{R} \underset{\sigma_i}{\overset{\eta_i}{\rightleftharpoons}} \mathbf{S_i^b}, \\ \mathbf{S_i^b}+\mathbf{B_i} \underset{k^b_{r_i}}{\overset{k^b_{f_i}}{\rightleftharpoons}} \mathbf{C_i^b} \xrightarrow{k^b_{c_i}} \mathbf{A_i}+\mathbf{B_i}+\mathbf{R}+\mathbf{P_i^b}. \end{cases} \tag{S6}$$

Observe that in all these four scenarios, the reactions occur in two similar steps as depicted schematically in Figure 1. In the first step, two of the three limiting species bind to each other and form an intermediate complex $\mathbf{S_i^b}$. In the second step, once the intermediary $\mathbf{S_i^b}$ has been formed, the remaining species binds to $\mathbf{S_i^b}$ to form the resource-substrate complex $\mathbf{C_i^b}$. This ternary complex then releases the product $\mathbf{P_i^b}$ and the other reactants. The intermediate complex $\mathbf{S_i^b}$ can be released as a whole or as its individual constituents.

Observe from the representations (1)-(3) in A Mathematical Framework to Model Intracellular Resource Competition that the smooth function $v_i^o$, with $o \in \{z, u, b\}$, was defined to only capture the effect of a single reaction $\mathcal{R}_i^o$ on the rate at which $\mathbf{P_i^{\mathit{o}}}$ is being produced. Of course, the production/degradation propensities of $\mathbf{P_i^{\mathit{o}}}$ undergo changes according to many other interactions that the product $\mathbf{P_i^{\mathit{o}}}$ might have with the rest of the network, too. It is implicitly assumed that the species $\mathbf{S_i^u}$, $\mathbf{A_i}$, $\mathbf{B_i}$, $\mathbf{P_i^z}$, $\mathbf{P_i^u}$, and $\mathbf{P_i^b}$ are representing known, but not necessarily unique, species of the considered CRN. In fact, without loss of generality, $\mathbf{S_i^u}$ and $\mathbf{P_i^u}$ (or $\mathbf{A_i}$, $\mathbf{B_i}$, and $\mathbf{P_i^b}$ in the bimolecular case) may represent the same biochemical species and the product can be one or multiple species, including the reactant species $\mathbf{S_i^u}$, $\mathbf{A_i}$, and $\mathbf{B_i}$ themselves. In the following subsections, we attempt to derive (approximate) formulae for describing each production rate $v_i^z$, $v_i^u$, and $v_i^b$ based solely on the concentrations of the substrates $\mathbf{S_i^u}$s, $\mathbf{A_i}$s, or $\mathbf{B_i}$s that all compete for $\mathbf{R}$.

#### S1.2 Main Derivation

Considering mass-action kinetics, the dynamic model of the reaction set (S1), representing a resource-limited zeroth-order (catalytic) production reaction $\mathcal{R}_i^z$, follows

$$\begin{cases} \dot{c}_i^z = k^z_{f_i}\bar{s}_i^z r - (k^z_{r_i}+k^z_{c_i})c_i^z, & \text{(S7a)} \\ \dot{p}_i^z = k^z_{c_i}c_i^z =: v_i^z, & \text{(S7b)} \end{cases}$$

in which we left out writing the dynamics of the state variable $r$. Later, we will compute $r$ through a conservation law and based on imposing quasi-steady state assumptions on some other state variables. Consistent with the manuscript, lower-case letters represent the concentration of the associated species denoted by bold, capital letters. Similarly, for a resource-limited unimolecular catalytic production reaction $\mathcal{R}_i^u$ we have that

$$\begin{cases} \dot{s}_i^u = -k^u_{f_i}s_i^u r + (k^u_{r_i}+k^u_{c_i})c_i^u, & \text{(S8a)} \\ \dot{c}_i^u = k^u_{f_i}s_i^u r - (k^u_{r_i}+k^u_{c_i})c_i^u, & \text{(S8b)} \\ \dot{p}_i^u = k^u_{c_i}c_i^u =: v_i^u. & \text{(S8c)} \end{cases}$$

Now, we consider the dynamic model of the resource-limited bimolecular catalytic production reactions, restricting our consideration to only the first four possible scenarios. Consistent with the notations in Section S1.1 and that of Figure 1, we continue indicating these scenarios by the labels "S1" to "S4", respectively. So, we consider that the dynamic model of the chemical reactions describing $\mathcal{R}_i^b$ corresponds to one of the following scenarios. For $\mathcal{R}_i^b$ being of the first scenario, we have

$$\begin{cases} \dot{a}_i = -\eta_i a_i b_i + \sigma_i s_i^b, & \text{(S9a)} \\ \dot{b}_i = -\eta_i a_i b_i + \sigma_i s_i^b, & \text{(S9b)} \\ \dot{s}_i^b = \eta_i a_i b_i - \sigma_i s_i^b - k_{f_i}^b s_i^b r + (k_{r_i}^b + k_{c_i}^b) c_i^b, & \text{(S9c)} \\ \dot{c}_i^b = k_{f_i}^b s_i^b r - (k_{r_i}^b + k_{c_i}^b) c_i^b, & \text{(S9d)} \\ \dot{p}_i^b = k_{c_i}^b c_i^b. & \text{(S9e)} \end{cases}$$

If $\mathcal{R}_i^b$ belongs to the second scenario

$$\begin{cases} \dot{a}_i = -\eta_i a_i r + \sigma_i s_i^b, & \text{(S10a)} \\ \dot{b}_i = -k_{f_i}^b s_i^b b_i + (k_{r_i}^b + k_{c_i}^b) c_i^b, & \text{(S10b)} \\ \dot{s}_i^b = \eta_i a_i r - \sigma_i s_i^b - k_{f_i}^b s_i^b b_i + (k_{r_i}^b + k_{c_i}^b) c_i^b, & \text{(S10c)} \\ \dot{c}_i^b = k_{f_i}^b s_i^b b_i - (k_{r_i}^b + k_{c_i}^b) c_i^b, & \text{(S10d)} \\ \dot{p}_i^b = k_{c_i}^b c_i^b. & \text{(S10e)} \end{cases}$$

For $\mathcal{R}_i^b$ following the third scenario, it reads

$$\begin{cases} \dot{a}_i = -\eta_i a_i b_i + \sigma_i s_i^b + k_{c_i}^b c_i^b, & \text{(S11a)} \\ \dot{b}_i = -\eta_i a_i b_i + \sigma_i s_i^b + k_{c_i}^b c_i^b, & \text{(S11b)} \\ \dot{s}_i^b = \eta_i a_i b_i - \sigma_i s_i^b - k_{f_i}^b s_i^b r + k_{r_i}^b c_i^b, & \text{(S11c)} \\ \dot{c}_i^b = k_{f_i}^b s_i^b r - (k_{r_i}^b + k_{c_i}^b) c_i^b, & \text{(S11d)} \\ \dot{p}_i^b = k_{c_i}^b c_i^b. & \text{(S11e)} \end{cases}$$

Finally, if it is associated with the fourth scenario we get

$$\begin{cases} \dot{a}_i = -\eta_i a_i r + \sigma_i s_i^b + k_{c_i}^b c_i^b, & \text{(S12a)} \\ \dot{b}_i = -k_{f_i}^b s_i^b b_i + (k_{r_i}^b + k_{c_i}^b) c_i^b, & \text{(S12b)} \\ \dot{s}_i^b = \eta_i a_i r - \sigma_i s_i^b - k_{f_i}^b s_i^b b_i + k_{r_i}^b c_i^b, & \text{(S12c)} \\ \dot{c}_i^b = k_{f_i}^b s_i^b b_i - (k_{r_i}^b + k_{c_i}^b) c_i^b, & \text{(S12d)} \\ \dot{p}_i^b = k_{c_i}^b c_i^b. & \text{(S12e)} \end{cases}$$

Let us define $v_i^b$ to be $v_i^b := k_{c_i}^b c_i^b$ in all the above-mentioned four scenarios.

To ease referring to multiple bimolecular scenarios and ease compact representation of them in single equations, let us assign to each reaction $\mathcal{R}_i^b$ of scenario S$j$ (with $j \in \{1, 2, 3, 4\}$) a new parameter called $\zeta_i$ defined by $\zeta_i := j$. Let the indicator function

$\mathbb{1}_j : \mathbb{R} \to \{0,1\}$ with $j$ an integer be defined such that $\mathbb{1}_j(x) = 1$ whenever $x = j$ and $\mathbb{1}_j(x) = 0$ otherwise (note that, for any $x$, $\mathbb{1}_k(x)\mathbb{1}_l(x) = \mathbb{1}_k(x)$ if $k = l$ and is zero if $k \neq l$). With a deterministic point of view and on average levels, we can then write down the following mass balance equality

$$r(t) = R_{\text{const}} - \sum_{i=1}^{q_0} c_i^z(t) - \sum_{i=1}^{q_1} c_i^u(t) - \sum_{i=1}^{q_2} [c_i^b(t) + (\mathbb{1}_2(\zeta_i) + \mathbb{1}_4(\zeta_i)) s_i^b(t)], \tag{S13}$$

where $R_{\text{const}} \in \mathbb{R}_{>0}$ may be considered the total capacity of the resource pool. We assume implicitly that $R_{\text{const}}$ remains unchanged over the time interval of interest, thereby treating it as an unknown yet fixed constant. Speaking of generality, however, this total capacity may be taken as a variable instead, for instance, if there are sources producing or consuming the resources $\mathbf{R}$ with varying rates and within the considered time interval of interest.

Under the assumption A1 made in the main manuscript, in A Mathematical Framework to Model Intracellular Resource Competition, we carry out the following quasi-steady state approximations

$$c_i^z \approx \frac{k_{f_i}^z}{k_{r_i}^z + k_{c_i}^z} \bar{s}_i^z r, \tag{S14}$$

$$c_i^u \approx \frac{k_{f_i}^u}{k_{r_i}^u + k_{c_i}^u} s_i^u r, \tag{S15}$$

$$c_i^b \approx (\mathbb{1}_1(\zeta_i) + \mathbb{1}_3(\zeta_i)) \frac{k_{f_i}^b}{k_{r_i}^b + k_{c_i}^b} s_i^b r + (\mathbb{1}_2(\zeta_i) + \mathbb{1}_4(\zeta_i)) \frac{k_{f_i}^b}{k_{r_i}^b + k_{c_i}^b} s_i^b b_i. \tag{S16}$$

In fact, we have assumed that for every $i$ the resource-binding complexes $\mathbf{C_i^z}$, $\mathbf{C_i^u}$, and $\mathbf{C_i^b}$ together with the intermediate complexes $\mathbf{S_i^b}$ are all at dynamic equilibrium. This is often a fair assumption to make in an intracellular genetic system, noting that the reactions forming $\mathbf{C_i^z}$, $\mathbf{C_i^u}$, $\mathbf{S_i^b}$, and $\mathbf{C_i^b}$ are assumed to be instances of binding/unbinding reactions, which often occur on a much faster time scale than the time scale at which the concentrations of the substrates $\mathbf{S_i^u}$s, $\mathbf{A_i}$s, or $\mathbf{B_i}$s evolve.

Also, for the intermediate complexes $\mathbf{S_i^b}$ we get

$$s_i^b \approx \mathbb{1}_1(\zeta_i) \frac{\eta_i a_i b_i}{\sigma_i} + \mathbb{1}_2(\zeta_i) \frac{\eta_i a_i r}{\sigma_i} + \mathbb{1}_3(\zeta_i) \frac{\eta_i a_i b_i}{\sigma_i + \frac{k_{c_i}^b k_{f_i}^b}{k_{r_i}^b + k_{c_i}^b} r} + \mathbb{1}_4(\zeta_i) \frac{\eta_i a_i r}{\sigma_i + \frac{k_{c_i}^b k_{f_i}^b}{k_{r_i}^b + k_{c_i}^b} b_i}. \tag{S17}$$

On substituting (S14)-(S17) into the equation (S13), we can obtain the following quasi-steady state approximation for $r(t)$ which depends on the concentrations of all network species competing for $\mathbf{R}$

$$r(t) \approx \frac{R_{\text{const}}}{1 + \sum_{i=1}^{q_0} \frac{\bar{s}_i^z}{k_{m_i}^z} + \sum_{i=1}^{q_1} \frac{s_i^u}{k_{m_i}^u} + \sum_{i=1}^{q_2} f^b(a_i, b_i, r, \zeta_i)}, \tag{S18}$$

where

$$f^b := (\mathbb{1}_1(\zeta_i) + \mathbb{1}_2(\zeta_i)) \frac{a_i b_i}{k_{m_i}^b} + \zeta_i^a a_i + \frac{\mathbb{1}_3(\zeta_i) a_i b_i}{k_{m_i}^b (1 + \frac{k_{c_i}^b k_{f_i}^b}{\sigma_i (k_{r_i}^b + k_{c_i}^b)} r)} + \frac{\mathbb{1}_4(\zeta_i) a_i b_i}{k_{m_i}^b (1 + \frac{k_{c_i}^b k_{f_i}^b}{\sigma_i (k_{r_i}^b + k_{c_i}^b)} b_i)}, \tag{S19}$$

$$\zeta_i^a := \mathbb{1}_2(\zeta_i) \frac{\eta_i}{\sigma_i} + \mathbb{1}_4(\zeta_i) \frac{\eta_i}{\sigma_i + \frac{k_{c_i}^b k_{f_i}^b}{k_{r_i}^b + k_{c_i}^b} b_i},$$

$$k^z_{m_i} := (k^z_{r_i} + k^z_{c_i})/k^z_{f_i},$$

$$k^u_{m_i} := (k^u_{r_i} + k^u_{c_i})/k^u_{f_i},$$

and

$$k^b_{m_i} := (k^b_{r_i} + k^b_{c_i})\sigma_i/k^b_{f_i}\eta_i.$$

We shall adopt the approximations given in (S14)-(S17) and (S18) throughout. By noting that the product rates $v^o_i$ are equivalent to $v^o_i = k^o_{c_i}c^o_i$ (where $o \in \{z, u, b\}$), we have already characterized the resource-limited production rates.

Under the assumptions A2-A3 considered in the main manuscript, we can rearrange the above findings as follows

$$\begin{aligned} v^z_i &= \beta^z_i r & &\text{with } \beta^z_i = k^z_{c_i}\bar{s}^z_i/k^z_{m_i}, \\ v^u_i &= \beta^u_i s^u_i r & &\text{with } \beta^u_i = k^u_{c_i}/k^u_{m_i}, \\ v^b_i &= \beta^b_i a_i b_i r & &\text{with } \beta^b_i = k^b_{c_i}/k^b_{m_i}, \end{aligned} \tag{S20}$$

where

$$r = \frac{R_{\text{const}}(1+Q_0)^{-1}}{1 + \sum_{i=1}^{q_1} \frac{(1+Q_0)^{-1}}{k^u_{m_i}} s^u_i(t) + \sum_{i=1}^{q_2} \frac{(1+Q_0)^{-1}}{k^b_{m_i}} a_i(t)b_i(t) + \frac{\zeta^a_i}{1+Q_0}a_i(t)}. \tag{S21}$$

Here, $Q_0 := \sum_{i=1}^{q_0} \bar{s}^z_i/k^z_{m_i}$ and the lumped parameters $\zeta^a_i$ reduce to $\zeta^a_i = (\mathbb{1}_2(\zeta_i) + \mathbb{1}_4(\zeta_i))\eta_i/\sigma_i$. In the main text, we defined

$$w^u_i := k^u_{m_i}/(1+Q_0),$$

$$w^{ab}_i := k^b_{m_i}/(1+Q_0),$$

$$w^a_i := \zeta^a_i/(1+Q_0).$$

The aggregate parameters $k^o_{m_i}$ are given by $k^z_{m_i} := (k^z_{r_i} + k^z_{c_i})/k^z_{f_i}$, $k^u_{m_i} := (k^u_{r_i} + k^u_{c_i})/k^u_{f_i}$, and $k^b_{m_i} := (k^b_{r_i} + k^b_{c_i})\sigma_i/k^b_{f_i}\eta_i$. If the two assumptions A2-A3 are not always fulfilled, one would instead have to directly solve for $r(t)$ for every $t$ from the formula given by the equations (S18) and (S19). It is easy to see that the equation (S18), whose right-hand side is a bounded non-decreasing function of $r$ for $r \in [0, \infty)$, always gives rise to a unique solution for $r(t)$ provided that none of the reverse binding rates $\sigma_i$ belonging to the third scenarios are zero.

#### S1.3 Two Additional Bimolecular Cases (Fifth and Sixth Scenarios)

We now extend our results to include two additional bimolecular cases that may be plausible in certain situations. In these cases, one of the substrates, $\mathbf{A_i}$ or $\mathbf{B_i}$, is recycled in a reversible manner before the complete release of the product and resources. This process leads to the formation of a new species from the residual species, called $\mathbf{Q_i}$. This new intermediate species is assumed to be distinct from the complexes $\mathbf{C^b_i}$ or $\mathbf{S^b_i}$, either in terms of their constituents or their structural configuration. We consider two different scenarios for this to happen, referred to as the fifth and sixth scenarios (also indicated by the labels S5 and S6,

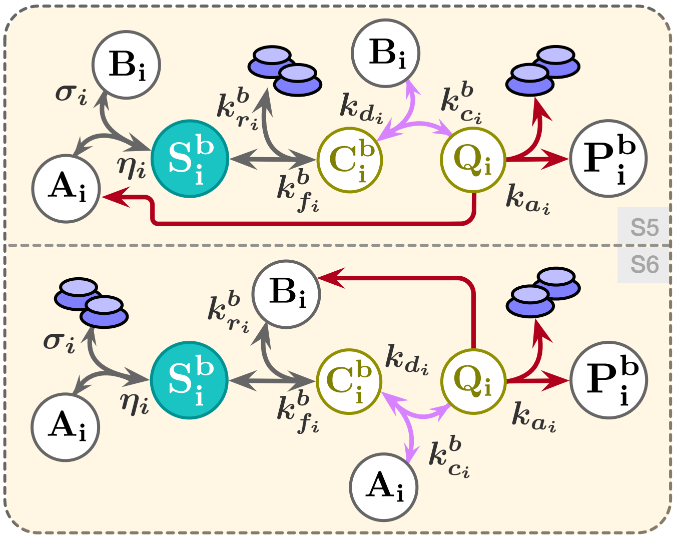

Figure S1: **The two additional scenarios of resource-limited bimolecular reactions.** These scenarios, referred to as the fifth and sixth scenarios of bimolecular interactions involving shared resources, are discussed separately and integrated into the framework in Section S1.3.

respectively). In doing so, we introduce the new species $\mathbf{Q_i}$ and two new reaction rate constants, $k_{d_i}$ and $k_{a_i}$, for each reaction $\mathcal{R}_i^b$. Note that the introduced species and rate constants are exclusive to reactions of the fifth and sixth scenarios; if $\mathcal{R}_i^b$ for the chosen $i$ does not belong to either of these two scenarios, then $\mathbf{Q_i}$, $k_{a_i}$, and $k_{d_i}$ do not exist. In chemical reaction forms, we represent a resource-limited bimolecular reaction $\mathcal{R}_i^b$ of these two scenarios as

$$\text{S5}\begin{cases} \mathbf{A_i}+\mathbf{B_i} \underset{\sigma_i}{\overset{\eta_i}{\rightleftharpoons}} \mathbf{S_i^b}, \\ \mathbf{S_i^b}+\mathbf{R} \underset{k^b_{r_i}}{\overset{k^b_{f_i}}{\rightleftharpoons}} \mathbf{C_i^b} \underset{k_{d_i}}{\overset{k^b_{c_i}}{\rightleftharpoons}} \mathbf{B_i}+\mathbf{Q_i}, \\ \mathbf{Q_i} \xrightarrow{k_{a_i}} \mathbf{A_i}+\mathbf{R}+\mathbf{P_i^b}, \end{cases} \tag{S22}$$

and

$$\text{S6}\begin{cases} \mathbf{A_i}+\mathbf{R} \underset{\sigma_i}{\overset{\eta_i}{\rightleftharpoons}} \mathbf{S_i^b}, \\ \mathbf{S_i^b}+\mathbf{B_i} \underset{k^b_{r_i}}{\overset{k^b_{f_i}}{\rightleftharpoons}} \mathbf{C_i^b} \underset{k_{d_i}}{\overset{k^b_{c_i}}{\rightleftharpoons}} \mathbf{A_i}+\mathbf{Q_i}, \\ \mathbf{Q_i} \xrightarrow{k_{a_i}} \mathbf{B_i}+\mathbf{R}+\mathbf{P_i^b}, \end{cases} \tag{S23}$$

respectively. Schematic drawings of them are provided in Figure S1. Considering mass-action kinetics, the governing dynamical behavior of (S22) and (S23) can be described as what follows. If $\mathcal{R}_i^b$ is associated with the fifth scenario, then

$$\begin{cases} \dot{a}_i = -\eta_i a_i b_i + \sigma_i s_i^b + k_{a_i} q_i, & \text{(S24a)} \\ \dot{b}_i = -\eta_i a_i b_i + \sigma_i s_i^b + k^b_{c_i} c_i^b - k_{d_i} b_i q_i, & \text{(S24b)} \\ \dot{s}_i^b = \eta_i a_i b_i - \sigma_i s_i^b - k^b_{f_i} s_i^b r + k^b_{r_i} c_i^b, & \text{(S24c)} \\ \dot{c}_i^b = k^b_{f_i} s_i^b r - (k^b_{r_i} + k^b_{c_i}) c_i^b + k_{d_i} b_i q_i, & \text{(S24d)} \\ \dot{q}_i = k^b_{c_i} c_i^b - k_{d_i} b_i q_i - k_{a_i} q_i, & \text{(S24e)} \\ \dot{p}_i^b = k_{a_i} q_i. & \text{(S24f)} \end{cases}$$

For $\mathcal{R}_i^b$ belonging to the sixth scenario, we write

$$\begin{cases} \dot{a}_i = -\eta_i a_i r + \sigma_i s_i^b + k_{c_i}^b c_i^b - k_{d_i} a_i q_i, & \text{(S25a)} \\ \dot{b}_i = -k_{f_i}^b s_i^b b_i + k_{r_i}^b c_i^b + k_{a_i} q_i, & \text{(S25b)} \\ \dot{s}_i^b = \eta_i a_i r - \sigma_i s_i^b - k_{f_i}^b s_i^b b_i + k_{r_i}^b c_i^b, & \text{(S25c)} \\ \dot{c}_i^b = k_{f_i}^b s_i^b b_i - (k_{r_i}^b + k_{c_i}^b) c_i^b + k_{d_i} a_i q_i, & \text{(S25d)} \\ \dot{q}_i = k_{c_i}^b c_i^b - k_{d_i} a_i q_i - k_{a_i} q_i, & \text{(S25e)} \\ \dot{p}_i^b = k_{a_i} q_i. & \text{(S25f)} \end{cases}$$

For each reaction $\mathcal{R}_i^b$, we assign the auxiliary parameter $\zeta_i = 5$ if it belongs to the fifth scenario, or $\zeta_i = 6$ if it belongs to the sixth scenario, respectively. We redefine $v_i^b$ as $v_i^b := (\mathbb{1}_1(\zeta_i) + \mathbb{1}_2(\zeta_i) + \mathbb{1}_3(\zeta_i) + \mathbb{1}_4(\zeta_i)) k_{c_i}^b c_i^b + (\mathbb{1}_5(\zeta_i) + \mathbb{1}_6(\zeta_i)) k_{a_i} q_i$.

Let us update the conservation law for $r(t)$ as follows

$$\begin{aligned} r(t) = R_{\text{const}} &- \sum_{i=1}^{q_0} c_i^z(t) - \sum_{i=1}^{q_1} c_i^u(t) \\ &- \sum_{i=1}^{q_2} [c_i^b(t) + (\mathbb{1}_2(\zeta_i) + \mathbb{1}_4(\zeta_i) + \mathbb{1}_6(\zeta_i)) s_i^b(t) + (\mathbb{1}_5(\zeta_i) + \mathbb{1}_6(\zeta_i)) q_i(t)]. \end{aligned} \tag{S26}$$

We revisit the assumption A1 to include also the new intermediate species $\mathbf{Q_i}$.

**Assumptions.** *The following condition holds for every considered resource-limited reaction:*

*A1 For every $i$, the resource-binding complexes $\mathbf{C_i^z}$, $\mathbf{C_i^u}$, and $\mathbf{C_i^b}$ together with the intermediate complexes $\mathbf{S_i^b}$ and $\mathbf{Q_i}$ (if exists) are all at dynamic equilibrium.*

Based on this main assumption, we set the dynamics of the resource-binding and intermediate complexes to zero and calculate their representative quasi-steady state approximations. Meanwhile, we will refer to the previously obtained results for the zeroth, unimolecular, and bimolecular reactions (up to fourth scenario) as needed. For $\mathbf{Q_i}$ at equilibrium, we find

$$q_i \approx k_{c_i}^b c_i^b \left( \frac{\mathbb{1}_5(\zeta_i)}{k_{a_i} + k_{d_i} b_i} + \frac{\mathbb{1}_6(\zeta_i)}{k_{a_i} + k_{d_i} a_i} \right). \tag{S27}$$

Also, for the resource-binding complexes $\mathbf{C_i^b}$ we obtain

$$\begin{aligned} c_i^b \approx (\mathbb{1}_1(\zeta_i) + \mathbb{1}_3(\zeta_i)) \frac{k_{f_i}^b}{k_{r_i}^b + k_{c_i}^b} s_i^b r + (\mathbb{1}_2(\zeta_i) + \mathbb{1}_4(\zeta_i)) \frac{k_{f_i}^b}{k_{r_i}^b + k_{c_i}^b} s_i^b b_i \\ + \mathbb{1}_5(\zeta_i) \frac{k_{f_i}^b}{k_{r_i}^b + \dfrac{k_{c_i}^b k_{a_i}}{k_{a_i} + k_{d_i} b_i}} s_i^b r \\ + \mathbb{1}_6(\zeta_i) \frac{k_{f_i}^b}{k_{r_i}^b + \dfrac{k_{c_i}^b k_{a_i}}{k_{a_i} + k_{d_i} a_i}} s_i^b b_i. \end{aligned} \tag{S28}$$

Using (S27) and (S28), we arrive at

$$s_i^b \approx \mathbb{1}_1(\zeta_i)\frac{\eta_i a_i b_i}{\sigma_i} + \mathbb{1}_2(\zeta_i)\frac{\eta_i a_i r}{\sigma_i} + \mathbb{1}_3(\zeta_i)\frac{\eta_i a_i b_i}{\sigma_i + \dfrac{k^b_{c_i}k^b_{f_i}}{k^b_{r_i}+k^b_{c_i}}r} + \mathbb{1}_4(\zeta_i)\frac{\eta_i a_i r}{\sigma_i + \dfrac{k^b_{c_i}k^b_{f_i}}{k^b_{r_i}+k^b_{c_i}}b_i}$$
$$+ \mathbb{1}_5(\zeta_i)\frac{\eta_i a_i b_i}{\sigma_i + \dfrac{k^b_{c_i}k^b_{f_i}}{k^b_{r_i}+k^b_{c_i}+\dfrac{k_{d_i}}{k_{a_i}}b_i}r}$$
$$+ \mathbb{1}_6(\zeta_i)\frac{\eta_i a_i r}{\sigma_i + \dfrac{k^b_{c_i}k^b_{f_i}}{k^b_{r_i}+k^b_{c_i}+\dfrac{k_{d_i}}{k_{a_i}}a_i}b_i}. \tag{S29}$$

Substituting (S27)-(S29) into (S26) reads the following quasi-steady state approximation for $\mathbf{R}$

$$r(t) \approx \frac{R_{\text{const}}}{1 + \sum_{i=1}^{q_0}\dfrac{\bar{s}_i^z}{k^z_{m_i}} + \sum_{i=1}^{q_1}\dfrac{s_i^u}{k^u_{m_i}} + \sum_{i=1}^{q_2} f^b(a_i, b_i, r, \zeta_i)}, \tag{S30}$$

where

$$f^b := (\mathbb{1}_1(\zeta_i)+\mathbb{1}_2(\zeta_i))\frac{a_i b_i}{k^b_{m_i}} + \zeta_i^a a_i + \frac{\mathbb{1}_3(\zeta_i)a_i b_i}{k^b_{m_i}(1+\dfrac{k^b_{c_i}k^b_{f_i}}{\sigma_i(k^b_{r_i}+k^b_{c_i})}r)} + \frac{\mathbb{1}_4(\zeta_i)a_i b_i}{k^b_{m_i}(1+\dfrac{k^b_{c_i}k^b_{f_i}}{\sigma_i(k^b_{r_i}+k^b_{c_i})}b_i)}$$
$$+ \mathbb{1}_5(\zeta_i)\frac{\eta_i a_i b_i}{\sigma_i + \dfrac{k^b_{c_i}k^b_{f_i}}{k^b_{r_i}+k^b_{c_i}+\dfrac{k_{d_i}}{k_{a_i}}b_i}r} \times \frac{k^b_{f_i}(1+k^b_{c_i}/(k_{a_i}+k_{d_i}b_i))}{k^b_{r_i}+\dfrac{k^b_{c_i}k_{a_i}}{k_{a_i}+k_{d_i}b_i}}$$
$$+ \mathbb{1}_6(\zeta_i)\frac{\eta_i a_i b_i}{\sigma_i + \dfrac{k^b_{c_i}k^b_{f_i}}{k^b_{r_i}+k^b_{c_i}+\dfrac{k_{d_i}}{k_{a_i}}a_i}b_i} \times \frac{k^b_{f_i}(1+k^b_{c_i}/(k_{a_i}+k_{d_i}a_i))}{k^b_{r_i}+\dfrac{k^b_{c_i}k_{a_i}}{k_{a_i}+k_{d_i}a_i}}. \tag{S31}$$

Where, also,

$$\zeta_i^a := \mathbb{1}_2(\zeta_i)\frac{\eta_i}{\sigma_i} + \mathbb{1}_4(\zeta_i)\frac{\eta_i}{\sigma_i + \dfrac{k^b_{c_i}k^b_{f_i}}{k^b_{r_i}+k^b_{c_i}}b_i} + \mathbb{1}_6(\zeta_i)\frac{\eta_i}{\sigma_i + \dfrac{k^b_{c_i}k^b_{f_i}}{k^b_{r_i}+k^b_{c_i}+\dfrac{k_{d_i}}{k_{a_i}}a_i}b_i}.$$

Again, we shall adopt the approximations given in (S27)-(S29) and (S30) throughout. It is straightforward to verify that solving for $r(t)$ from equation (S30), while keeping every other variable fixed, always yields a unique positive solution if all $\sigma_i$s from the third and fifth scenarios are non-zero. This is because all $r$-dependent terms in (S31) are bounded non-increasing functions of $r$ over the interval $r \in [0, \infty)$. In addition to the assumptions A1-A3, let us also impose the two simplifying assumptions below.

**Assumptions.** *In the considered CRN, the following two conditions hold true for every resource-limited bimolecular reaction associated with the fifth and sixth scenarios:*

*A4 Each bimolecular reaction* $\mathcal{R}_i^b$ *of fifth scenario, following the representation given in (S22), satisfies* $\sigma_i \gg k^b_{c_i}k^b_{f_i}r(t)/(k^b_{r_i} +$

$k^b_{c_i} + \frac{k_{d_i}}{k_{a_i}} b_i(t))$.

*A5 Each bimolecular reaction $\mathcal{R}^b_i$ of sixth scenario, following the representation given in (S23), satisfies $\sigma_i \gg k^b_{c_i} k^b_{f_i} b_i(t)/(k^b_{r_i} + k^b_{c_i} + \frac{k_{d_i}}{k_{a_i}} a_i(t))$.*

Given the assumptions A1-A5, one can recast the obtained results as

$$
\begin{aligned}
v^z_i &= \beta^z_i r && \text{with } \beta^z_i = k^z_{c_i} \bar{s}^z_i / k^z_{m_i}, \\
v^u_i &= \beta^u_i s^u_i r && \text{with } \beta^u_i = k^u_{c_i} / k^u_{m_i}, \\
v^b_i &= \beta^b_i a_i b_i r && \text{with } \beta^b_i = \sum_{m=1}^{4} \mathbb{1}_m(\zeta_i) \frac{k^b_{c_i}}{k^b_{m_i}} + \frac{\mathbb{1}_5(\zeta_i) k^b_{f_i} \eta_i / \sigma_i}{1 + \frac{k^b_{r_i}}{k^b_{c_i}}(1 + \frac{k_{d_i}}{k_{a_i}} b_i)} + \frac{\mathbb{1}_6(\zeta_i) k^b_{f_i} \eta_i / \sigma_i}{1 + \frac{k^b_{r_i}}{k^b_{c_i}}(1 + \frac{k_{d_i}}{k_{a_i}} a_i)},
\end{aligned}
\tag{S32}
$$

where

$$
r = \frac{R_{\text{const}}(1 + Q_0)^{-1}}{1 + \sum_{i=1}^{q_1} \frac{(1 + Q_0)^{-1}}{k^u_{m_i}} s^u_i(t) + \sum_{i=1}^{q_2} \frac{\sum_{m=1}^{4} \mathbb{1}_m(\zeta_i)}{k^b_{m_i}(1 + Q_0)} a_i(t) b_i(t) + \frac{\zeta^a_i}{1 + Q_0} a_i(t) + \frac{g^b_i(a_i, b_i)}{1 + Q_0} a_i(t) b_i(t)},
\tag{S33}
$$

$g^b_i : \mathbb{R}^2_{\geq 0} \to \mathbb{R}_{\geq 0}$ is a function defined by

$$
g^b_i(a_i, b_i) := \frac{\eta_i}{\sigma_i} [\mathbb{1}_5(\zeta_i) \frac{k^b_{f_i}(1 + k^b_{c_i}/(k_{a_i} + k_{d_i} b_i))}{k^b_{r_i} + \frac{k^b_{c_i} k_{a_i}}{k_{a_i} + k_{d_i} b_i}} + \mathbb{1}_6(\zeta_i) \frac{k^b_{f_i}(1 + k^b_{c_i}/(k_{a_i} + k_{d_i} a_i))}{k^b_{r_i} + \frac{k^b_{c_i} k_{a_i}}{k_{a_i} + k_{d_i} a_i}}],
$$

and $\zeta^a_i$ reads $\zeta^a_i = (\mathbb{1}_2(\zeta_i) + \mathbb{1}_4(\zeta_i) + \mathbb{1}_6(\zeta_i))\eta_i/\sigma_i$. Note that $\beta^o_i$ are no longer all constants in terms of binding affinities involved in the constituting elementary reactions. Rather in the fifth (sixth) scenario, $\beta^b_i$ is a function of the substrate species $\mathbf{B_i}$ ($\mathbf{A_i}$).

#### S1.4 Considerations for Intracellular Resource-Limited Conversion Reactions

The catalytic production reactions considered above rely on both resources and reactants acting as enzymatic catalysts. The occurrence of conversion reactions can also be limited by the availability of resources. In contrast, we refer to such reactions, in which one or more of the reactants are gradually depleted over time, as resource-limited conversion reactions. The catalytic productions through the reactions $\mathcal{R}^u_i$ or $\mathcal{R}^b_i$ become conversion reactions if any of the rate-limiting substrates $\mathbf{S^u_i}$, $\mathbf{A_i}$, or $\mathbf{B_i}$ are considerably consumed within the time interval of interest without being replenished (fast enough). In this case, one needs to account for such a substrate depletion.

Following from Figure 1 in the main text, corresponding resource-limited unimolecular conversion reactions can be simply obtained if the recovery arrows in olive color pointing towards $\mathbf{S^u_i}$ are removed. Similarly, the considered bimolecular catalytic production reaction turns to a bimolecular conversion one by removing the olive-colored incoming arrow to the intermediate $\mathbf{S^b_i}$ (in the first scenario), by removing the olive-colored incoming arrow to the species $\mathbf{B_i}$ (in the second scenario), or by removing at least one of the red arrows pointing towards the substrates $\mathbf{A_i}$ and $\mathbf{B_i}$ (in the third and fourth scenarios). Similar procedure applies to the other scenarios as well.

Our previous results for catalytic production reactions can be easily extended to these special cases, where some of the catalysts are being consumed, and some of the positive charges in the dynamics of the species $\mathbf{S_i^u}$, $\mathbf{S_i^b}$, $\mathbf{A_i}$ or $\mathbf{B_i}$ might be elliminated. To do so in the unimolecular cases, one needs to consider the addition of the decay rate $-v_i^u$ to the dynamic model of species $\mathbf{S_i^u}$. For modeling a resource-limited bimolecular conversion reaction $\mathcal{R}_i^b$, depending on which scenario it follows, one may need to consider adding the outflow $-v_i^b$ to the dynamics of either the species $\mathbf{A_i}$ alone, the species $\mathbf{B_i}$ alone, or both of $\mathbf{A_i}$ and $\mathbf{B_i}$ at the same time.

If $\mathcal{R}_i^b$ follows the first scenario and, hence, unlike $\mathbf{R}$ the substrate $\mathbf{S_i^b}$ now does not appear on the product side of the reaction set (S3) anymore, then both of $\mathbf{A_i}$ and $\mathbf{B_i}$ are being depleted with the same rate $-v_i^b$. If it is of second order and, hence, the species $\mathbf{B_i}$ does not recover in the product side of (S4), then one needs to consider $\mathbf{B_i}$ undergoing a decay with the negative drift $-v_i^b$. If $\mathcal{R}_i^b$ follows the third, fourth, fifth, or sixth scenario, then we shall have to consider the species that are no longer considered catalysts in the reaction set (S5), (S6), (S22), or (S23), respectively, as being consumed with a rate $-v_i^b$. We note that in the fifth and sixth scenarios, only the substrate species recycled in the last step—that is, $\mathbf{A_i}$ in the fifth and $\mathbf{B_i}$ in the sixth scenario which are recovered post-reaction through the red arrows in Figure S1—is allowed to be converted. The other substrate is by formulation supposed to act as a catalyst (follow from Figure S1 the recycling arrows in pink color).

In the unimolecular conversion case when the substrate $\mathbf{S_i^u}$ is in surplus such that small changes of it can be neglected, it essentially reduces to the zeroth-order catalytic production. Consider the example of transportation through transmembrane carrier proteins given in Section S2.1. Similarly, whenever one of the substrates $\mathbf{A_i}$ or $\mathbf{B_i}$ in a bimolecular conversion reaction is in excess and not rate-limiting, this reaction boils down to a unimolecular catalytic production form. See the ubiquitination mechanism given in Section S2.3 as an example.

#### S1.5 On the Effect of Diluting Intermediate Complexes

In the main manuscript in A Mathematical Framework to Model Intracellular Resource Competition, we do not explicitly account for the effect of concentration dilution due to cellular growth and division on the intermediary complexes. We rather consider this effect to be negligible, as it results in the loss of substrate species, avoiding the formulation of resource-limited catalytic production reactions. If the cell culture is maintained in the exponential growth phase with no consideration for the cell growth feedback coming into play by burdens on the resources $\mathbf{R}$, one may model this effect with a constant dilution term. It assumes that the cells grow in the exponential phase with a fixed growth rate. Denote this constant growth rate by $\delta$. Only considering constant degradation terms to account for the effect of cellular dilution, here we extend our resource-aware framework to the cases where the dilution of intermediate species is considerable.

For that, we assign to each intermediate complex $\mathbf{X}$ a positive parameter $\delta_x$, and to its dynamic model a negative constant degradation term as $-\delta_x x$. This parameter $\delta_x$ includes both the effect of global dilution of intracellular concentrations due to exponential cell growth as well as the native degradation of the species $\mathbf{X}$ within the cell's endogenous machinery (if non-negligible, for example if $\mathbf{X}$ represents a critically less stable protein). In our derivations, we limit ourselves to the bimolecular cases up to the first four scenarios, thus extending the entire results from Section S1.2 (we may refer to the previous result therein without explicitly mentioning).

We shall take that the following mass balance equation holds

$$r(t) = R_{\text{const}}(\delta) - \sum_{i=1}^{q_0} c_i^z(t) - \sum_{i=1}^{q_1} c_i^u(t) - \sum_{i=1}^{q_2} [c_i^b(t) + (\mathbb{1}_2(\zeta_i) + \mathbb{1}_4(\zeta_i)) s_i^b(t)]. \tag{S34}$$

The reason we have kept the total resource pool as constant does not mean that the increase in cell volume has no effect on it. Indeed, it does, and the faster the cell growth, the more the total resources get diluted, resulting in a lower $R_{\text{const}}$. However, we refrain from explicitly accounting for the dynamics of $R_{\text{const}}$ here. Instead, we consider it as a time-invariant function of $\delta$, having a fixed value for a given fixed $\delta$. Of course, if $\delta$ changes due to metabolic burden within a cell, for example, this value would also change. Within the time interval of interest, however, we shall consider $\delta$ as fixed. One could, on the other hand, accommodate the modeling of non-static total resources by associating a dynamical subsystem to it. This subsystem takes as input the current concentration of the species in the CRN, the dilution rate $\delta$, and possibly time $t$, and outputs the value of $R_{\text{const}}$ at time $t$, explicitly treating the effect of constant dilution within its dynamics. We note, however, that imposing quasi-steady state assumptions on this variable in such a setting would reduce it again to the above simplified formulation.

Similar to Section S1.2, let us hold again the main assumption A1 valid. Under this assumption, we update in what follows the quasi-steady state approximations previously derived through (S14)-(S17) in Section S1.2. For the resource-binding complexes $\mathbf{C_i^o}$ we have

$$c_i^z \approx \frac{k_{f_i}^z}{k_{r_i}^z + k_{c_i}^z + \delta_{c_i^z}} \bar{s}_i^z r, \tag{S35}$$

$$c_i^u \approx \frac{k_{f_i}^u}{k_{r_i}^u + k_{c_i}^u + \delta_{c_i^u}} s_i^u r, \tag{S36}$$

$$c_i^b \approx (\mathbb{1}_1(\zeta_i) + \mathbb{1}_3(\zeta_i)) \frac{k_{f_i}^b}{k_{r_i}^b + k_{c_i}^b + \delta_{c_i^b}} s_i^b r + (\mathbb{1}_2(\zeta_i) + \mathbb{1}_4(\zeta_i)) \frac{k_{f_i}^b}{k_{r_i}^b + k_{c_i}^b + \delta_{c_i^b}} s_i^b b_i. \tag{S37}$$

We mark the changes compared to the previous results in Section S1.2 in red. For the intermediate complexes $\mathbf{S_i^b}$, we have

$$\begin{aligned} s_i^b \approx\ & \mathbb{1}_1(\zeta_i) \frac{\eta_i a_i b_i}{\sigma_i + \delta_{s_i^b} + \dfrac{\delta_{c_i^b} k_{f_i}^b}{k_{r_i}^b + k_{c_i}^b + \delta_{c_i^b}} r} + \mathbb{1}_2(\zeta_i) \frac{\eta_i a_i r}{\sigma_i + \delta_{s_i^b} + \dfrac{\delta_{c_i^b} k_{f_i}^b}{k_{r_i}^b + k_{c_i}^b + \delta_{c_i^b}} b_i} \\ & + \mathbb{1}_3(\zeta_i) \frac{\eta_i a_i b_i}{\sigma_i + \delta_{s_i^b} + \dfrac{(k_{c_i}^b + \delta_{c_i^b}) k_{f_i}^b}{k_{r_i}^b + k_{c_i}^b + \delta_{c_i^b}} r} + \mathbb{1}_4(\zeta_i) \frac{\eta_i a_i r}{\sigma_i + \delta_{s_i^b} + \dfrac{(k_{c_i}^b + \delta_{c_i^b}) k_{f_i}^b}{k_{r_i}^b + k_{c_i}^b + \delta_{c_i^b}} b_i}. \end{aligned} \tag{S38}$$

Moreover, calculating from the dynamics in (S7)-(S12), we find that the dynamic models of the substrate catalysts are now affected by a negative drift, thus they are being diluted too with a rate potentially dependent on the available resource $\mathbf{R}$. Ideally, we would like to account for such a depletion in terms of the production rates $v_i^o$. Recall that $v_i^o = k_{c_i}^o c_i^o$, where $o \in \{z, u, b\}$. From the dynamic models (S7)-(S12) reported in Section S1.2, it is easy to see that the negative drifts have the forms $-\delta_{c_i^u} c_i^u$ and $-\delta_{c_i^b} c_i^b - \delta_{s_i^b} s_i^b$ for the unimolecular and bimolecular cases, respectively. In the bimolecular cases, both $\mathbf{A_i}$ and $\mathbf{B_i}$ undergo co-degradation with such a rate. From the above derivation, one can simplify these terms as

$$-\frac{\delta_{c_i^u}}{k_{c_i}^u} v_i^u \tag{S39}$$

and

$$-\left(\frac{\delta_{c_i^b}}{k_{c_i}^b}+\frac{\delta_{s_i^b}/k_{c_i}^b}{(\mathbb{1}_1(\zeta_i)+\mathbb{1}_3(\zeta_i))\frac{k_{f_i}^b}{k_{r_i}^b+k_{c_i}^b+\delta_{c_i^b}}r+(\mathbb{1}_2(\zeta_i)+\mathbb{1}_4(\zeta_i))\frac{k_{f_i}^b}{k_{r_i}^b+k_{c_i}^b+\delta_{c_i^b}}b_i}\right)v_i^b \tag{S40}$$

for the unimolecular and bimolecular cases, respectively. These expressions ease the accounting for the aforementioned substrate depletions. Later in this subsection, we provide the updated formulations of $v_i^o$s. For the zeroth-order cases, we do not account for the substrate depletion, as this would violate the assumption of constant inflows. Rather, we assume in these cases that the depletion of the substrate species is comparably minute and does not influence the constant rates $\bar{s}_i^z$ tangibly.

On substituting (S35)-(S38) into the equation (S34), one obtains the following quasi-steady state approximation for $r(t)$

$$r(t)\approx\frac{R_{\text{const}}(\delta)}{1+\sum_{i=1}^{q_0}\frac{\bar{s}_i^z}{k_{m_i}^z}+\sum_{i=1}^{q_1}\frac{s_i^u}{k_{m_i}^u}+\sum_{i=1}^{q_2}f^b(a_i,b_i,r,\zeta_i)}, \tag{S41}$$

where

$$\begin{aligned}
f^b &:= \frac{\mathbb{1}_1(\zeta_i)a_ib_i}{k_{m_i}^b\left(1+\frac{\delta_{c_i^b}k_{f_i}^b}{(\sigma_i+\delta_{s_i^b})(k_{r_i}^b+k_{c_i}^b+\delta_{c_i^b})}r\right)}+\frac{\mathbb{1}_2(\zeta_i)a_ib_i}{k_{m_i}^b\left(1+\frac{\delta_{c_i^b}k_{f_i}^b}{(\sigma_i+\delta_{s_i^b})(k_{r_i}^b+k_{c_i}^b+\delta_{c_i^b})}b_i\right)}+\zeta_i^a a_i \\
&\quad+\frac{\mathbb{1}_3(\zeta_i)a_ib_i}{k_{m_i}^b\left(1+\frac{(k_{c_i}^b+\delta_{c_i^b})k_{f_i}^b}{(\sigma_i+\delta_{s_i^b})(k_{r_i}^b+k_{c_i}^b+\delta_{c_i^b})}r\right)}+\frac{\mathbb{1}_4(\zeta_i)a_ib_i}{k_{m_i}^b\left(1+\frac{(k_{c_i}^b+\delta_{c_i^b})k_{f_i}^b}{(\sigma_i+\delta_{s_i^b})(k_{r_i}^b+k_{c_i}^b+\delta_{c_i^b})}b_i\right)}, \\
\zeta_i^a &:= \mathbb{1}_2(\zeta_i)\frac{\eta_i}{\sigma_i+\delta_{s_i^b}+\frac{\delta_{c_i^b}k_{f_i}^b}{k_{r_i}^b+k_{c_i}^b+\delta_{c_i^b}}b_i}+\mathbb{1}_4(\zeta_i)\frac{\eta_i}{\sigma_i+\delta_{s_i^b}+\frac{(k_{c_i}^b+\delta_{c_i^b})k_{f_i}^b}{k_{r_i}^b+k_{c_i}^b+\delta_{c_i^b}}b_i}, \\
k_{m_i}^z &:= (k_{r_i}^z+k_{c_i}^z+\delta_{c_i^z})/k_{f_i}^z, \\
k_{m_i}^u &:= (k_{r_i}^u+k_{c_i}^u+\delta_{c_i^u})/k_{f_i}^u,
\end{aligned} \tag{S42}$$

and

$$k_{m_i}^b := (k_{r_i}^b+k_{c_i}^b+\delta_{c_i^b})(\sigma_i+\delta_{s_i^b})/k_{f_i}^b\eta_i.$$

We shall adopt this approximation. Let us update the two simplifying assumptions A2 to A3 according to below.

**Assumptions.** *In the considered CRN, the following two conditions hold for every resource-limited bimolecular reaction:*

A2 *Each bimolecular reaction $\mathcal{R}_i^b$ of the first and third scenarios, following the representation given in (S3) or (S5), satisfies* $\sigma_i+\delta_{s_i^b}\gg(k_{c_i}^b+\delta_{c_i^b})k_{f_i}^b r(t)/(k_{r_i}^b+k_{c_i}^b+\delta_{c_i^b})$.

A3 *Each bimolecular reaction $\mathcal{R}_i^b$ of the second and fourth scenarios, following the representation given in (S4) or (S6), satisfies* $\sigma_i+\delta_{s_i^b}\gg(k_{c_i}^b+\delta_{c_i^b})k_{f_i}^b b_i(t)/(k_{r_i}^b+k_{c_i}^b+\delta_{c_i^b})$.

Under the assumptions A1-A3, we can then write out

$$v_i^z=\beta_i^z r \qquad \text{with } \beta_i^z=k_{c_i}^z\bar{s}_i^z/k_{m_i}^z, \tag{S43}$$

$$v_i^u = \beta_i^u s_i^u r \qquad \text{with } \beta_i^u = k_{c_i}^u / k_{m_i}^u,$$

$$v_i^b = \beta_i^b a_i b_i r \qquad \text{with } \beta_i^b = k_{c_i}^b / k_{m_i}^b,$$

where

$$r = \frac{R_{\text{const}}(\delta) \times (1+Q_0)^{-1}}{1 + \sum_{i=1}^{q_1} \frac{(1+Q_0)^{-1}}{k_{m_i}^u} s_i^u(t) + \sum_{i=1}^{q_2} \frac{(1+Q_0)^{-1}}{k_{m_i}^b} a_i(t) b_i(t) + \frac{\zeta_i^a}{1+Q_0} a_i(t)}. \tag{S44}$$

As can be seen, the above results (S43) and (S44) follow the same structure as those obtained in Section S1.2, more specifically the equations (S20) and (S21) reported therein. Aside from the depletion of substrates that should be accounted for in the substrate dynamics by employing the terms (S39) and (S40) above, the only differences are that $k_{m_i}^o$s are now updated constants scaled by the dilution-related constants $\delta_{c_i^o}$ and $\delta_{s_i^o}$, and the parameters $\zeta_i^a$ now have dependencies on $\delta_{s_i^o}$s in their denominators. This suggests that the considerations for the effect of diluting intermediate complexes keep the same form of formulation as with the catalytic productions discussed in the main text A Mathematical Framework to Model Intracellular Resource Competition, with the difference that now the substrates should no longer be treated as enzymatic catalysts.

#### S1.6 Extensions to Resource-Limited (Shared) Degradation/Sequestration Reactions

We now turn to the case of resource-limited degradation and resource-limited (irreversible) sequestration reactions. These reactions we model by referencing to an equivalent conversion reaction discussed in Section S1.4. Here, we call a reaction $\mathcal{R}_i^{\text{deg}}$ ($\mathcal{R}_i^{\text{seq}}$) a resource-limited *degradation* (*sequestration*) reaction if it has one (two) limiting substrate that is (substrates that both are) being depleted over time, and the presence of the shared resources $\mathbf{R}$ catalyzes the depletion(s). This equivalently follows a corresponding conversion reaction, with the only difference that now, instead of paying attention to the modeling of product species's dynamics, we merely focus on the changes in the consumed/depleting substrates. This, by induction, maps a resource-limited degradation (sequestration) reaction to an equivalent resource-limited unimolecular (bimolecular) conversion reaction (with no changes to the according scenario that it follows). A resource-limited degradation of zeroth-order is apparently not well defined, as in a degradation reaction we expect to lose at least one of the substrates over time, which is in contrary with the assumption of zeroth-order reactions, that is, the flow of its inflow reaction remains fixed over time. Let us disregard the effect of diluting intermediate complexes, though the results are readily generalizable.

Let $\mathcal{R}_i^{\text{deg}}$ and $\mathcal{R}_i^{\text{seq}}$ conform to the generic form of a resource-limited degradation reaction given by

$$\mathcal{R}_i^{\text{deg}} : \mathbf{S_i^u} \rightarrow \varnothing \quad \text{with } \mathrm{d}s_i^u/\mathrm{d}t = -v_i^{\text{deg}}(s_i^u, r), \tag{S45}$$

and a resource-limited sequestration reaction given by

$$\mathcal{R}_i^{\text{seq}} : \mathbf{A_i} + \mathbf{B_i} \rightarrow \varnothing \qquad \text{with } \mathrm{d}a_i/\mathrm{d}t = -v_i^{\text{seq}}(a_i, b_i, r) \quad \text{and} \quad \mathrm{d}b_i/\mathrm{d}t = -v_i^{\text{seq}}(a_i, b_i, r), \tag{S46}$$

respectively. In line with the settings in Section S1.2, let us take that, in the considered CRN of interest, there exist $q_1$ resource-

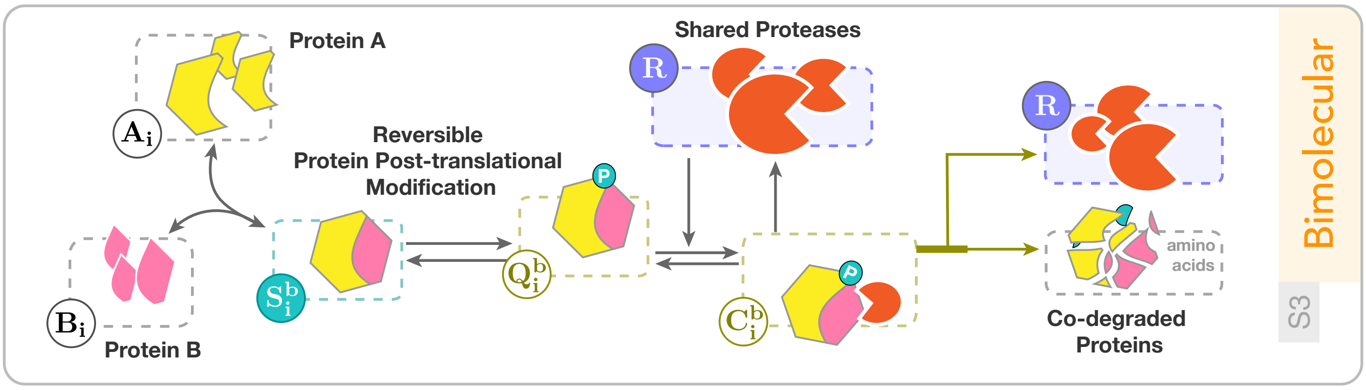

Figure S2: **A generic example of resource-limited sequestration reactions, which involves the reversible post-translational modifications of two proteins.** Here, both of the proteins A and B sequester each other depending on the availability of some shared protease complex, seen as the limited resource species $\mathbf{R}$.

limited degradation and $q_2$ resource-limited sequestration reactions that draw from $\mathbf{R}$. The goal is to obtain explicit formulas for the terms $v_i^{\text{deg}}$ and $v_i^{\text{seq}}$ based on the concentration of the substrate species, which are among the known species within the CRN under study. They apparently depend on all the species sharing $\mathbf{R}$. Assume that the resource-limited degradation reaction represented above follows the reaction-level representation in (S2) (substitute the product species $\mathbf{P_i^b}$ therein with $\varnothing$ to see this). Similarly, assume the sequestration reaction above follows either of the first four scenarios of the bimolecular reactions discussed through the equations (S3) to (S6). Let the relevant assumptions made therein also apply. This way, one can arrive at the following result straightforwardly by directly referencing the results obtained in Section S1.2

$$v_i^{\text{deg}} = \beta_i^u s_i^u r \qquad \text{with } \beta_i^u = k_{c_i}^u / k_{m_i}^u, \tag{S47}$$

$$v_i^{\text{seq}} = \beta_i^b a_i b_i r \qquad \text{with } \beta_i^b = k_{c_i}^b / k_{m_i}^b,$$

where

$$r = \frac{R_{\text{const}}}{1 + \sum_{i=1}^{q_1} \frac{s_i^u(t)}{k_{m_i}^u} + \sum_{i=1}^{q_2} \frac{a_i(t) b_i(t)}{k_{m_i}^b} + \zeta_i^a a_i(t)}. \tag{S48}$$

The parameters $\zeta_i^a$s are as previously defined in Section S1.2 corresponding to the bimolecular scenario that $\mathcal{R}_i^{\text{seq}}$s follow, so are $k_{m_i}^u$s and $k_{m_i}^b$s. In Figure S2, we illustrate a typical example of resource-limited sequestration reactions, where two proteins undergo post-translational modifications that enable for their co-degradation through proteases. See, for example, the phosphorylation-triggered protein degradation mechanisms in [1]. The protease in this scenario serves as the shared resource species, which may also be involved in other degradation activities as a catalyst. The availability of such a shared degradation facilitator affects the co-degradation of both substrate proteins, and the concentration of each protein directly limits the availability of the protease, from which tangled indirect couplings emerge. Note that if one of the reactants, either $\mathbf{A_i}$ or $\mathbf{B_i}$, exists in excess such that the impact of the co-degradation on it can be neglected, then the reaction simplifies to a corresponding resource-limited degradation reaction.

#### S1.7 Extensions to Biomolecular Reactions Limited by Two Different Resource Pools

Here, we extend the framework to resource-limited reactions constrained by the availability of two resource pools simultaneously. To distinguish, we shall call such reactions *bi-resource* limited. In contrast, we refer to those that solely depend on a single resource pool as *mono-resource* limited reactions. We only consider extensions of the results from Section S1.2, thus focusing solely on resource-limited catalytic production reactions. Generalizations to other extended settings discussed above are, however, straightforward. Let us define two different resource species, $\mathbf{R_1}$ and $\mathbf{R_2}$, to separately handle these two resource pools. Apparently, $\mathbf{R_1}$ might be shared both in mono- and bi-resource limited reactions simultaneously. The same holds for $\mathbf{R_2}$. How to address each case of mono-resource limited reactions based on the shared resources and corresponding substrate concentrations was already elaborated in previous sections (see Section S1.2). Thus, we will not repeat it here. What changes when considering the effect of simultaneous dependency on both $\mathbf{R_1}$ and $\mathbf{R_2}$ is that now the availability of $\mathbf{R_1}$ at any given time may depend on the availability of $\mathbf{R_2}$ at that moment, and vice versa. In this subsection, we expand upon our previous derivations to address these emerging intricacies.

Let us denote by $\bar{\mathcal{R}}_i$, $\bar{\bar{\mathcal{R}}}_i$, and $\tilde{\mathcal{R}}_i$ the mono-resource limited reactions within the CRN under consideration that draw solely from $\mathbf{R_1}$, solely from $\mathbf{R_2}$, and bi-resource limited reactions that draw simultaneously from both $\mathbf{R_1}$ and $\mathbf{R_2}$, respectively. Consider that, in the CRN of interest, there exist $\bar{q}$, $\bar{\bar{q}}$, and $\tilde{q}$ numbers of these three reactions, respectively. We will distinguish every term related to them—including their constitutive reactants and reaction rate constants—by using the same superscript notations. Let us think that each $\tilde{\mathcal{R}}_i$ conforms to one of the reaction types and scenarios drawn in Figure S3. We take that the reactions $\tilde{\mathcal{R}}_i$ conform to either the zeroth-order or unimolecular types provided in this figure. Only a single zeroth-order scenario is conceivable, whereas we consider six scenarios of the unimolecular types. Refer to Figure S3 for more details. Assume $\tilde{q}_0$ out of $\tilde{q}$ bi-resource limited reactions limited by the two resources are of zeroth-order types. Similarly, assume the rest, $\tilde{q}_1$, of them follow either of the six unimolecular scenarios illustrated in Figure S3. Let us notationally discriminate between $\tilde{\mathcal{R}}_i$s of zeroth-order and unimolecular types by $\tilde{\mathcal{R}}_i^z$ and $\tilde{\mathcal{R}}_i^u$, respectively. Let the following generic forms represent

$$\varnothing \rightarrow \tilde{\mathbf{P}}_{\mathbf{i}}^{\mathbf{z}} \qquad \text{with } \mathrm{d}\tilde{p}_i^z/\mathrm{d}t = \tilde{v}_i^z(r_1, r_2), \tag{S49}$$

$$\tilde{\mathbf{S}}_{\mathbf{i}}^{\mathbf{u}} \rightarrow \tilde{\mathbf{S}}_{\mathbf{i}}^{\mathbf{u}} + \tilde{\mathbf{P}}_{\mathbf{i}}^{\mathbf{u}} \qquad \text{with } \mathrm{d}p_i^u/\mathrm{d}t = \tilde{v}_i^u(\tilde{s}_i^u, r_1, r_2), \tag{S50}$$

the bi-resource limited zeroth-order and unimolecular reactions $\tilde{\mathcal{R}}_i^z$ and $\tilde{\mathcal{R}}_i^u$, respectively. Needless to say, the species $\tilde{\mathbf{S}}_{\mathbf{i}}^{\mathbf{u}}$ are the substrates limiting the production of $\tilde{\mathbf{P}}_{\mathbf{i}}^{\mathbf{u}}$ through $\tilde{\mathcal{R}}_i^u$s.

Our objective is to find explicit formulae for $\tilde{v}_i^o$, $o \in \{z, u\}$, based solely on the substrate concentrations $\tilde{s}_i^u$ as well as all the other substrate concentrations that share $\mathbf{R_1}$ or $\mathbf{R_2}$ through mono-resource limited reactions. To that end, we shall again find and adopt quasi-steady state approximations of $r_1$ and $r_2$ based on these substrate species, similar to the previous derivations in Section S1.2. We keep the main assumptions made in A Mathematical Framework to Model Intracellular Resource Competition to hold for all the mono-resource limited reactions $\bar{\mathcal{R}}_i$ and $\bar{\bar{\mathcal{R}}}_i$. Additionally, we shall assume that similar assumptions adapted to the formulation of the bi-resource limited reactions also hold, including, for example, that the resource-binding complexes $\tilde{\mathbf{C}}_{\mathbf{i}}^{\mathbf{z}}$ and $\tilde{\mathbf{C}}_{\mathbf{i}}^{\mathbf{u}}$ together with the intermediate complexes $\tilde{\mathbf{S}}_{\mathbf{i}}^{\mathbf{u}}$ are all at dynamic equilibrium (for every $i$).

Given the setting described above and based on the derivations in Section S1.2, one can readily arrive at the following

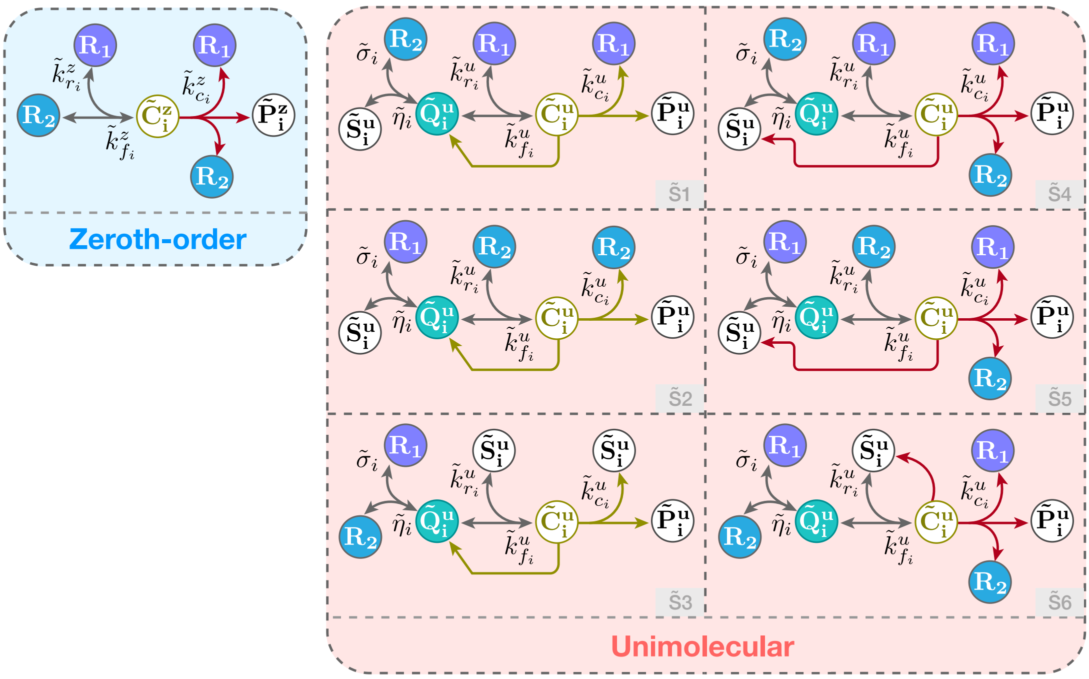

Figure S3: **Bi-resource limited catalytic production reactions.** On the left, we symbolically depict the CRN representation of the zeroth-order types of such reactions, which we denote by $\tilde{\mathcal{R}}_i^z$. They require the simultaneous availability of two different resources to catalyze the production of a species $\tilde{\mathbf{P}}_\mathbf{i}^\mathbf{z}$. The first three scenarios of bi-resource limited unimolecular reactions are drawn in the left column. We denote the $i$th reaction of such by $\tilde{\mathcal{R}}_i^u$. The right column continues with the unimolecular cases to scenarios where the post-reaction recovery of the substrate species and the catalyzing resources are direct. Note that, consistently throughout our extension, we distinguish between bi-resource and mono-resource limited reactions and their related mathematical formulations using the superscript notation "$\sim$".

relationships for $\tilde{v}_i^o$

$$\tilde{v}_i^z = \tilde{\beta}_i^z r_1 r_2 \qquad \text{with } \tilde{\beta}_i^z = \tilde{k}_{c_i}^z / \tilde{k}_{m_i}^z, \tag{S51}$$

$$\tilde{v}_i^u = \tilde{\beta}_i^u \tilde{s}_i^u r_1 r_2 \qquad \text{with } \tilde{\beta}_i^u = \tilde{k}_{c_i}^u / \tilde{k}_{m_i}^u,$$

and formulate the following expressions for the available $\mathbf{R_1}$ and $\mathbf{R_2}$

$$r_1(t) = \frac{R_{tot1}}{1 + \underbrace{\sum_{i=1}^{\bar{q}_1} \bar{w}_i^u \bar{s}_i^u(t) + \sum_{i=1}^{\bar{q}_2} \bar{w}_i^{ab} \bar{a}_i(t) \bar{b}_i(t) + \bar{w}_i^a \bar{a}_i(t)}_{f_{R_1}(\bar{\boldsymbol{s}}^{\boldsymbol{u}}, \bar{\boldsymbol{a}}, \bar{\boldsymbol{b}})} + \sum_{i=1}^{\tilde{q}_0} \tilde{w}_i^z r_2(t) + \sum_{i=1}^{\tilde{q}_1} \underbrace{\tilde{w}_i^{u_{11}} \tilde{s}_i^u(t) r_2(t) + \tilde{w}_i^{u_{12}} \tilde{s}_i^u(t) + \tilde{w}_i^{u_{13}} r_2(t)}_{h_{R_1}^i(\tilde{s}_i^u, r_2)}}, \tag{S52}$$

and

$$r_2(t) = \frac{R_{tot2}}{1 + \underbrace{\sum_{i=1}^{\bar{\bar{q}}_1} \bar{\bar{w}}_i^u \bar{\bar{s}}_i^u(t) + \sum_{i=1}^{\bar{\bar{q}}_2} \bar{\bar{w}}_i^{ab} \bar{\bar{a}}_i(t) \bar{\bar{b}}_i(t) + \bar{\bar{w}}_i^a \bar{\bar{a}}_i(t)}_{f_{R_2}(\bar{\bar{\boldsymbol{s}}}^{\boldsymbol{u}}, \bar{\bar{\boldsymbol{a}}}, \bar{\bar{\boldsymbol{b}}})} + \sum_{i=1}^{\tilde{q}_0} \tilde{w}_i^z r_1(t) + \sum_{i=1}^{\tilde{q}_1} \underbrace{\tilde{w}_i^{u_{21}} \tilde{s}_i^u(t) r_1(t) + \tilde{w}_i^{u_{22}} \tilde{s}_i^u(t) + \tilde{w}_i^{u_{23}} r_1(t)}_{h_{R_2}^i(\tilde{s}_i^u, r_1)}}, \tag{S53}$$

respectively. For exact expressions for the summands $h_{R_1}^i(\tilde{s}_i^u, r_1)$ and $h_{R_2}^i(\tilde{s}_i^u, r_2)$, one needs to take a deeper look at the mappings between the bi-resource limited unimolecular reactions in Figure S3 and the mono-resource limited bimolecular scenarios in Figure 1 in A Mathematical Framework to Model Intracellular Resource Competition. We have done this mapping reported in Table S1. The way we do the mapping is that we first think of $\mathbf{R_1}$ as the limiting resource and $\mathbf{R_2}$ as a substrate, and view the dynamics of $r_1$ through the lens of a corresponding mono-resource limited bimolecular scenario. We then repeat the same procedure by flipping $\mathbf{R_1}$ and $\mathbf{R_2}$, viewing everything now through the lens of $\mathbf{R_2}$ being the shared resource. See Table S1 for a report of the expressions for $h^i_{R_1}$ and $h^i_{R_2}$. As can be found therefrom, for every $\tilde{\mathcal{R}}^u_i$ there exist corresponding strictly positive competition gains $\tilde{w}^{u_{11}}_i$ and $\tilde{w}^{u_{21}}_i$, and that

$$\tilde{w}^{u_{11}}_i = \tilde{w}^{u_{21}}_i \tag{S54a}$$

holds. Moreover, for every scenario $\tilde{\text{S}}1$-$\tilde{\text{S}}6$ the following relationships always hold for every $i$

$$\tilde{w}^{u_{13}}_i = \tilde{w}^{u_{23}}_i, \tag{S54b}$$

$$\tilde{w}^{u_{12}}_i \tilde{w}^{u_{22}}_i = 0. \tag{S54c}$$

To avoid clutter, we will not explicitly write down the expressions of the lumped parameters such as $\tilde{\beta}^o_i$ or the competition gains involved. By induction from Section S1.2, it is an easy task, however, to compute them in terms of the constituting reaction rate constants. For now, think of them as some non-negative constants. As can be seen from (S52) and (S53), the two available resources are now tangled with each other, each tightly depending on the other.

Our task is to untangle these dependencies. The functions $f_{R_1}$ and $f_{R_2}$ are defined to simplify the notation. For compact writing, their input arguments are written in vector forms, with $\boldsymbol{X}$ being a vector comprising all the corresponding $x_i$s. Given the above findings, the equalities in (S54), and Table S1, one can write the following relation by subtracting the equality (S52) from (S53) and canceling the common terms

$$r_1\left(1 + f_{R_1}(\bar{\boldsymbol{s}}^{\boldsymbol{u}}, \bar{\boldsymbol{a}}, \bar{\boldsymbol{b}}) + \sum_{i=1}^{\tilde{q}_1} \tilde{w}^{u_{12}}_i \tilde{s}^u_i\right) - r_2\left(1 + f_{R_2}(\bar{\bar{\boldsymbol{s}}}^{\boldsymbol{u}}, \bar{\bar{\boldsymbol{a}}}, \bar{\bar{\boldsymbol{b}}}) + \sum_{i=1}^{\tilde{q}_1} \tilde{w}^{u_{22}}_i \tilde{s}^u_i\right) = R_{tot1} - R_{tot2}. \tag{S55}$$

Accordingly, one can recast (S52) and (S53) as below

$$r_1 = \frac{R_{tot1}}{1 + f_{R_1}(\bar{\boldsymbol{s}}^{\boldsymbol{u}}, \bar{\boldsymbol{a}}, \bar{\boldsymbol{b}}) + \sum_{i=1}^{\tilde{q}_1} \tilde{w}^{u_{12}}_i \tilde{s}^u_i + \left(\sum_{i=1}^{\tilde{q}_0} \tilde{w}^z_i + \sum_{i=1}^{\tilde{q}_1} \tilde{w}^{u_{11}}_i \tilde{s}^u_i + \tilde{w}^{u_{13}}_i\right) \dfrac{r_1 g_{R_1} - (R_{tot1} - R_{tot2})}{g_{R_2}}}, \tag{S56}$$

and

$$r_2 = \frac{R_{tot2}}{1 + f_{R_2}(\bar{\bar{\boldsymbol{s}}}^{\boldsymbol{u}}, \bar{\bar{\boldsymbol{a}}}, \bar{\bar{\boldsymbol{b}}}) + \sum_{i=1}^{\tilde{q}_1} \tilde{w}^{u_{22}}_i \tilde{s}^u_i + \left(\sum_{i=1}^{\tilde{q}_0} \tilde{w}^z_i + \sum_{i=1}^{\tilde{q}_1} \tilde{w}^{u_{11}}_i \tilde{s}^u_i + \tilde{w}^{u_{13}}_i\right) \dfrac{r_2 g_{R_2} + (R_{tot1} - R_{tot2})}{g_{R_1}}}, \tag{S57}$$

where

$$g_{R_1}(\bar{\boldsymbol{s}}^{\boldsymbol{u}}, \bar{\boldsymbol{a}}, \bar{\boldsymbol{b}}, \tilde{\boldsymbol{s}}^{\boldsymbol{u}}) := \left(1 + f_{R_1}(\bar{\boldsymbol{s}}^{\boldsymbol{u}}, \bar{\boldsymbol{a}}, \bar{\boldsymbol{b}}) + \sum_{i=1}^{\tilde{q}_1} \tilde{w}^{u_{12}}_i \tilde{s}^u_i\right), \tag{S58}$$

Unimolecular Scenarios from Figure S3

| Scenario | Mapping $\mathbf{R_1}$ | Mapping $\mathbf{R_2}$ | Corresponding $h^i_{R_1}(\tilde{s}^u_i, r_2)$ | Corresponding $h^i_{R_2}(\tilde{s}^u_i, r_1)$ |
|---|---|---|---|---|
| $\tilde{\text{S}}1$ | S1 | S2 | $\tilde{w}^{u_{11}}_i \tilde{s}^u_i r_2$ | $\tilde{w}^{u_{21}}_i \tilde{s}^u_i r_1 + \tilde{w}^{u_{22}}_i \tilde{s}^u_i$ |
| $\tilde{\text{S}}2$ | S2 | S1 | $\tilde{w}^{u_{11}}_i \tilde{s}^u_i r_2 + \tilde{w}^{u_{12}}_i \tilde{s}^u_i$ | $\tilde{w}^{u_{21}}_i \tilde{s}^u_i r_1$ |
| $\tilde{\text{S}}3$ | S2 | S2 | $\tilde{w}^{u_{11}}_i \tilde{s}^u_i r_2 + \tilde{w}^{u_{13}}_i r_2$ | $\tilde{w}^{u_{21}}_i \tilde{s}^u_i r_1 + \tilde{w}^{u_{23}}_i r_1$ |
| $\tilde{\text{S}}4$ | S3 | S4 | $\tilde{w}^{u_{11}}_i \tilde{s}^u_i r_2$ | $\tilde{w}^{u_{21}}_i \tilde{s}^u_i r_1 + \tilde{w}^{u_{22}}_i \tilde{s}^u_i$ |
| $\tilde{\text{S}}5$ | S4 | S3 | $\tilde{w}^{u_{11}}_i \tilde{s}^u_i r_2 + \tilde{w}^{u_{12}}_i \tilde{s}^u_i$ | $\tilde{w}^{u_{21}}_i \tilde{s}^u_i r_1$ |
| $\tilde{\text{S}}6$ | S4 | S4 | $\tilde{w}^{u_{11}}_i \tilde{s}^u_i r_2 + \tilde{w}^{u_{13}}_i r_2$ | $\tilde{w}^{u_{21}}_i \tilde{s}^u_i r_1 + \tilde{w}^{u_{23}}_i r_1$ |

Table S1: **Mappings between bi-resource limited unimolecular reactions and mono-resource limited bimolecular reactions.** For a given reaction $\tilde{\mathcal{R}}^u_i$ of the former cases, the first column indicates its corresponding scenario according to the six scenarios provided in Figure S3. The second column aims to provide a mapping between the scenario in the first column and the four bimolecular scenarios of the mono-resource limited reactions, correspondingly. These four bimolecular scenarios were previously denoted by S1-S4 and discussed in Sections S1.1 and S1.2. The illustrations of these four scenarios are available in Figure 1 from A Mathematical Framework to Model Intracellular Resource Competition. In the second column, we essentially report two entries. The first entry refers to the bimolecular scenario that $\tilde{\mathcal{R}}^u_i$ would follow if considering $\mathbf{R_1}$ as the single resource pool, treating the other resource species as a substrate. The second entry refers to the same but with $\mathbf{R_2}$ being the limiting resource species. For example, the scenario $\tilde{\text{S}}1$ in Figure S3 maps to the scenario S4 in Figure 1 and follows the representative CRN form in (S6), if we consider $\mathbf{R_2}$ as the shared resource species. Conversely, it would follow S3 in (S5) if we consider $\mathbf{R_1}$ to be the resource species $\mathbf{R}$ therein. The last two columns report the resulting summands $h^i_{R_1}(\tilde{s}^u_i, r_1)$ and $h^i_{R_2}(\tilde{s}^u_i, r_2)$ that appear in the denominators of (S52) and (S53), respectively.

$$g_{R_2}(\bar{\boldsymbol{s}}^{\boldsymbol{u}}, \bar{\bar{\boldsymbol{a}}}, \bar{\bar{\boldsymbol{b}}}, \tilde{\boldsymbol{s}}^{\boldsymbol{u}}) := \left(1 + f_{R_2}(\bar{\bar{\boldsymbol{s}}}^{\boldsymbol{u}}, \bar{\bar{\boldsymbol{a}}}, \bar{\bar{\boldsymbol{b}}}) + \sum_{i=1}^{\tilde{q}_1} \tilde{w}^{u_{22}}_i \tilde{s}^u_i\right). \tag{S59}$$

Notice that in the above, $r_1(t)$ does not depend on $r_2(t)$, and vice versa. This paves the way for solving $r_1$ ($r_2$) solely in terms of all the substrates influencing it, without explicit reliance on $r_2$ ($r_1$ ). From (S56)-(S57), one can easily see that $r_1$ and $r_2$ are the solutions of the following quadratic equations

$$r_1^2\left(\left(\sum_{i=1}^{\tilde{q}_0} \tilde{w}^z_i + \sum_{i=1}^{\tilde{q}_1} \tilde{w}^{u_{11}}_i \tilde{s}^u_i + \tilde{w}^{u_{13}}_i\right)\frac{g_{R_1}}{g_{R_2}}\right) + r_1 \mathcal{Y}_{R_1}(\bar{\boldsymbol{s}}^{\boldsymbol{u}}, \bar{\boldsymbol{a}}, \bar{\boldsymbol{b}}, \bar{\bar{\boldsymbol{s}}}^{\boldsymbol{u}}, \bar{\bar{\boldsymbol{a}}}, \bar{\bar{\boldsymbol{b}}}, \tilde{\boldsymbol{s}}^{\boldsymbol{u}}) - R_{tot1} = 0, \tag{S60}$$

and

$$r_2^2\left(\left(\sum_{i=1}^{\tilde{q}_0} \tilde{w}^z_i + \sum_{i=1}^{\tilde{q}_1} \tilde{w}^{u_{11}}_i \tilde{s}^u_i + \tilde{w}^{u_{13}}_i\right)\frac{g_{R_2}}{g_{R_1}}\right) + r_2 \mathcal{Y}_{R_2}(\bar{\boldsymbol{s}}^{\boldsymbol{u}}, \bar{\boldsymbol{a}}, \bar{\boldsymbol{b}}, \bar{\bar{\boldsymbol{s}}}^{\boldsymbol{u}}, \bar{\bar{\boldsymbol{a}}}, \bar{\bar{\boldsymbol{b}}}, \tilde{\boldsymbol{s}}^{\boldsymbol{u}}) - R_{tot2} = 0, \tag{S61}$$

where

$$\mathcal{Y}_{R_1} := \left(1 + f_{R_1}(\bar{\boldsymbol{s}}^{\boldsymbol{u}}, \bar{\boldsymbol{a}}, \bar{\boldsymbol{b}}) + \sum_{i=1}^{\tilde{q}_1} \tilde{w}^{u_{12}}_i \tilde{s}^u_i - \left(\sum_{i=1}^{\tilde{q}_0} \tilde{w}^z_i + \sum_{i=1}^{\tilde{q}_1} \tilde{w}^{u_{11}}_i \tilde{s}^u_i + \tilde{w}^{u_{13}}_i\right)\frac{R_{tot1} - R_{tot2}}{g_{R_2}}\right), \tag{S62}$$

$$\mathcal{Y}_{R_2} := \left(1 + f_{R_2}(\bar{\bar{\boldsymbol{s}}}^{\boldsymbol{u}}, \bar{\bar{\boldsymbol{a}}}, \bar{\bar{\boldsymbol{b}}}) + \sum_{i=1}^{\tilde{q}_1} \tilde{w}_i^{u_{22}} \tilde{s}_i^u + \left(\sum_{i=1}^{\tilde{q}_0} \tilde{w}_i^z + \sum_{i=1}^{\tilde{q}_1} \tilde{w}_i^{u_{11}} \tilde{s}_i^u + \tilde{w}_i^{u_{13}}\right) \frac{R_{tot1} - R_{tot2}}{g_{R_1}}\right). \tag{S63}$$

Looking at the quadratic formulations (S60) and (S61), it is easy to see that both always guarantee a unique positive solution for $r_1$ and $r_2$, regardless of the signs of $\mathcal{Y}_{R_i}$s. One can compactly write these solutions as follows

$$r_1 = \frac{-\mathcal{Y}_{R_1}(\bar{\boldsymbol{s}}^{\boldsymbol{u}}, \bar{\boldsymbol{a}}, \bar{\boldsymbol{b}}, \bar{\bar{\boldsymbol{s}}}^{\boldsymbol{u}}, \bar{\bar{\boldsymbol{a}}}, \bar{\bar{\boldsymbol{b}}}, \tilde{\boldsymbol{s}}^{\boldsymbol{u}}) + \sqrt{\mathcal{Y}_{R_1}^2(\bar{\boldsymbol{s}}^{\boldsymbol{u}}, \bar{\boldsymbol{a}}, \bar{\boldsymbol{b}}, \bar{\bar{\boldsymbol{s}}}^{\boldsymbol{u}}, \bar{\bar{\boldsymbol{a}}}, \bar{\bar{\boldsymbol{b}}}, \tilde{\boldsymbol{s}}^{\boldsymbol{u}}) + 4R_{tot1}\left(\sum_{i=1}^{\tilde{q}_0} \tilde{w}_i^z + \sum_{i=1}^{\tilde{q}_1} \tilde{w}_i^{u_{11}} \tilde{s}_i^u + \tilde{w}_i^{u_{13}}\right)\frac{g_{R_1}}{g_{R_2}}}}{\frac{2g_{R_1}}{g_{R_2}}\left(\sum_{i=1}^{\tilde{q}_0} \tilde{w}_i^z + \sum_{i=1}^{\tilde{q}_1} \tilde{w}_i^{u_{11}} \tilde{s}_i^u + \tilde{w}_i^{u_{13}}\right)}, \tag{S64}$$

and

$$r_2 = \frac{-\mathcal{Y}_{R_2}(\bar{\boldsymbol{s}}^{\boldsymbol{u}}, \bar{\boldsymbol{a}}, \bar{\boldsymbol{b}}, \bar{\bar{\boldsymbol{s}}}^{\boldsymbol{u}}, \bar{\bar{\boldsymbol{a}}}, \bar{\bar{\boldsymbol{b}}}, \tilde{\boldsymbol{s}}^{\boldsymbol{u}}) + \sqrt{\mathcal{Y}_{R_2}^2(\bar{\boldsymbol{s}}^{\boldsymbol{u}}, \bar{\boldsymbol{a}}, \bar{\boldsymbol{b}}, \bar{\bar{\boldsymbol{s}}}^{\boldsymbol{u}}, \bar{\bar{\boldsymbol{a}}}, \bar{\bar{\boldsymbol{b}}}, \tilde{\boldsymbol{s}}^{\boldsymbol{u}}) + 4R_{tot2}\left(\sum_{i=1}^{\tilde{q}_0} \tilde{w}_i^z + \sum_{i=1}^{\tilde{q}_1} \tilde{w}_i^{u_{11}} \tilde{s}_i^u + \tilde{w}_i^{u_{13}}\right)\frac{g_{R_2}}{g_{R_1}}}}{\frac{2g_{R_2}}{g_{R_1}}\left(\sum_{i=1}^{\tilde{q}_0} \tilde{w}_i^z + \sum_{i=1}^{\tilde{q}_1} \tilde{w}_i^{u_{11}} \tilde{s}_i^u + \tilde{w}_i^{u_{13}}\right)}, \tag{S65}$$

where $\mathcal{Y}_{R_1}^2$ and $\mathcal{Y}_{R_2}^2$ are provided in (S63), and $g_{R_1}$ and $g_{R_2}$ are given in (S58) and (S59), respectively. This concludes our task, as we have fully characterized $r_1$, $r_2$, and accordingly all the production rates $\tilde{v}_i^o$, based solely on the substrate species sharing the two resources, whether sharing through mono- or bi-resource limited reactions.

### S2 Biological Examples and Interpretations on the Introduced Framework

This section provides for each type of resource-limited production reactions considered in the manuscript, a minimum of two different biological realizations. These examples are visually represented through Figure S4 to Figure S7. The notation and terminology used in this section are consistent with A Mathematical Framework to Model Intracellular Resource Competition of the main text and also Section S1 above. We only focus on resource-limited reactions constrained by single resource pools. In the sequel, we will also review various types of the above-discussed resource-limited reactions and the reaction networks associated to them.

#### S2.1 Resource-limited Zeroth-order Production Reactions

For a zeroth-order production reaction $\mathcal{R}_i^z$, which is limited by the availability of resources $\mathbf{R}$, a constant-in-rate inflow (whose rate is denoted by $\bar{s}_i^z \in \mathbb{R}_{>0}$) forms an active intermediary complex $\mathbf{C_i^z}$. However, this process requires the participation of another species, $\mathbf{R}$. The production rate of the intermediate complex is, thus, limited by the availability of $\mathbf{R}$. We think of $\mathbf{R}$ as the shared resource species. This complex then transforms into the product $\mathbf{P_i^z}$ following the reactions

$$\mathrm{G} + \mathbf{R} \underset{k_{r_i}^z}{\overset{k_{f_i}^z \bar{s}_i^z}{\rightleftharpoons}} \mathbf{C_i^z} \xrightarrow{k_{c_i}^z} \mathbf{R} + \mathbf{P_i^z}. \tag{S66}$$

Note that $\mathbf{P_i^z}$ is being produced while the species $\mathbf{R}$ is not consumed and the flow rate $\bar{s}_i^z$ remains fixed. This essentially requires some other species to be consumed in between so that the production of $\mathbf{P_i^z}$ is possible. We have grouped all such underlying, but hidden, essential substances together as a single substrate species, called $\mathbf{G}$. Indeed, we have implicitly assumed that these additional substrates are available in excessive amounts so that the consumption of them (in small quantities) does not affect their whole concentration within the time interval of interest. This means that the species $\mathbf{G}$ is not a limiting substrate in the considered reactions. The faint color in (S66) is used to indicate this facet. Think of them as being continuously refilled (in large quantities) through other processes happening outside of the considered CRN of interest, for example. Note that, without loss of generality and in the zeroth-order cases, it is possible to consider that the constant inflow rate $\bar{s}_i^z$ originates from those excluded species $\mathbf{G}$.

Depending on the context and the biological system at hand, a range of possible realizations for $\mathbf{G}$ might include, but are not limited to: Adenosine triphosphate (abbreviated "ATP"), nucleotides, amino acids, and etc. If the cells receive enough nutrients such that their housekeeping genes are actively producing sufficient essential elements to sustain cell viability and maintain its existence, then such examples can be valid components to be grouped as the species $\mathbf{G}$. Otherwise, if, for example, the cells are not receiving enough high-quality nutrients and have entered a starvation state, or are already transitioning into the death phase where their death rate exceeds their replication rate for extended periods within the time interval of interest, then grouping some or all of the above examples as the species $\mathbf{G}$ may no longer be appropriate. In this section, we generally assume the cell culture growth is in the exponential phase within the time interval of interest, unless otherwise specified.

For the first example, we think of the transportation of an extracellular molecule into the cytoplasm as an instance of resource-limited zeroth-order reaction. That is, consider the transmembrane carrier proteins, which are available in limited quantities on the plasma membrane, as the restricting resources required for the transport of specific molecules, solutes, or analytes, such as carbon sources, from the extracellular space (ECS) into the cell. If available in limited quantities, transmembrane carrier proteins—whether uniporters or multiporters (the latter capable of transporting more than one type of molecule, solute, or analyte simultaneously)—will face competition for binding. Notably, the reaction follows zeroth-order kinetics if the extracellular molecules are available in abundant quantities, such that the removal of a few does not significantly change the inflow rate, $\bar{s}_i^z$. Note that the carrier proteins bind to this specific substance first, and undergo conformational changes then, to ease the procedure. The resulting bound between porter and the uptaken molecule is deemed as the intermediary resource-substrate complex $\mathbf{C_i^z}$. Once transported, we treat this just-passed-through-the-membrane molecule as the desired product $\mathbf{P_i^z}$. This way, the chemical reactions in the form of (S66) follow. Illustrations can be found in Figure S4A. Note that the steep concentration gradient across the membrane will render the reverse reaction $\mathbf{C_i^z} \xrightarrow{k_{r_i}^z} \mathbf{R}$ in (S66) highly unlikely, which will therefore be disregarded. In this example, dihydrogen monoxide can be considered as a trivial candidate for the additional substrate $\mathbf{G}$, as the biochemical reactions typically occur in the presence of abundant water molecules, making it a non-rate-limiting factor.

As the second example, in Figure S4B we consider the DNA in nucleus as a constant quantity (the constant rate $\bar{s}_i^z$), which requires as the shared resource an RNA polymerase (RNAP in short) in order to start the transcription process. It starts after the transcription initiation complex, an intermediary active species represented by $\mathbf{C_i^z}$, is formed. A product of this process, $\mathbf{P_i^z}$, is the messenger RNA (abbreviated mRNA). At the end of the procedure, the occupied RNA polymerase (RNAP in short) will be released and return to the resource pool (highlighting the assumption that the total resources are conserved). Nucleotides can be seen as the species $\mathbf{G}$.

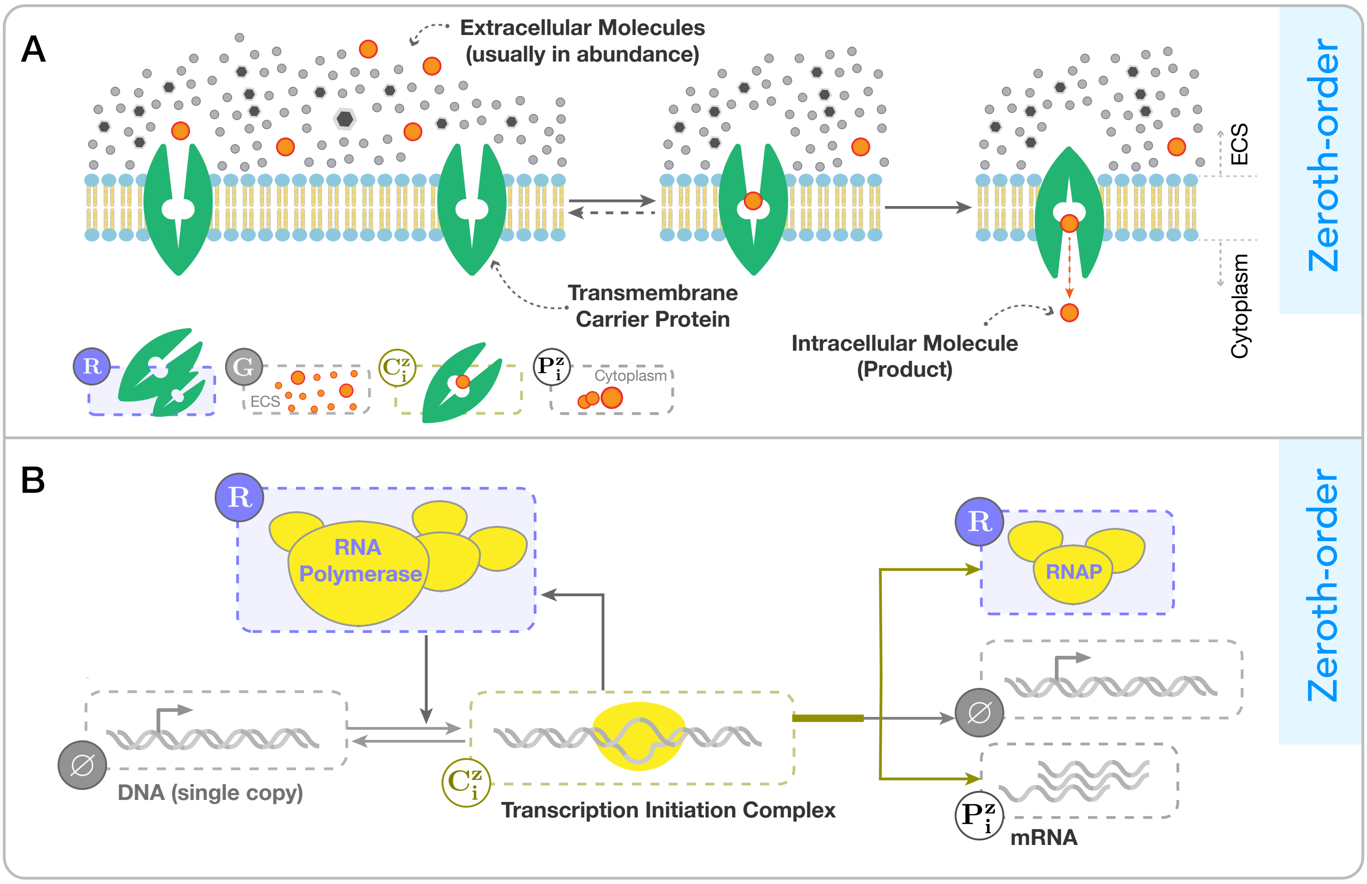

Figure S4: **Two examples of biochemical processes that illustrate resource-limited zeroth-order production reactions.** (**A**) The transportation process of a cognate analyte, solute, or biomolecule from the extracellular space (ECS) into the cytoplasm through transmembrane carrier proteins. (**B**) The transcription process of a gene's DNA sequence.

#### S2.2 Resource-limited Unimolecular Production Reactions

For the unimolecular catalytic production reaction $\mathcal{R}_i^u$ limited by free available $\mathbf{R}$, we follow the general form

$$\mathbf{G} + \mathbf{S_i^u} + \mathbf{R} \underset{k_{r_i}^u}{\overset{k_{f_i}^u}{\rightleftharpoons}} \mathbf{C_i^u} \xrightarrow{k_{c_i}^u} \mathbf{S_i^u} + \mathbf{R} + \mathbf{P_i^u}. \tag{S67}$$

It is now recognized that the substrate concentration can vary over time, rather than remaining constant as is an assumption with the zeroth-order types.

The first example, as shown in Figure S5A, provides a simplified description of the translation process as a unimolecular reaction that is limited by the availability of translational resources. These resources, which collectively function as the shared resources represented by $\mathbf{R}$, may include ribosomal subunits, initiation factors, aminoacyl-tRNAs (aa-tRNAs), and other non-coding RNAs or proteins involved in the process. The availability of these translational resources may be limited due to factors such as their shared use by other genetic components, cellular stress, growth, or exposure to specific drugs. In this example, the target mRNA is regarded as the rate-limiting substrate $\mathbf{S_i^u}$ (whose concentration can be influenced, for example, by downstream transcription processes that produce the same RNA sequence). The process involves the formation of a translationally active complex $\mathbf{C_i^u}$ between mRNA and translational resources, which slowly produces a (folded) protein $\mathbf{P_i^u}$ as the end product. In this example, $\mathbf{G}$ can be taken as ATPs and amino acids.

In the second example, depicted in Figure S5B, we consider a more detailed model of the previously discussed transcription

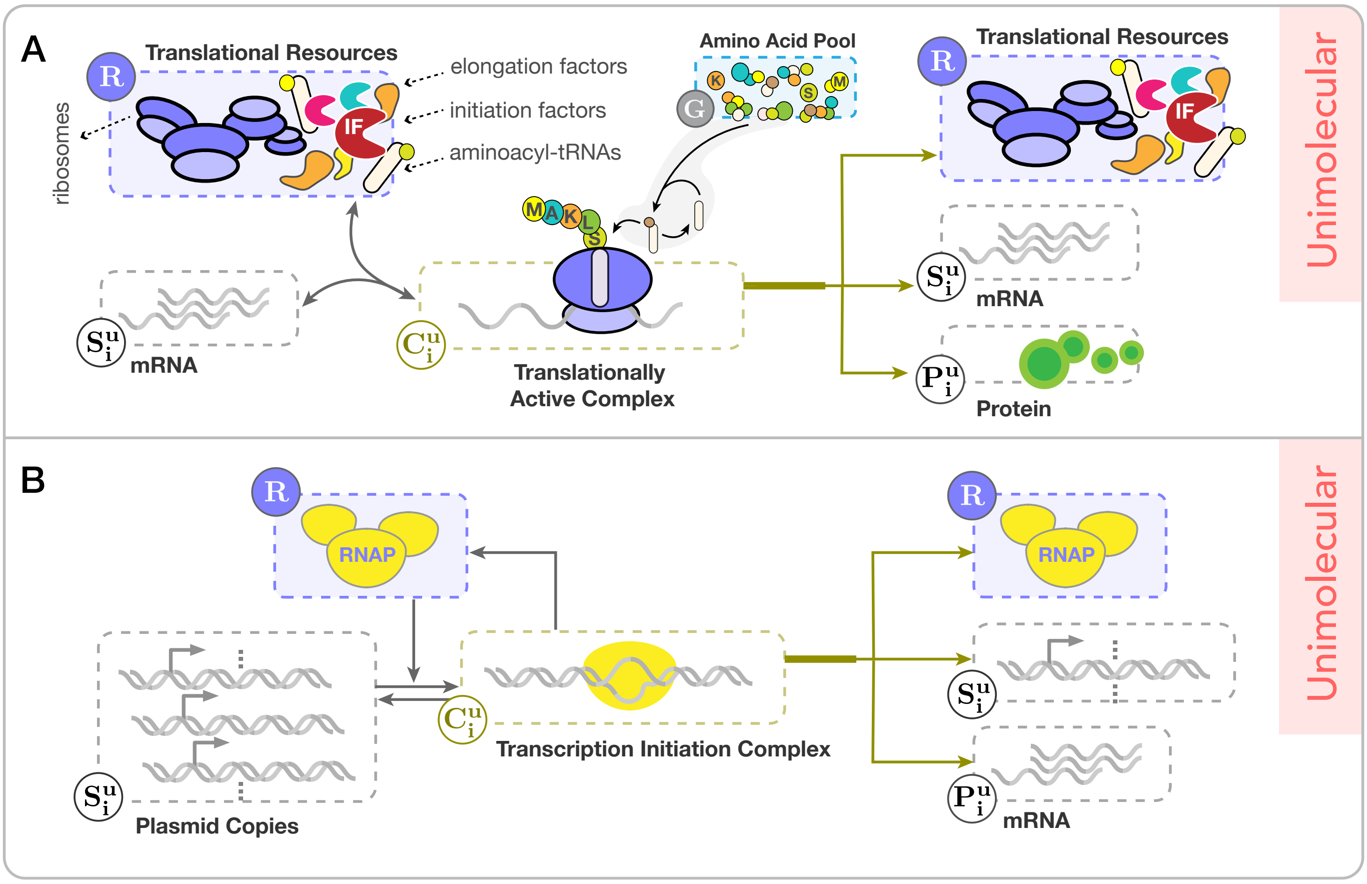

Figure S5: **Examples of resource-limited unimolecular catalytic production reactions.** (**A**) The translation process guided by the availability of specific translational resources, where the complex involving mRNA turns into some polypeptide chain and, then, gradually to a (folded) protein as the end product. This simplified, non-mechanistic model of RNA translation follows the unimolecular type of core resource-limited catalytic reactions. This is based on the assumption that the amino acid pool is plentiful and that removal of the amino acids needed to synthesize the protein from this pool does not result in significant depletion of the pool. (**B**) A transcription process reproduced from Figure S4B with the difference that now the plasmid copies are substituted with the single-copied DNA.

process (Figure S4B). Here, we consider that the availability of substrate DNA might vary over time, as is the case with fluctuating plasmid copy numbers. This means the availability of free plasmids per cell impacts the propensity to form a complex $\mathbf{C}_\mathbf{i}^\mathbf{u}$. For instance, variations in the origin of replications or alterations in the concentration of the Rep protein can result in different numbers of plasmid copies in bacteria. Refer to the example provided in Figure 2 in A Mathematical Framework to Model Intracellular Resource Competition for an illustration of the latter scenario.

#### S2.3 Resource-limited Bimolecular Production Reactions

For a resource-limited bimolecular reaction $\mathcal{R}_i^b$ of the first scenario, we consider the two-step procedure

$$\text{S1}\begin{cases} \text{Step 1:} \quad G_1 + \mathbf{A}_\mathbf{i} + \mathbf{B}_\mathbf{i} \underset{\sigma_i}{\overset{\eta_i}{\rightleftharpoons}} \mathbf{S}_\mathbf{i}^\mathbf{b}, \\ \text{Step 2:} \quad G_2 + \mathbf{S}_\mathbf{i}^\mathbf{b} + \mathbf{R} \underset{k_{r_i}^b}{\overset{k_{f_i}^b}{\rightleftharpoons}} \mathbf{C}_\mathbf{i}^\mathbf{b} \xrightarrow{k_{c_i}^b} \mathbf{S}_\mathbf{i}^\mathbf{b} + \mathbf{R} + \mathbf{P}_\mathbf{i}^\mathbf{b}. \end{cases} \tag{S68}$$

The first step considers the formation of the substrate $\mathbf{S}_\mathbf{i}^\mathbf{b}$ from $\mathbf{A}_\mathbf{i}$ and $\mathbf{B}_\mathbf{i}$. In fact, we have assumed that Step 1 in (S68) takes place without consuming resources and at a faster time scale than Step 2, which is often the case if, for example, $\mathbf{S}_\mathbf{i}^\mathbf{b}$ is the

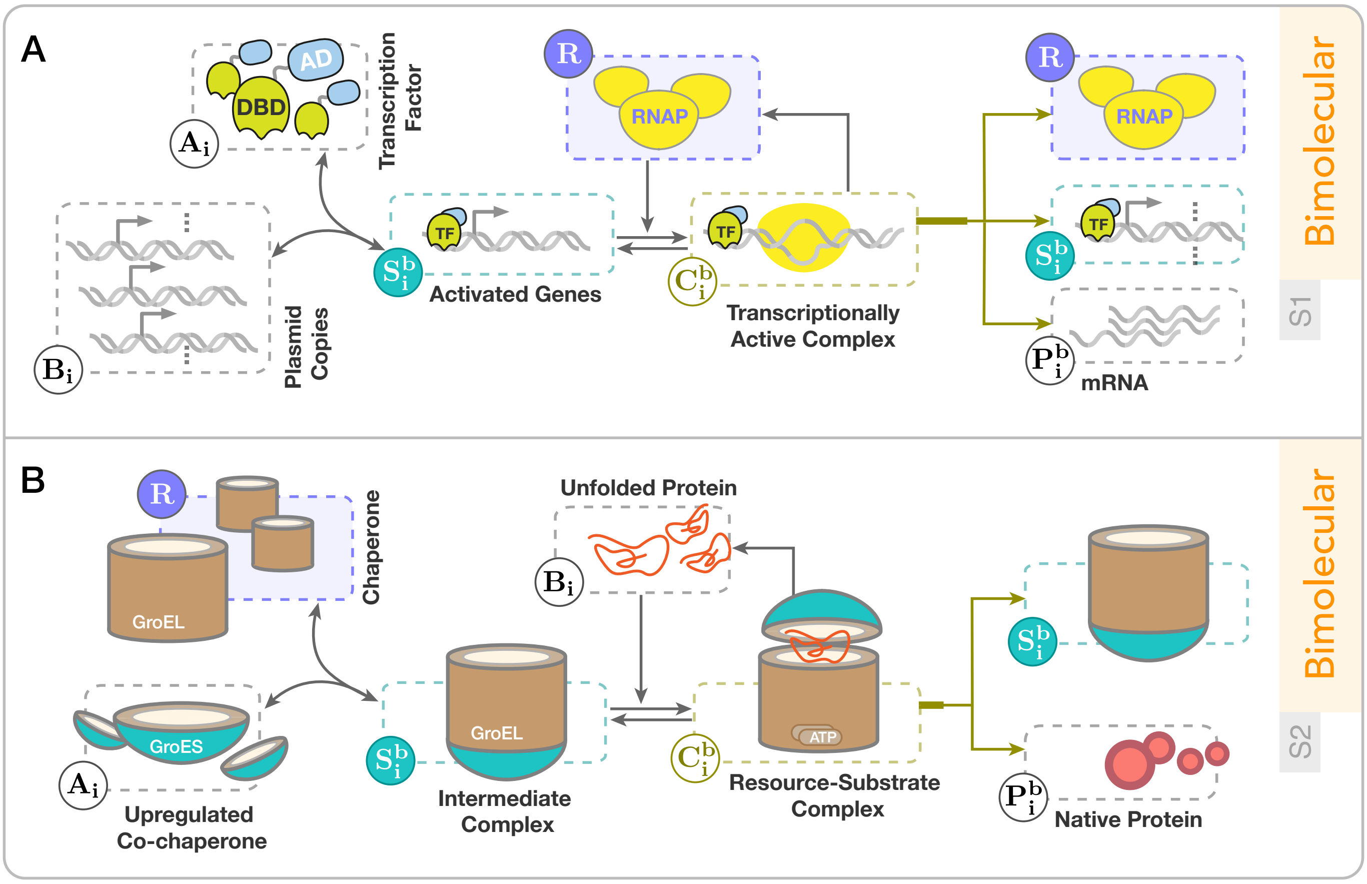

Figure S6: **Different biological examples resembling resource-limited bimolecular production reactions.** (**A**) An extension of the transcription process in Figure S5B to include the role of activating transcription factors ("AD" and "DBD" stand for activation domain and DNA binding domain, respectively). (**B**) A simplified mechanism of chaperone-mediated protein folding in bacteria formulated as a resource-limited bimolecular conversion reaction. By this reaction, an unfolded protein converts to a folded protein as the end product. It is often useful, especially in bioprocesses, to upregulate either of chapreone or co-chaperon to obtain better folding results with the synthetic gene [2, 3].

product of a dimerization/phosphorylation/or ligation process. Similarly, for $\mathcal{R}_i^b$ belonging to the second scenario, we have

$$\text{S2}\begin{cases}\text{Step 1:}\quad \mathrm{G_1} + \mathbf{A_i} + \mathbf{R} \underset{\sigma_i}{\overset{\eta_i}{\rightleftharpoons}} \mathbf{S_i^b}, \\ \text{Step 2:}\quad \mathrm{G_2} + \mathbf{S_i^b} + \mathbf{B_i} \underset{k^b_{r_i}}{\overset{k^b_{f_i}}{\rightleftharpoons}} \mathbf{C_i^b} \xrightarrow{k^b_{c_i}} \mathbf{S_i^b} + \mathbf{B_i} + \mathbf{P_i^b}.\end{cases} \tag{S69}$$

With different rearrangements, the other bimolecular scenarios also exhibit a similar representation. In what follows, we provide four different biological examples of various scenarios. We note that one example from a specific bimolecular scenario may align well with other scenarios as well through minor variations or additional details.

For the first example of resource-limited bimolecular reactions, we consider a more detailed model of gene transcription process, taking into account the role of transcription factors (TFs). Depicted in Figure S6A, we define the protein $\mathbf{A_i}$ to be an activating TF, which once bound to a specific DNA fragment already inserted into the plasmids, facilitates the expression of a target gene by increasing the likelihood of recruiting RNA polymerases (RNAPs). This happens through a reversible binding/unbinding reaction to form the substrate $\mathbf{S_i^b}$, say the product of the first step. If this reaction channel fires fast enough, which is realistic to be the case, then the procedure drawn out in Figure S6A presents a valid example of a bimolecular reaction limited by the presence of shared RNAPs. For this example, one might take nucleotides as both $\mathbf{G_1}$ and $\mathbf{G_2}$.

For the second example, we consider a simplified mechanism of protein folding in bacteria, mediated by chaperones and co-chaperones. One of the substrates, $\mathbf{B_i}$, is the target unfolded or partially folded protein. Chaperones are proteins that bind to the exposed hydrophobic regions of $\mathbf{B_i}$ and prevent it from aggregating, assisting the folding of $\mathbf{B_i}$. We consider chaperones as the resource $\mathbf{R}$. We may think of the co-chaperones, which can help catalyzing this folding process, as the substrate $\mathbf{A_i}$. The bound between chaperon and co-chaperon may be viewed as shaping the intermediate complex $\mathbf{S_i^b}$. After ATP hydrolysis and some conformational changes, the bond between this complex and the unfolded protein $\mathbf{B_i}$ forms a lid-like mechanism that encapsulates the unfolded protein, creating a stable environment for the folding of $\mathbf{B_i}$. This shapes the resource-complex $\mathbf{C_i^b}$, which in turn will gradually convert the unfolded protein into the native polypeptide. We expect the substrate-complex $\mathbf{S_i^b}$ to be recovered as a byproduct of this process. The ATP may be seen as the additional species $\mathbf{G_1}$ and $\mathbf{G_2}$, needed for a successful folding process. This simplified mechanism, as described above, represents the resource-limited bimolecular reaction of the second scenario. However, it is a conversion type since the unfolded protein $\mathbf{B_i}$ does not recycle post-reaction. Schematic illustration is provided in Figure S6B.

A third example is given in Figure S7A, which is the process of ubiquitination of a substrate protein. This process facilitates the degradation of this substrate through cellular endogenous machinery. In this example, a synthetic E3 ubiquitin ligase recognizes and binds to another protein S to first shape the intermediary complex E3S. The protein S is indeed the protein of interest (POI) that is targeted for ubiquitination. Consider the enzyme E3, the protein S, and the resulting complex E3S as the species $\mathbf{A_i}$, $\mathbf{B_i}$, and the substrate $\mathbf{S_i^b}$, respectively. As shown in the figure, the complex E3S recruits the ubiquitin-conjugating enzyme loaded with ubiquitin to form in return another intermediary complex $E2^{Ub}E3S$, called $\mathbf{C_i^b}$. We think of the ubiquitin-loaded E2 enzyme ($E2^{Ub}$) as the limiting shared resources $\mathbf{R}$. Through another reaction, the complex $E2^{Ub}E3S$ undergoes a transformation into three different products. The first product is the ubiquitin-attached S, or $S^{Ub}$, which forms after the ligase E3 has assisted the movement of ubiquitin (Ub) from its carrier (E2) to the target protein S. While the two other products are the free E3 and E2 enzymes released thereupon. The species $\mathbf{S_i^b}$ admits a quasi-steady state proportional to the the product of the original substrates $\mathbf{A_i}$ and $\mathbf{B_i}$. One can easily observe that this example follows the formulation of a resource-limited bimolecular conversion reaction of the third scenario, given in (S5), if we define $\mathbf{P_i^b}$ to be the ubiquitin-attached product ($S^{Ub}$). The ATP is a candidate for the non-rate-limiting abundant species $\mathbf{G_i}$.

Notice that the resources in this example (ubiquitin-attached E2) do not appear directly in the list of products. It is through another intermediary step that the resources recycle, which results in the attachment of the ubiquitins taken up from some other ubiquitin-loaded E1 enzymes to the free (available) E2 enzymes. This intermediary cycle is marked in Figure S7A by shaded-gray color. If this intermediary recycling occurs on a time scale that is faster than or comparable to the production of the complex $\mathbf{C_i^b}$, then it is not necessary to explicitly account for the dynamics of the shaded-gray cycle.

Notice that the third example would be reduced to a resource-limited unimolecular catalytic reaction if we were to consider that the target protein S exists in surplus, meaning that its availability is not rate-limiting within the time interval of interest. In fact, one could think of the enzyme E3, the protein S, and the complex E3S as the substrate $\mathbf{S_i^u}$, the additional substrate $\mathbf{G}$, and some new intermediary species $\mathbf{I}$, respectively. By further considering the ubiquitin-attached protein $S^{Ub}$ as the product species $\mathbf{P_i^u}$ and that the intermediary species $\mathbf{I}$ is at dynamic equilibrium with a quasi-state state value proportional to the current concentration of E3, a unimolecular catalytic formalism would be followed instead.

The fourth example builds upon the previous one, with the primary change being that we now consider the ligase E3 as the

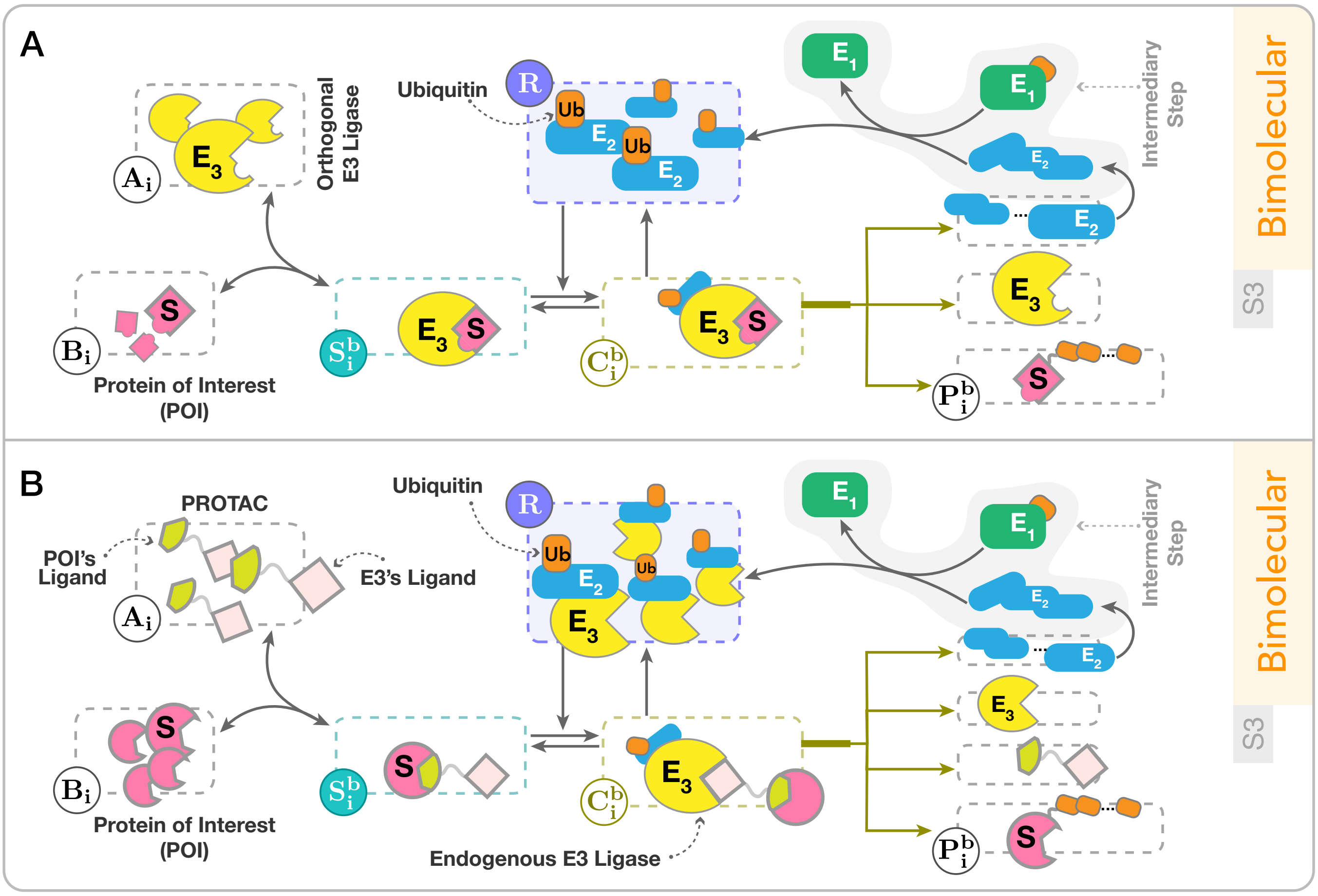

Figure S7: **Examples of resource-limited bimolecular conversion reactions based on the ubiquitylation mechanism.** As a protein post-translational modification mechanism, targeted ubiquitylation facilitates the degradation of a protein of interest (POI) through the cell's endogenous proteseome machinery. (**A**) The mechanism of ubiquitination described as a bimolecular conversion reaction that is limited by the availability of ubiquitin-loaded E2 enzymes. The POI, labeled S, converts to the ubiquitinated protein $S^{Ub}$ as the end product. This example is particularly fitting if an orthogonal pool of E3 ligase that is specific to the POI is synthesized through a synthetically inserted gene. Directing the ligase activity to the POI could be achieved by replacing the native substrate binding domains of the E3 ligase with recombinant modules [4]. (**B**) A targeted protein degrader mechanism based on PROTAC [5] depicted as a bimolecular conversion. The E3 ligase is synthesized through the cell's endogenous machinery, which might be competitively shared between other substrates. The complex $E2^{Ub}E3$ is compactly taken as the shared limiting resources. By assuming that the protein S exists in surplus (grouping it into the species **G**), the both examples would instead resemble a resource-limited unimolecular catalytic prodcution reaction.

cell's endogenous ligase. This necessitates an additional mixed ligand species to enable attachment to the target protein of interest (POI). This species can be proteolysis targeting chimeras, or PROTAC in short. The resulting mechanism provides a targeted protein degradation platform [5]. We have categorized the entire complex, $E2^{Ub}E3$, as the limited resource **R**, which may also be shared between some endogenous genes for their ubiquitination and facilitated degradation. The subsequent details are illustrated in Figure S7B. Note that the ultimate result of this mechanism is the ubiquitination of the POI, which significantly increases the likelihood of its degradation by the endogenous proteasome complexes.

### Supplementary Notes

#### Supplementary Note 1: A Note on the Minimal Realization of Autocatalytic Integral Controllers and the Proposed Layered Version

**Minimal Realization:** Let $\mathbf{Z_1}$ be the only species constituting the biomolecular controller to be designed. Denote the output species to be regulated by $\mathbf{X_n}$. Assume this species is part of a larger network, comprising $n$ distinct species. We shall refer to it as the *process network* of interest. The process species, denoted by $\mathbf{X_i}$, with $i \in \{1, \cdots, n\}$, react with each other through different reaction channels. Think of the controlled process as an uncertain reaction network with two known species $\mathbf{X_1}$ and $\mathbf{X_n}$. It is through these two nodes, referred to as input and output species, that the controller can respectively actuate on and get feedback from the process network, thereby closing the loop (see Figure 3A-B). We consider the following set of chemical reactions

$$\begin{aligned}
\mathbf{Z_1} &\underset{b_{c1}}{\overset{\gamma_{c1}}{\rightleftharpoons}} \varnothing, \\
\mathbf{Z_1} &\xrightarrow{\alpha_1} 2\mathbf{Z_1}, \\
\mathbf{X_n} + \mathbf{Z_1} &\xrightarrow{\theta_1} \mathbf{X_n}, \\
\mathbf{Z_1} &\xrightarrow{k} \mathbf{X_1} + \mathbf{Z_1},
\end{aligned}$$

for the realization of the minimal autocatalytic controller. It can realize, under certain standard assumptions, a constrained IFC mechanism in control of the process network of interest. The assumptions include set-point admissibility, closed-loop stability, and that the basal expression of $\mathbf{Z_1}$ is negligible.

Given mass-action kinetics, an autocatalytic control strategy follows the dynamic equations

$$\begin{cases}
\dot{x} = f(x) + e_1 k z_1, & x_i(0) = x_i^0 \quad \text{(S70a)} \\
\dot{z}_1 = b_{c1} + \alpha_1 z_1 - \theta_1 z_1 x_n - \gamma_{c1} z_1, & z_1(0) = z_1^0 \quad \text{(S70b)}
\end{cases}$$

where the overall deterministic behavior of the process network is represented by the nonlinear function $f : \mathbb{R}^n_{\geq 0} \to \mathbb{R}^n$, $x = [x_1, \cdots, x_n] \in \mathbb{R}^n_{\geq 0}$, and $x_i^0, z_1^0 \in \mathbb{R}_{\geq 0}$ represent the initial conditions. Let us denote the steady-state values by the notation superscript "$*$". In the absence of basal expression rate $b_{c1}$, one can easily observe that at steady-state, the controller dynamics alone can already lead to two different fixed points. One corresponding to $z_1^* = 0$ and the other to $x_n^* = (\alpha_1 - \gamma_{c1})/\theta_1$ (assume that $\alpha_1 > \gamma_{c1}$). Define a logarithmic integral variable $v$ as $v := \ln(z_1)/\theta_1$ to see the underlying integrator. As long as $z_1 \neq 0$, it follows $\mathrm{d}v/\mathrm{d}t = \dot{z}_1/\theta_1 z_1 = (\alpha_1 - \gamma_{c1})/\theta_1 - x_n$, and, hence, $v(t) = c \int_{t_0}^{t} (\alpha_1 - \gamma_{c1})/\theta_1 - x_n(t)$ with $c$ some constant. If it is shown that the solution of $f(x^*) + e_1 k z_1^* = 0$ with $x_n^*$ set to $(\alpha_1 - \gamma_{c1})/\theta_1$ results in a strictly positive $z_1^*$ and non-negative equilibrium concentrations for the rest of network, then $(\alpha_1 - \gamma_{c1})/\theta_1$ will be an admissible set-point. If we further assume that such a positive equilibrium point for (S70) is stable, then, starting from its region of attraction forward in time, RPA will be achieved for the output species $\mathbf{X_n}$. Sample trajectories of this system are shown in Figure 3C.

Note that, in principle, the inhibition of $\mathbf{Z_1}$ by $\mathbf{X_n}$ does not necessarily need to be catalytic. It could involve the degradation of $\mathbf{X_n}$ as well, for example by being a sequestration between $\mathbf{Z_1}$ and $\mathbf{X_n}$, as long as the second-order reaction rate in (S70b) remains a valid approximation. Indeed, our model of $\mathbf{Z_1}$'s inhibition via $\mathbf{X_n}$ in the equation (S70b) represents just one approach to conceptualizing the sensing reaction. Alternative inhibition mechanisms, such as competitive inhibitions with $\mathbf{X_n}$ acting as a transcriptional repressor for $\mathbf{Z_1}$, could also be considered. The resulting dynamic model would then follow, for example, $\dot{z}_1 = \alpha_1 z_1/(1+\theta_1 x_n) - \gamma_{c1} z_1$. This gives rise to alternative versions of the minimal autocatalytic integrator, which may become more relevant given the considered context and available biological components. While our study remains centered on cases of active degradation characterized by (S70b), the variants mentioned, under certain constraints, would likewise enable achieving RPA and support controlled output regulation that remains unaffected by process-side disturbances.

Now, we are in position to incorporate the resource-aware modeling framework introduced in A Mathematical Framework to Model Intracellular Resource Competition to examine the impact of resource-limited interactions on the performance of the minimal autocatalytic IFC motif. We reformulate the dynamic model (S70) within the context of resource competition by substituting all its production reactions with resource-limited counterparts, all of which draw from the same resource pool, $\mathbf{R}$. In the controller side of the model, there is only a single resource-limited reaction, which is of unimolecular type. This reaction hence contributes to the denominator of the functional form for $r(t)$ in (4) by a linear term. The resulting closed-loop dynamic model follows

$$\begin{cases} \dot{x} = f(x,r) + e_1 k^\star z_1 r, & \text{(S71a)} \\ \dot{z}_1 = \alpha_1^\star z_1 r - \theta_1 z_1 x_n - \gamma_{c1} z_1, & \text{(S71b)} \end{cases}$$

where

$$r = R_{tot}/(1 + w_{p1}x + x^{\mathrm{T}} w_{p2} x + w_1 z_1)$$

and the constants $w_1 > 0$, $w_{p1} \in \mathbb{R}^n_{\geq 0}$ and $w_{p2} \in \mathbb{R}^{n\times n}_{\geq 0}$ represent the competition gains resulting from the competition for $\mathbf{R}$.

**Layered Version:** Similarly for this biocontroller, we first consider resource-unlimited settings. Let us add the auxiliary controller species $\mathbf{Z_2}$ to the network and introduce the reactions associated to it.

$$\mathbf{Z_2} + \mathbf{X_n} \xrightarrow{\alpha_2} \mathbf{X_n} + 2\mathbf{Z_2}$$
$$\mathbf{Z_2} \underset{b_{c2}}{\overset{\gamma_{c2}}{\rightleftarrows}} \varnothing.$$

In these reactions, $\mathbf{X_n}$ catalyzes the production of $\mathbf{Z_2}$ while $\mathbf{Z_2}$ is degraded (or diluted) at a constant rate. The reaction network realizing the proposed controller then becomes

$$\mathcal{R}_{c1} : \mathbf{Z_1} \underset{b_{c2}}{\overset{\gamma_{c1}}{\rightleftarrows}} \varnothing,$$
$$\mathcal{R}_{c2} : \mathbf{Z_1} \xrightarrow{\alpha_1} 2\mathbf{Z_1},$$
$$\mathcal{R}_{c3} : \mathbf{X_n} + \mathbf{Z_1} \xrightarrow{\theta_1} \mathbf{X_n},$$
$$\mathcal{R}_{c4} : \mathbf{Z_2} + \mathbf{X_n} \xrightarrow{\alpha_2} \mathbf{X_n} + 2\mathbf{Z_2},$$
$$\mathcal{R}_{c5} : \mathbf{Z_2} \underset{b_{c2}}{\overset{\gamma_{c2}}{\rightleftarrows}} \varnothing,$$
$$\mathcal{R}_{c6} : \mathbf{Z_1} \xrightarrow{k} \mathbf{X_1} + \mathbf{Z_1}.$$

Under mass-action kinetics, the time evolution of the concentrations of the controller species in the network is given by

$$\dot{z}_1 = b_{c1} + \alpha_1 z_1 - \theta_1 z_1 x_n - \gamma_{c1} z_1,$$
$$\dot{z}_2 = b_{c2} + \alpha_2 z_2 x_n - \gamma_{c2} z_2.$$

Compared to the minimal autocatalytic integral feedback in a non-competing scenario (see above), our newly introduced autocatalytic controller does not offer any improvements. In fact, the controller may not even allow stable regulation without the extinction of either $z_1$ or $z_2$ at steady state. Let us incorporate our resource-aware framework from A Mathematical Framework to Model Intracellular Resource Competition to introduce resource competition among the two controller species (as well as process species) for shared resources $\mathbf{R}$. We assume that the production rates of these two species hinge on the availability of the resources. The closed-loop dynamic model would then take the form of:

$$\begin{cases} \dot{x} = f(x, r) + e_1 k^\star z_1 r, & \text{(S72a)} \\ \dot{z}_1 = \alpha_1^\star z_1 r - \theta_1 z_1 x_n - \gamma_{c1} z_1, & \text{(S72b)} \\ \dot{z}_2 = \alpha_2^\star z_2 x_n r - \gamma_{c2} z_2, & \text{(S72c)} \end{cases}$$

with

$$r = R_{tot}/(1 + w_{p1}x + x^{\mathrm{T}} w_{p2} x + w_1 z_1 + w_2 z_2 x_n).$$

Without loss of generality, we have assumed that the bimolecular reaction $\mathcal{R}_{c4}$ is a resource-limited production reaction of the first scenario (see the respective illustration in Figure 1 or the representation S1 in Section S1.1). Note that in the resource-aware model presented in (S72), all production reactions are taken to be resource limited, with the assumption that the basal expressions of the controller species are negligible compared to other production/degradation rates that dictate the dynamics of $\mathbf{Z_1}$ and $\mathbf{Z_2}$ ($b_{c1} = 0$ and $b_{c2} = 0$). Similar to process parameters, we treat the competition gains $w_{p1} \in \mathbb{R}^n_{\geq 0}$, $w_{p2} \in \mathbb{R}^{n\times n}_{\geq 0}$, $w_1, w_2 \in \mathbb{R}_{>0}$ and $R_{tot}$ as unknown quantities which may neither be precisely measurable nor tunable. There exist two intertwined integral feedback loops working in tandem. One can define the logarithmic integral variables $v_1 := \ln(z_1)/\theta_1$ and $v_2 := \ln(z_2)/\alpha_2^\star$ to see this. Given the closed-loop stability, one integral loop steers the variable $r(t)x_n(t)$ towards $\gamma_{c2}/\alpha_2^\star$, the other regulates $x_n(t)$ at $(\alpha_1^\star r(t) - \gamma_{c1})/\theta_1$ simultaneously. As mentioned earlier, the inhibitory sensing reaction from $\mathbf{X_n}$ to $\mathbf{Z_1}$ does not necessarily need to be through active degradation. Indeed, a competitive transcriptional repression dynamically modeled as $\dot{z}_1 = \alpha_1^\star z_1 r/(1 + \theta_1 x_n) - \gamma_{c1} z_1$ would essentially lead to similar results.

#### Supplementary Note 2: A Systematic Search for Autocatalytic Integral Feedback Mechanisms Capable of Robust Perfect Adaptation in Competitive Scenarios

This section explores various modifications within a class of autocatalytic biomolecular integral feedback motifs. Specifically, we conduct a systematic search across eight distinct variations of the core autocatalytic controller that represents the minimal realization, involving the species $\mathbf{Z_1}$ and its interactions with $\mathbf{X_1}$ and $\mathbf{X_n}$. For reactions from $\mathbf{X_n}$ to $\mathbf{Z_1}$, we always consider the effect of $\mathbf{Z_1}$ as a substrate limiting the associated reactions rates, although we may drop explicit drawings in the schematics that indicate this influence to avoid clutter. This way, we ensure that the dynamical modeling of the controller species includes all autocatalytic terms. For the inhibitory reactions from $\mathbf{X_n}$ to $\mathbf{Z_1}$, we consider active degradation or sequestration types, both in resource-limited and resource-unlimited scenarios. Additionally, we examine three different buffering mechanisms for each of the considered eight categories. The *passive* buffering mechanism entails regulating the free resources $\mathbf{R}$ at a fixed, preset value, regardless of the output levels. In contrast, *active* buffering adjusts the resource buffering corresponding to the equilibrium concentration of the output species, $\mathbf{X_n}$, thereby aligning resource availability with the needs of the output species.

The schematic diagrams of all 24 variants are presented in Figure S8. Notably, some variants failed to maintain the robust perfect adaptation (RPA) property, due to issues such as the structural absence of feasible equilibria in the positive orthant. We report in Tables S2 and S3 the variants that demonstrated RPA. These tables present their dynamic models and steady-state relationships, enabling us to differentiate between controller motifs that sustain RPA under any controller parameter settings and those limited to specific parametric conditions (referred to as "constrained" controllers). While our preference leans towards the former category for obvious reasons, constrained controllers may still be advantageous for practical implementation. That being said, a design with restrictions may sometimes be preferred over one that is mathematically less constrained, given the context and biological limitations that arise when considering the implementation of a candidate controller topology. Through our systematic analysis, we found that 16 out of the 24 variants maintained the RPA property. Among these, only four were able to achieve this property unconditionally and 14 required at least one type of resource-limited bimolecular reactions to take place, whether involving catalytic production, conversion, or sequestration types. The variant 1.3 has been selected as our primary core motif, and it is the main focus of our discussion in the main manuscript.

#### Supplementary Note 3: Higher-order Implementations of the Proposed Layered Autocatalytic Integral Feedback Controller

Here, we discuss a class of higher-dimensional systems that can be structurally reduced to the layered autocatalytic integral feedback controller considered in Multi-layer Autocatalytic Biomolecular Controllers: An Effective Solution. We provide assumptions under which this model-order reduction holds and will consider an example of it before concluding this note. Consider the following closed-loop dynamical system whose controller consists of two subsystems: $\mathcal{Y}$ and $\mathcal{Z}$.

$$
\begin{cases}
\dot{x} = f_x(x, r_1, \cdots, r_m) \overbrace{+e_1 k^\star u(z_1, z_2, r_1, \cdots, r_m)}^{\text{actuation term (control input)}}, & \text{(S73a)}\\
\epsilon \dot{y} = f_y(x, y, z_1, z_2, r_1, \cdots, r_m), & \text{(S73b)}\\
\dot{z}_1 = B_1 y h_1(r_1, \cdots, r_m) - \theta_1 z_1 x_n - \gamma_{c1} z_1, & \text{(S73c)}\\
\dot{z}_2^c = \dfrac{B_2^c}{\epsilon_2^c} y h_2^c(r_1, \cdots, r_m) - \theta_2 z_2^c x_n - \dfrac{\gamma_{c2}^c}{\epsilon_2^c} z_2^c, & \text{(S73d)}\\
\dot{z}_2 = \theta_2 z_2^c x_n - \gamma_{c2} z_2. & \text{(S73e)}
\end{cases}
$$

The state variables $x \in \mathbb{R}^n_{\geq 0}$ associate to the process network under control. The vector $y \in \mathbb{R}^q_{\geq 0}$ is a column vector representing the state variables of the subsystem $\mathcal{Y}$. $B_1$ and $B_2^c$ are row vectors of the same size as $y$. The variables $r_i$, $i \in \{1, \cdots, m\}$, account for indirect couplings which might emerge, for example, from sharing resources in the system. Let us hold the following assumptions on this system, also referred to as the full system, over a finite-time interval $[t_0, t_1]$.

**Assumptions.** *Within the time interval $t \in [t_0, t_1]$ and for a given set of parameters, the full system (S73) fulfills the following:*

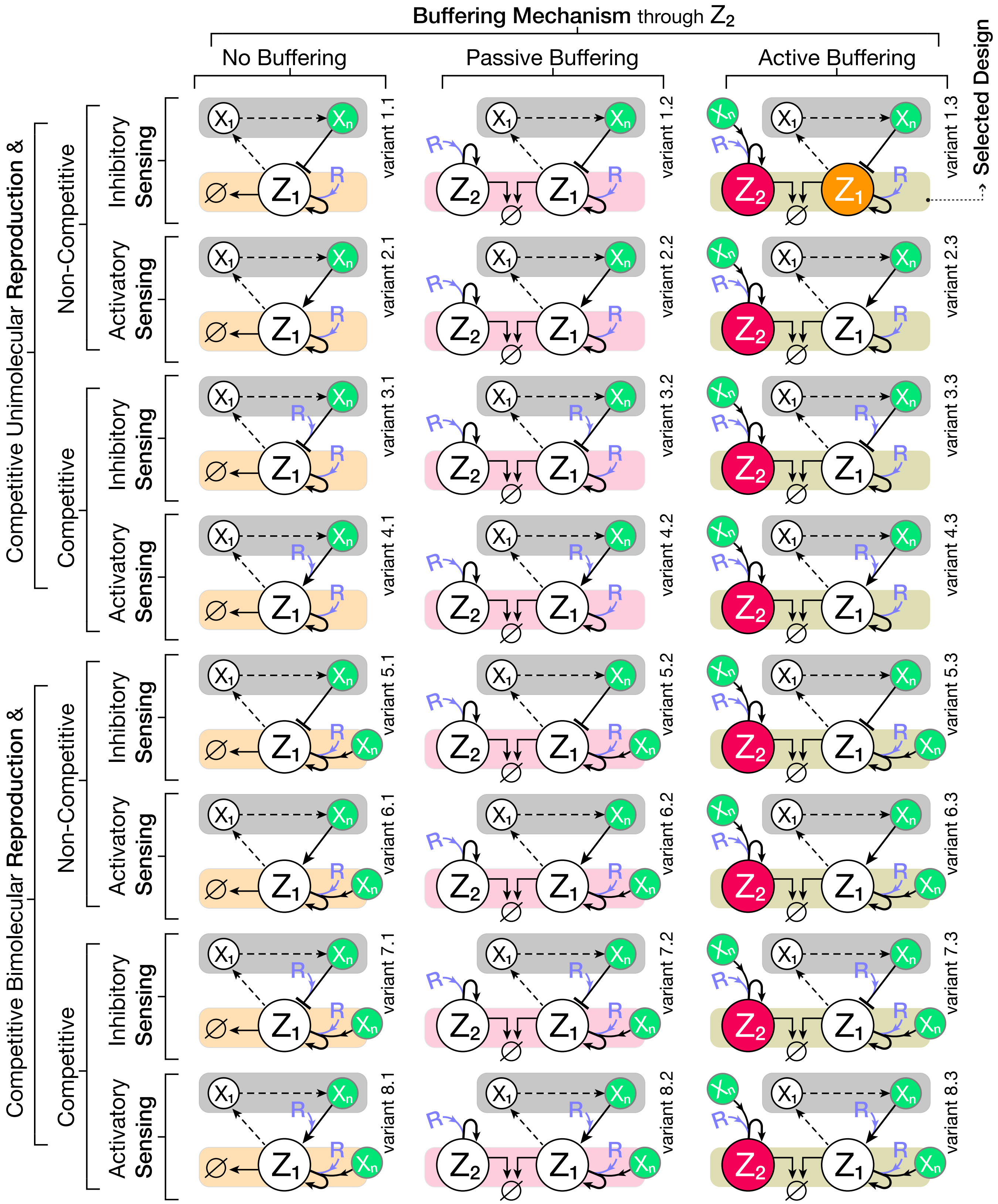

Figure S8: **Exploring variants of the autocatalytic feedback mechanism and their layered versions for robust perfect adaptation (RPA) under resource constraints.** Each row features an identical circuit for, and connections between, $\mathbf{Z_1}$ and the process. The self-replication of $\mathbf{Z_1}$ is assumed to be limited by the availability of some finite, shared resources, denoted by $\mathbf{R}$. It is through the known nodes of contact $\mathbf{X_1}$ and $\mathbf{X_n}$ that the controller can compute and apply the control signal to the process. Within the same row, the distinction in each column arises from the consideration of three distinct buffering mechanisms via introducing a new controller species $\mathbf{Z_2}$. The first column has no buffering species, while the second and third ones have this buffering mechanism in "passive" and "active" manners, respectively. For the sensing reaction from $\mathbf{X_n}$ to $\mathbf{Z_1}$, we consider four scenarios: it can influence $\mathbf{Z_1}$'s dynamics either in an inhibitory or activatory manner, and this can be carried out in a resource-limited (drawing from $\mathbf{R}$) or a resource-agnostic manner.

| | Variant | Constrained? | Controller Dynamic Model | Steady-State Relations |
|---|---|---|---|---|
| | 3.1 | Yes | $\dot{z}_1 = \alpha_1^\star z_1 r - \theta_1^\star z_1 x_n r - \gamma_{c1} z_1$ | $x_n^* \approx \alpha_1^\star/\theta_1^\star \quad (\gamma_{c1} \ll \alpha_1 r^*)$<br>$r^*$ not regulated |
| | 6.1 | Yes | $\dot{z}_1 = \alpha_1^\star z_1 x_n r + \theta_1 z_1 x_n - \gamma_{c1} z_1$ | $x_n^* \approx \gamma_{c1}/\theta_1 \quad (\alpha_1 r^* \ll \theta_1)$<br>$r^*$ not regulated |
| Passive Buffering Mechanism | 1.2 | Yes | $\dot{z}_1 = \alpha_1^\star z_1 r - \theta_1 z_1 x_n - \gamma_{c1} z_1$<br>$\dot{z}_2 = \alpha_2^\star z_2 r - \gamma_{c2} z_2$ | $x_n^* = (\alpha_1^\star \gamma_{c2}/\alpha_2^\star - \gamma_{c1})/\theta_1$<br>$r^* = \gamma_{c2}/\alpha_2^\star$ |
| Passive Buffering Mechanism | 2.2 | Yes | $\dot{z}_1 = \alpha_1^\star z_1 r + \theta_1 z_1 x_n - \gamma_{c1} z_1$<br>$\dot{z}_2 = \alpha_2^\star z_2 r - \gamma_{c2} z_2$ | $x_n^* = (\gamma_{c1} - \alpha_1^\star \gamma_{c2}/\alpha_2^\star)/\theta_1$<br>$r^* = \gamma_{c2}/\alpha_2^\star$ |
| Passive Buffering Mechanism | 3.2 | Yes | $\dot{z}_1 = \alpha_1^\star z_1 r - \theta_1^\star z_1 x_n r - \gamma_{c1} z_1$<br>$\dot{z}_2 = \alpha_2^\star z_2 r - \gamma_{c2} z_2$ | $x_n^* = (\alpha_1^\star - \alpha_2^\star \gamma_{c1}/\gamma_{c2})/\theta_1^\star$<br>$r^* = \gamma_{c2}/\alpha_2^\star$ |
| Passive Buffering Mechanism | 4.2 | Yes | $\dot{z}_1 = \alpha_1^\star z_1 r + \theta_1^\star z_1 x_n r - \gamma_{c1} z_1$<br>$\dot{z}_2 = \alpha_2^\star z_2 r - \gamma_{c2} z_2$ | $x_n^* = (\alpha_2^\star \gamma_{c1}/\gamma_{c2} - \alpha_1^\star)/\theta_1^\star$<br>$r^* = \gamma_{c2}/\alpha_2^\star$ |
| Passive Buffering Mechanism | 5.2 | Yes | $\dot{z}_1 = \alpha_1^\star z_1 x_n r - \theta_1 z_1 x_n - \gamma_{c1} z_1$<br>$\dot{z}_2 = \alpha_2^\star z_2 r - \gamma_{c2} z_2$ | $x_n^* = \gamma_{c1}/(\alpha_1^\star \gamma_{c2}/\alpha_2^\star - \theta_1)$<br>$r^* = \gamma_{c2}/\alpha_2^\star$ |
| Passive Buffering Mechanism | 6.2 | No | $\dot{z}_1 = \alpha_1^\star z_1 x_n r + \theta_1 z_1 x_n - \gamma_{c1} z_1$<br>$\dot{z}_2 = \alpha_2^\star z_2 r - \gamma_{c2} z_2$ | $x_n^* = \gamma_{c1}/(\alpha_1^\star \gamma_{c2}/\alpha_2^\star + \theta_1)$<br>$r^* = \gamma_{c2}/\alpha_2^\star$ |
| Passive Buffering Mechanism | 7.2 | Yes | $\dot{z}_1 = \alpha_1^\star z_1 x_n r - \theta_1^\star z_1 x_n r - \gamma_{c1} z_1$<br>$\dot{z}_2 = \alpha_2^\star z_2 r - \gamma_{c2} z_2$ | $x_n^* = \gamma_{c1}\alpha_2^\star/\gamma_{c2}(\alpha_1^\star - \theta_1^\star)$<br>$r^* = \gamma_{c2}/\alpha_2^\star$ |
| Passive Buffering Mechanism | 8.2 | No | $\dot{z}_1 = \alpha_1^\star z_1 x_n r + \theta_1^\star z_1 x_n r - \gamma_{c1} z_1$<br>$\dot{z}_2 = \alpha_2^\star z_2 r - \gamma_{c2} z_2$ | $x_n^* = \gamma_{c1}\alpha_2^\star/\gamma_{c2}(\alpha_1^\star + \theta_1^\star)$<br>$r^* = \gamma_{c2}/\alpha_2^\star$ |

Table S2: **Dynamic model and steady-state relationships pertaining to those considered variants of autocatalytic and layered autocatalytic IFC mechanisms that have the potential to achieve RPA at the output level with respect to a wide range of perturbations influencing the process network.** Variant numbers match with the corresponding circuit diagrams in Figure S8. The variants not listed in the table may not be able to hold the RPA property or may result in an infeasible (always negative) equilibrium point. The steady-state equations in the last column pertain to only the desired equilibrium point, in which neither $\mathbf{Z_1}$ nor $\mathbf{Z_2}$ is zero (implying that the initial conditions of these two species are also bounded away from their absorbing states at zero). In the second column we report if the considered variant is able to achieve RPA without the need of satisfying any controller-related parametric condition, though still the constraints arising from the process side and the closed-loop stability conditions have to be taken into account. In this column, a "No" means that the variant works for every select of controller parameters, thus it is NOT constrained. On the opposite side, a "Yes" means that the considered variation of the controller has to fulfill certain necessary parametric condition(s) in order to work. Ideally, we would prefer to have non-constrained controllers, though the constrained ones also may find application in certain circuit design paradigms.

| | Variant | Constrained? | Controller Dynamic Model | Steady-State Relations |
|---|---|---|---|---|
| Active Buffering Mechanism | 1.3 | No | $\dot{z}_1 = \alpha_1^\star z_1 r - \theta_1 z_1 x_n - \gamma_{c1} z_1$<br>$\dot{z}_2 = \alpha_2^\star z_2 x_n r - \gamma_{c2} z_2$ | $x_n^* = \sqrt{(\gamma_{c1}/2\theta_1)^2 + \alpha_1^\star \gamma_{c2}/\alpha_2^\star \theta_1} - \gamma_{c1}/2\theta_1$<br>$r^* = \gamma_{c2}/\alpha_2^\star x_n^*$ |
| | 2.3 | Yes | $\dot{z}_1 = \alpha_1^\star z_1 r + \theta_1 z_1 x_n - \gamma_{c1} z_1$<br>$\dot{z}_2 = \alpha_2^\star z_2 x_n r - \gamma_{c2} z_2$ | $x_n^* = \gamma_{c1}/2\theta_1 \pm \sqrt{(\gamma_{c1}/2\theta_1)^2 - \alpha_1^\star \gamma_{c2}/\alpha_2^\star \theta_1}$<br>$r^* = \gamma_{c2}/\alpha_2^\star x_n^*$ if $\gamma_{c1}/2\theta_1 \geq \sqrt{\alpha_1^\star \gamma_{c2}/\alpha_2^\star \theta_1}$ |
| | 3.3 | No | $\dot{z}_1 = \alpha_1^\star z_1 r - \theta_1^\star z_1 x_n r - \gamma_{c1} z_1$<br>$\dot{z}_2 = \alpha_2^\star z_2 x_n r - \gamma_{c2} z_2$ | $x_n^* = \alpha_1^\star/(\theta_1^\star + \gamma_{c1}\alpha_2^\star/\gamma_{c2})$<br>$r^* = \gamma_{c2}/\alpha_2^\star x_n^*$ |
| | 4.3 | Yes | $\dot{z}_1 = \alpha_1^\star z_1 r + \theta_1^\star z_1 x_n r - \gamma_{c1} z_1$<br>$\dot{z}_2 = \alpha_2^\star z_2 x_n r - \gamma_{c2} z_2$ | $x_n^* = \alpha_1^\star/(\gamma_{c1}\alpha_2^\star/\gamma_{c2} - \theta_1^\star)$<br>$r^* = \gamma_{c2}/\alpha_2^\star x_n^*$ |
| | 5.3 | Yes | $\dot{z}_1 = \alpha_1^\star z_1 x_n r - \theta_1 z_1 x_n - \gamma_{c1} z_1$<br>$\dot{z}_2 = \alpha_2^\star z_2 x_n r - \gamma_{c2} z_2$ | $x_n^* = (\alpha_1^\star \gamma_{c2}/\alpha_2^\star - \gamma_{c1})/\theta_1$<br>$r^* = \gamma_{c2}/\alpha_2^\star x_n^*$ |
| | 6.3 | Yes | $\dot{z}_1 = \alpha_1^\star z_1 x_n r + \theta_1 z_1 x_n - \gamma_{c1} z_1$<br>$\dot{z}_2 = \alpha_2^\star z_2 x_n r - \gamma_{c2} z_2$ | $x_n^* = (\gamma_{c1} - \alpha_1^\star \gamma_{c2}/\alpha_2^\star)/\theta_1$<br>$r^* = \gamma_{c2}/\alpha_2^\star x_n^*$ |

Table S3: **Continuation of Table S2 for the cases of active buffering mechanism.**

A3.1 *The boundary layer associated to the fast subsystem $\mathcal{Y}$ is exponentially stable, meaning that there exists a slow manifold $y = \mathscr{H}(x, z_1, z_2, r_1, \cdots, r_m)$ in the positive orthant such that the state variables $y$ obtained from (S73) approach to it arbitrarily fast in the limit $\epsilon \to 0$.*

A3.2 *Given the dynamic model $f_y$, the vectors $B_1$ and $B_2^c$ are chosen in such a way that the resulting quasi-steady state value corresponding to the terms $B_1 y h_1(r_1, \cdots, r_m)$ and $B_2^c y h_2^c(r_1, \cdots, r_m)$ are linearly dependent on $z_1$ and $z_2$ multiplied by the same function $R(r_1, \cdots, r_m)$, respectively. That is, there exist positive constants $\Theta_1$ and $\Theta_2^c$ such that*

$$B_1 \mathscr{H}(x, z_1, z_2, r_1, \cdots, r_m) \times h_1(r_1, \cdots, r_m) = \Theta_1 z_1 R(r_1, \cdots, r_m)$$

*and*

$$B_2^c \mathscr{H}(x, z_1, z_2, r_1, \cdots, r_m) \times h_2^c(r_1, \cdots, r_m) = \Theta_2^c z_2 R(r_1, \cdots, r_m).$$

A3.3 *The asymptotic inequality $\gamma_{c2}^c \gg \theta_2 x_n$ holds at steady state.*

A3.4 *The function $h_2^c(r_1, \cdots, r_m)$ is independent of $z_2^c$, meaning $\partial h_2^c/\partial z_2^c = 0$.*

Note that, in the limit $\epsilon_2^c \to 0$, we face a singular perturbation (SP) problem. It is a regular SP problem as the boundary layer associated with it is non-singular throughout the state space (thanks to the assumption A3.4). Therefore, by referring to [6, Theorem 11.1] and [7, Theorem 3.1], the following statement can be concluded over any finite-time interval $t \in [t_0, t_1]$: Fix every parameter except $\epsilon_2^c$ and $\epsilon$. Then, there exist sufficiently small positive $\epsilon_2^{c*}$ and $\epsilon^*$ such that for every $0 < \epsilon_2^c \leq \epsilon_2^{c*}$ and

$0 < \epsilon \leq \epsilon^*$, the dynamic model in (S73) evolves within a tube of $\mathcal{O}(\epsilon_2^c) + \mathcal{O}(\epsilon)$ about the corresponding trajectory obtained from the reduced system

$$\begin{cases} \dot{x} = f_x(x, r_1, \cdots, r_m) + e_1 k^\star u(z_1, z_2, r_1, \cdots, r_m), & \text{(S74a)} \\ \dot{z}_1 = \Theta_1 z_1 R(r_1, \cdots, r_m) - \theta_1 z_1 x_n - \gamma_{c1} z_1, & \text{(S74b)} \\ \dot{z}_2 = \dfrac{\theta_2 \Theta_2^c}{\gamma_{c2}^c} z_2 R(r_1, \cdots, r_m) x_n - \gamma_{c2} z_2, & \text{(S74c)} \end{cases}$$

for the same initial conditions. This statement generalizes to infinite-time horizons if the reduced system admits a feasible equilibrium point, such a fixed point is exponentially stable, and if the initial conditions are picked from its region of attraction [6, Theorem 11.2]. The above reduced-order model follows the same dynamical representation as the layered autocatalytic controller motif considered in the main text, Multi-layer Autocatalytic Biomolecular Controllers: An Effective Solution, if we define: $r = R(r_1, \cdots, r_m)$, $\alpha_1^\star := \Theta_1$ and $\alpha_2^\star := \theta_2 \Theta_2^c / \gamma_{c2}^c$.

Let us revisit the example control circuit considered in Multi-layer Autocatalytic Biomolecular Controllers: An Effective Solution, Figure 4B. This circuit suggests a biological realization of the proposed layered autocatalytic controller. Let us group the transcriptional and translational cellular resources shared between the controller genes into two separate resource pools, represented by the species $\mathbf{R_1}$ and $\mathbf{R_2}$, respectively. Taking into account both transcriptional and translational resource competition, we write the mathematical model describing the dynamical behavior of the closed-loop as follows:

$$\begin{cases} \dot{x} = f_x(x, r_1, r_2) + e_1 k^\star u(z_1, r_1, r_2), & \text{(S75a)} \\ \epsilon \dot{y}_1 = \kappa_{y_1} z_1 r_1 - \gamma_{y_1} y_1, & \text{(S75b)} \\ \dot{z}_1 = \kappa_{z_1} y_1 r_2 - \theta_1 z_1 x_n - \gamma_{c1} z_1, & \text{(S75c)} \\ \epsilon \dot{y}_2 = \kappa_{y_2} z_2 r_1 - \gamma_{y_2} y_2, & \text{(S75d)} \\ \dot{z}_2^c = \kappa_{z_2^c} \dfrac{[0 \ 1]}{\epsilon_2^c} y r_2 - \theta_2 z_2^c x_n - \dfrac{\gamma_{c2}^c}{\epsilon_2^c} z_2^c, & \text{(S75e)} \\ \dot{z}_2 = \theta_2 z_2^c x_n - \gamma_{c2} z_2, & \text{(S75f)} \end{cases}$$

with

$$r_1 = R_{tot1} / (1 + w_{p11} x + x^{\mathrm{T}} w_{p12} x + w_{11} z_1 + w_{12} z_2),$$

and

$$r_2 = R_{tot2} / (1 + w_{p21} x + x^{\mathrm{T}} w_{p22} x + w_{21} y_1 + w_{22} y_2).$$

Unlike the gene expression control circuit provided in Figure 4, where the process network comprises a single species, the above model considers a general process network. The controller components, however, follow the same biological example as in Figure 4. The controller comprises the proteins $\mathbf{Z_1}$, $\mathbf{Z_2}$, and $\mathbf{Z_2^c}$. Additionally, certain mRNA species are included as controller species. The mRNA strand associated with $\mathbf{Z_2^c}$ is denoted by $\mathbf{Y_2}$. The species $\mathbf{Y_1}$ represents the mRNA strand corresponding to $\mathbf{Z_1}$. Refer to Figure 4B for illustrations. Here, the vector $y$ is given by $y = [y_1 \ y_2]^{\mathrm{T}}$. The species $\mathbf{X_n}$ actively degrades $\mathbf{Z_1}$ by means of an encapsulated protease. This is achieved through a degradation tag fused to $\mathbf{Z_1}$, which recognizes the protease and thereby facilitates the degradation of $\mathbf{Z_1}$. Similarly, $\mathbf{X_n}$ degrades $\mathbf{Z_2^c}$ by cleaving it from a specific domain. This cleavage releases the cargo active complex $\mathbf{Z_2}$ caged within the complex $\mathbf{Z_2^c}$. This is made possible because the caging sequence is placed after the cleavage site; thus, cleaving $\mathbf{Z_2^c}$ by the protease in $\mathbf{X_n}$ deactivates the caging system. The cargo $\mathbf{Z_2}$ functions as a transcription factor (TF) for $\mathbf{Z_2^c}$'s gene upon release. Note that $\mathbf{Z_2^c}$ does not autoregulate its expression, thanks to the caging system involved. Additionally, a secondary degradation tag is inserted after the cleavage site of $\mathbf{Z_2^c}$ that facilitates its degradation through an orthogonal protease, without degrading the released $\mathbf{Z_2}$s. This provides additional flexibility to separately modulate the degradation rates $\gamma_{c2}^c$ and $\gamma_{c2}$, serving as an extra tuning knob for fulfilling the assumption A3.3. Alternatively, one may consider lowering the degradation rate $\theta_2$ to meet A3.3.

As can be seen, this example control circuit follows the same dynamic model as the full system above in (S73), if we take $m = 2$, $h_1 := r_2$, $h_2^c := r_2$, $B_1 := [\kappa_{z_1}\ 0]$, and $B_2^c := [0\ \kappa_{z_2^c}]$. It is straightforward to check that the assumption A3.1 is met if $\epsilon$ is chosen sufficiently small. By this assumption, we take that the transcriptional components are at dynamic equilibrium. By defining $R(r_1, r_2) := r_1 r_2$, the assumption A3.2 is also met as the quasi-steady states $y = \mathscr{H}$, with $\mathscr{H}$ given by $\mathscr{H} := [\kappa_{y_1} z_1 r_1/\gamma_{y_1}\ \kappa_{y_2} z_2 r_1/\gamma_{y_2}]^{\mathrm{T}}$, are linearly dependent on $z_1 R(r_1, r_2)$ and $z_2 R(r_1, r_2)$. The assumption A3.4 is already fulfilled as $r_2$, and hence $h_2^c$, is independent of $z_2^c$. Therefore, if $\epsilon_c^2$ is chosen sufficiently small and if $\gamma_{c2}^c$ is large enough to satisfy $\gamma_{c2}^c \gg \theta_2 x_n^*$, the full system (S75) reduces to the following closed-loop system:

$$\begin{cases} \dot{x} = f(x, r_1, r_2) + e_1 k^\star u(z_1, r_1, r_2), & \text{(S76a)} \\ \dot{z}_1 = \kappa_{z_1} \dfrac{\kappa_{y_1}}{\gamma_{y_1}} z_1 r_1 r_2 - \theta_1 z_1 x_n - \gamma_{c1} z_1, & \text{(S76b)} \\ \dot{z}_2 = \theta_2 \dfrac{\kappa_{z_2^c} \kappa_{y_2}}{\gamma_{c2}^c \gamma_{y_2}} z_2 x_n r_1 r_2 - \gamma_{c2} z_2, & \text{(S76c)} \end{cases}$$

wherein

$$r_1 = R_{tot1}/(1 + w_{p11} x + x^{\mathrm{T}} w_{p12} x + w_{11} z_1 + w_{12} z_2),$$

and

$$r_2 = R_{tot2}/(1 + w_{p21} x + x^{\mathrm{T}} w_{p22} x + w_{21} \frac{\kappa_{y_1}}{\gamma_{y_1}} z_1 r_1 + w_{22} \frac{\kappa_{y_2}}{\gamma_{y_2}} z_2 r_1).$$

It is now an easy task to check that by defining $r := r_1 r_2$, $\alpha_1^\star := \kappa_{z_1} \frac{\kappa_{y_1}}{\gamma_{y_1}}$, and $\alpha_2^\star := \theta_2 \frac{\kappa_{z_2^c} \kappa_{y_2}}{\gamma_{c2}^c \gamma_{y_2}}$, the above model (S76) recasts to the same structure as the layered controller introduced in Multi-layer Autocatalytic Biomolecular Controllers: An Effective Solution.

#### Supplementary Note 4: Analysis of the Embedded Gene-Expression Control Example

Let us first recall from the equation (8) in Embedded Gene-Expression Control in the Presence of External Resource Loads: A Case Study, the closed-loop dynamic model of the considered embedded gene-expression control system. It follows that

$$\begin{cases} \dot{x}_1 = k^\star z_1 r + b_{p1}^\star r - \gamma_{p1} x_1, & \text{(S77a)} \\ \dot{z}_1 = \alpha_1^\star z_1 r - \theta_1 z_1 x_1 - \gamma_{c1} z_1, & \text{(S77b)} \\ \dot{z}_2 = \alpha_2^\star z_2 x_1 r - \gamma_{c2} z_2, & \text{(S77c)} \\ r = \dfrac{R_{tot}}{1 + w_1 z_1 + w_2 z_2 x_1}, & \text{(S77d)} \end{cases}$$

where the additional (resource) load module-related parts are excluded. Recall that, since the modules' resource-limited reactions that compete with $\mathbf{X_1}$, $\mathbf{Z_1}$, or $\mathbf{Z_2}$ are all assumed to be constant inflows, the overall effect of such excluded parts reflects only through changes in the parameter $Q_0$. Also recall that this parameter is encoded in the competition gains as well as the parameter $R_{tot}$. This system can admit four different equilibrium points (or more depending on the dynamic of the process network under control). We shall specify the equilibrium points by '$*$' superscripts and represent them by the tuple $[x_1^*, z_1^*, z_2^*]$. These are,

$$
\begin{aligned}
q_0 &= [b_{p1}^\star R_{tot}/\gamma_{p1}, 0, 0], \\
q_1 &= [\sqrt{\frac{\gamma_{c2} b_{p1}^\star}{\alpha_2^\star \gamma_{p1}}}, 0, \frac{R_{tot}\alpha_2^\star}{w_2\gamma_{c2}} - \frac{1}{w_2 x_1^*}], \\
q_2 &= [\frac{\alpha_1^\star R_{tot}}{\theta_1(1+w_1 z_1^*)} - \frac{\gamma_{c1}}{\theta_1}, \frac{\alpha_1^\star - \gamma_{c1}/R_{tot}}{\frac{k^\star\theta_1}{\gamma_{p1}} + \frac{w_1\gamma_{c1}}{R_{tot}}}, 0],
\end{aligned}
$$

and

$$
q_d = [\sqrt{(\frac{\gamma_{c1}}{2\theta_1})^2 + \frac{\alpha_1^\star}{\alpha_2^\star}\frac{\gamma_{c2}}{\theta_1}} - \frac{\gamma_{c1}}{2\theta_1}, \frac{\alpha_2^\star \gamma_{p1} {x_1^*}^2}{\gamma_{c2}k^\star} - \frac{b_{p1}^\star}{k^\star}, \frac{R_{tot}\alpha_2^\star}{w_2\gamma_{c2}} - \frac{(1+w_1 z_1^*)}{w_2 x_1^*}].
$$

The last one is considered as the desired equilibrium, which we wish to have as a stable fixed point. It is straightforward to show that $q_d$ is positive and, thus, feasible iff we have $\gamma_{p1}\alpha_2^\star {x_1^*}^2/\gamma_{c2} > b_{p1}^\star$ and $R_{tot}\alpha_2^\star x_1^* > \gamma_{c2}(1 + w_1 z_1^*)$, where

$$
x_1^* = \sqrt{(\gamma_{c1}/2\theta_1)^2 + \alpha_1^\star\gamma_{c2}/\alpha_2^\star\theta_1} - \gamma_{c1}/2\theta_1 =: s
$$

and $z_1^* = \alpha_2^\star\gamma_{p1}{x_1^*}^2/\gamma_{c2}k^\star - b_{p1}^\star/k^\star$. These two conditions, taken together, already impose an admissible range $\mathcal{S}_{\text{in}} \subset \mathbb{R}_{\geq 0}$ for the set-point $s$, which can be compactly written as $\mathcal{S}_{\text{in}} = \{s \in \mathbb{R}_{\geq 0} : s_{\text{inf}} < s < s_{\text{sup}}\}$, where

$$
s_{\text{inf}} := \max\{\sqrt{\frac{b_{p1}^\star\gamma_{c2}}{\alpha_2^\star\gamma_{p1}}}, \frac{k^\star}{2w_1\gamma_{p1}}(R_{tot} - \sqrt{R_{tot}^2 - \frac{4w_1\gamma_{p1}\gamma_{c2}}{k^\star\alpha_2^\star}})\}
$$

and

$$
s_{\text{sup}} := \frac{k^\star}{2w_1\gamma_{p1}}(R_{tot} + \sqrt{R_{tot}^2 - \frac{4w_1\gamma_{p1}\gamma_{c2}}{k^\star\alpha_2^\star}}).
$$

We begin by evaluating the Jacobian matrix of this system, $J(x_1, z_1, z_2) := [\Delta^{\mathrm{T}}\dot{x}_1, \Delta^{\mathrm{T}}\dot{z}_1, \Delta^{\mathrm{T}}\dot{z}_2]^{\mathrm{T}} \in \mathbb{R}^{3\times3}$, at some point $(x_1, z_1, z_2) = (x_1^*, z_1^*, z_2^*)$, which later represents the positive equilibrium point considered. The Jacobian may be written as follows

$$
J = \begin{bmatrix} -J_{11} & -J_{12} & -J_{13} \\ -g_{10} & -g_{11} & -g_{12} \\ -g_{20} & -g_{21} & -g_{22} \end{bmatrix}, \tag{S78}
$$

where

$$J_{11} = (b_{p1}^{\star} + k^{\star} z_1^*) r_{x_1} + \gamma_{p1},$$

$$J_{12} = (b_{p1}^{\star} + k^{\star} z_1^*) r_{z_1} - k^{\star} r^*,$$

$$J_{13} = (b_{p1}^{\star} + k^{\star} z_1^*) r_{z_2},$$

$$g_{10} = (\alpha_1^{\star} z_1^*) r_{x_1} + \theta_1 z_1^*,$$

$$g_{11} = (\alpha_1^{\star} z_1^*) r_{z_1} + \epsilon_{22},$$

$$g_{12} = (\alpha_1^{\star} z_1^*) r_{z_2},$$

$$g_{20} = \alpha_2^{\star} z_2^* (x_1^* r_{x_1} - r^*),$$

$$g_{21} = (\alpha_2^{\star} z_2^* x_1^*) r_{z_1},$$

$$g_{22} = (\alpha_2^{\star} z_2^* x_1^*) r_{z_2} + \epsilon_{33},$$

$$\epsilon_{22} = \gamma_{c1} + \theta_1 x_1^* - \alpha_1^{\star} r^*,$$

and

$$\epsilon_{33} = \gamma_{c2} - \alpha_2^{\star} x_1^* r^*.$$

Here, by abuse of notation, $r_v$ is employed to denote the negative of the derivative of $r(t)$ w.r.t. to the variable $v$ evaluated at $(x_1^*, z_1^*, z_2^*)$. In other expression,

$$r_{x_1} = {r^*}^2 (w_2 z_2^*)/R_{tot},$$

$$r_{z_1} = {r^*}^2 (w_1)/R_{tot},$$

and

$$r_{z_2} = {r^*}^2 (w_2 x_1^*)/R_{tot}.$$

Recall that $r^* = R_{tot}/(1 + w_1 z_1^* + w_2 z_2^* x_1^*)$.

Now, we find the characteristic polynomial of the closed-loop model linearized about $(x, z_1, z_2) = (x^*, z_1^*, z_2^*)$. We write out the characteristic polynomial of system (S77) as $p(s) := |sI - J|$, where $I$ is the identity matrix of appropriate size. From (S78), it follows that

$$p(s) = (s + g_{22}) \begin{vmatrix} s + J_{11} & J_{12} \\ g_{10} & s + g_{11} \end{vmatrix} - g_{12} \begin{vmatrix} s + J_{11} & J_{12} \\ g_{20} & g_{21} \end{vmatrix} + J_{13} \begin{vmatrix} g_{10} & s + g_{11} \\ g_{20} & g_{21} \end{vmatrix}.$$

Simplifying this result gives the representation $p(s) = s^3 + \eta_2 s^2 + \eta_1 s + \eta_0$, where

$$\eta_2 = g_{22} + g_{11} + J_{11},$$

$$\eta_1 = g_{22} J_{11} + g_{11} g_{22} - g_{12} g_{21} + J_{11} g_{11} - J_{12} g_{10} - J_{13} g_{20},$$

$$\eta_0 = g_{22} g_{11} J_{11} - J_{12} g_{10} g_{22} - g_{12} g_{21} J_{11} + J_{12} g_{12} g_{20} + J_{13} g_{10} g_{21} - J_{13} g_{20} g_{11}. \tag{S79}$$

The following proposition explores some sufficient conditions such that guarantee the stability of closed-loop system.

**Proposition S1.** *Let the positive equilibrium point $q_d = [s, z_1^*, z_2^*]$ of closed-loop system (S77) with*

$$s = \sqrt{(\frac{\gamma_{c1}}{2\theta_1})^2 + \frac{\alpha_1^\star}{\alpha_2^\star}\frac{\gamma_{c2}}{\theta_1}} - \frac{\gamma_{c1}}{2\theta_1}, \quad z_1^* = (\gamma_{p1}s/r^* - b_{p1}^\star)/k^\star, \quad z_2^* = (R_{tot}/r^* - (1 + w_1 z_1^*))/w_2 s$$

*be admissible, that is, let*

$$\gamma_{p1}\alpha_2^\star s^2/\gamma_{c2} > b_{p1}^\star, \tag{S80}$$

$$R_{tot}/r^* > (1 + w_1 z_1^*), \tag{S81}$$

*where $r^* = \gamma_{c2}/\alpha_2^\star s$ (or, equivalently, let $s \in \mathcal{S}_{\text{in}}$). Then, $q_d$ is locally (asymptotically) stable iff*

$$g_{22}(g_{22}J_{11} + J_{11}g_{11} - J_{13}g_{20}) + g_{11}(J_{11}g_{11} - J_{12}g_{10}) \\ + J_{11}(g_{22}J_{11} + g_{11}g_{22} + J_{11}g_{11} - J_{12}g_{10} - J_{13}g_{20}) > J_{12}g_{12}g_{20} + J_{13}g_{10}g_{21}.$$

*Further, it is sufficient to satisfy this condition if $2\gamma_{c2} \le \gamma_{p1}$.*

*Proof.* From Routh-Hurwitz criterion, we know that the eigenvalues of $J$ evaluated at $(x_1, z_1, z_2) = [x_1^*, z_1^*, z_2^*]$ have negative real parts iff the coefficients $\eta_0$, $\eta_1$, and $\eta_2$ in (S79) are all strictly positive and satisfy $\eta_2\eta_1 > \eta_0$. All the elements of $J$ in (S78) are negative except two elements: the most left-bottom entry and potentially the one at the first row and second column. Note that it follows from the definition of $r$ which entails $v^* r_v$ to be always less than $r^*$.

An equilibrium with $x_1^* = s$ sets both $\epsilon_{22}$ and $\epsilon_{33}$ to zero (and so does $g_{11}g_{22}$ equal to $g_{12}g_{21}$). Thus, we have that $\eta_2$ is always positive,

$$\eta_1 = J_{11}(g_{22} + g_{11}) - J_{12}g_{10} - J_{13}g_{20},$$

and

$$\eta_0 = J_{12}(g_{12}g_{20} - g_{10}g_{22}) + J_{13}(g_{10}g_{21} - g_{20}g_{11}).$$

Substituting $g_{ij}$ and $J_{ij}$ appeared in the latter summarizes $\eta_0$ as

$$\eta_0 = k^\star r^* \alpha_2^\star z_1^* z_2^* r_{z_2}(\alpha_1^\star r^* + \theta_1 x_1^*), \tag{S82}$$

which is always positive. Hence, we are left to find conditions guaranteeing $\eta_1 > 0$. Expanding its two terms $J_{11}g_{11}$ and $-J_{12}g_{10}$, we rewrite $\eta_1$ as

$$\eta_1 = J_{11}g_{22} - J_{13}g_{20} + \gamma_{p1}r_{z_1}\alpha_1^\star z_1^* + k^\star r^* r_{x_1}\alpha_1^\star z_1^* + k^\star r^* \theta_1 z_1^* - r_{z_1}(b_{p_1}^\star + k^\star z_1^*)\theta_1 z_1^*. \tag{S83}$$

The only negative term in (S83) is the last one. We again use the fact that $r^* - r_{z_1}z_1^*$ is always nonnegative. Thus, $k^\star \theta_1^\star z_1^*(r^* -$

$r_{z_1} z_1^*) \geq 0$. We are left to find a positive term from (S83) that is greater equal than $r_{z_1} b_{p_1}^\star \theta_1 z_1^*$. The derivative of $s = \sqrt{(\gamma_{c1}/2\theta_1)^2 + \alpha_1^\star \gamma_{c2}/\alpha_2^\star \theta_1} - \gamma_{c1}/2\theta_1$ w.r.t. the parameter $\gamma_{c1}$ is strictly negative and, hence, $\sqrt{\alpha_1^\star \gamma_{c2}/\alpha_2^\star \theta_1}$ is a least upper bound for the range that $s$ can take regarding non-zero values for $\theta_1, \alpha_1^\star, \alpha_2^\star, \gamma_{c2}$. Upon choosing the third term $\gamma_{p1} r_{z_1} \alpha_1^\star z_1^*$, one can whereby show that the condition (S80) in the statement of proposition already implies $r_{z_1} z_1^* (\gamma_{p1} \alpha_1^\star - b_{p_1}^\star \theta_1) \geq 0$. In conclusion, $\eta_1$ will be always positive, as there exists at least one another term, e.g., $k^\star r^* r_{x_1} \alpha_1^\star z_1^*$, to keep it non-zero.

Now, it is just remained to find other conditions satisfying $\eta_2 \eta_1 > \eta_0$. Consider the term $k^\star r^* r_{x_1} \alpha_1^\star z_1^*$ from $\eta_1$ in (S83) and the term $(g_{22} + J_{11})$ from $\eta_2$ in (S79). We proceed by taking $k^\star r^* r_{x_1} \alpha_1^\star z_1^* (g_{22} + J_{11}) \geq \eta_0$ as a sufficient condition satisfying $\eta_2 \eta_1 > \eta_0$. Hence, after some rearrangements, we will get to show that

$$g_{22} + J_{11} > \alpha_2^\star x_1^* r^* (1 + \theta_1 x_1^* / \alpha_1^\star r^*). \tag{S84}$$

On substituting the expansion of $g_{22} + J_{11}$ with noting that $b_{p1}^\star + k^\star z_1^* = \gamma_{p1} x_1^* / r^*$ and $x_1^* r^* = \gamma_{c2}/\alpha_2^\star$, (S84) reduces to

$$\frac{2\gamma_{p1}}{\gamma_{c2}} > \frac{1 + w_1 z_1^*}{\alpha_2^\star x_1^* R_{tot}} (\gamma_{p1} + \gamma_{c2}) + \frac{\alpha_2^\star \theta_1}{\alpha_1^\star \gamma_{c2}} {x_1^*}^2. \tag{S85}$$

From (S81) we know that the term $(1 + w_1 z_1^*)/\alpha_2^\star x_1^* R_{tot}$ is always less than $1/\gamma_{c2}$. As discussed already, $\sqrt{\alpha_1^\star \gamma_{c2}/\alpha_2^\star \theta_1}$ is an upper bound for $x_1^*$ and, thus, $\sup\{\alpha_2^\star \theta_1 {x_1^*}^2 / \alpha_1^\star \gamma_{c2}\} = 1$. Putting them together, having the only condition $\gamma_{p1}/\gamma_{c2} \geq 2$ met is sufficient to conclude the local (asymptotic) stability of equilibrium $q_d$, which completes the proof. ∎

#### Supplementary Note 5: Impact of Saturation and Constant Controller Degradation/Dilution on the Ratio Manifold

In this note, we first briefly discuss the modified layered autocatalytic motif introduced in Ratiometric Control using a Layered Autocatalytic Integral Feedback Strategy, which is capable of achieving ratiometric adaptation. We then analyze how accounting for saturation and constant degradation/dilution of the controller impacts the controller motif's ability to achieve adaptation.

**CRN Representation of the Proposed Motif:** A possible control law capable of achieving robust perfect ratiometric adaptation can be biochemically realized by the following set of reactions

$$\begin{aligned}
\mathcal{R}_{c1} &: \mathbf{Z_1} + \mathbf{X_2} \xrightarrow{\alpha_1} \mathbf{X_2} + 2\mathbf{Z_1}, \\
\mathcal{R}_{c2} &: \mathbf{X_1} + \mathbf{Z_1} \xrightarrow{\theta_1} \mathbf{X_1}, \\
\mathcal{R}_{c3} &: \mathbf{Z_2} + \mathbf{X_1} \xrightarrow{\alpha_2} \mathbf{X_1} + 2\mathbf{Z_2}, \\
\mathcal{R}_{c4} &: \mathbf{X_2} + \mathbf{Z_2} \xrightarrow{\theta_2} \mathbf{X_2},
\end{aligned}$$

in which the reactions $\mathcal{R}_{c1}$ and $\mathcal{R}_{c3}$ are resource-limited bimolecular reactions that are involved in the competition for a common resource $\mathbf{R}$. One may write the closed-loop dynamic model associated to this control circuit as the equations (11) in Ratiometric Control using a Layered Autocatalytic Integral Feedback Strategy. In this model, we have employed the nonlinear maps $H_i$ and $G_i$ to deal with possible saturation effects on the controller dynamics. We assume these functions to be of class $\mathcal{K}$

functions on their admissible domains. They are one-to-one and strictly monotonic over the non-negative real axis. Moreover, we do assume in (11) that the effect of controller degradation or dilution is negligible (later in this note, however, we separately account for such an effect). This may be seen as a fair assumption in certain design cases, such as when more stable proteins are chosen for the controller species compared to process ones or when ribonucleases are upregulated and degradation tags are utilized to synthetically promote the degradation of process species, depending on the context. If the resource competition for $\mathbf{R}$ between the species $\mathbf{Z_1}$, $\mathbf{Z_2}$, and the rest of the involved components is intracellular, then one can refer to A Mathematical Framework to Model Intracellular Resource Competition for the following description for the available resources

$$r = \frac{R_{tot}}{(1 + w_{p1}x + x^{\mathrm{T}}w_{p2}x + w_1 z_1 x_2 + w_2 z_2 x_1)},$$

in which, without loss of generality, we have assumed that the bimolecular reactions $\mathcal{R}_{c1}$ and $\mathcal{R}_{c3}$ are of the first scenario. As with the previous sections, the way the controller acts on the plant is case-specific. We have thus not specified the actuation reactions, leaving them to be chosen by the designer. Depending on the circuit under consideration or constraints arising from the actual biological limitations, one may opt for choosing from a range of actuation mechanisms. These can be activatory or inhibitory, may influence one or more process species—including both $\mathbf{X_1}$ and $\mathbf{X_2}$ as well—or may not even involve direct actuation, instead relying on the resource coupling to serve as an indirect actuator. In (11a), a function $u : \mathbb{R}_{\geq 0}^{n+2} \to \mathbb{R}^n$ is employed to represent for such (stabilizing) control inputs, taken together.

**Analysis of the Ratio Manifold in Scenarios Where the Saturation and Dilution Assumptions Are Violated:** Consider the following reformulated dynamic model from (11) noted in Ratiometric Control using a Layered Autocatalytic Integral Feedback Strategy

$$\begin{cases} \dot{x} = f(x, r) + u(z_1, z_2, x), & \text{(S86a)} \\ \dot{z}_1 = \alpha_1^\star z_1 G_1(x_2) r - \theta_1 z_1 H_1(x_1) - \gamma_{c1} z_1, & \text{(S86b)} \\ \dot{z}_2 = \alpha_2^\star z_2 G_2(x_1) r - \theta_2 z_2 H_2(x_2) - \gamma_{c2} z_2, & \text{(S86c)} \end{cases}$$

where the functions $H_i$ and $G_i$ are to encapsulate possible saturation effects while $\gamma_{ci}$ are to reflect the effect of constant controller degradation/dilution. At steady state, the following relation holds

$$x_1^* = \mathcal{F}^{-1}\left(\frac{\alpha_1^\star \theta_2}{\alpha_2^\star \theta_1} G_1(x_2^*) H_2(x_2^*) + \frac{\alpha_1^\star \gamma_{c2}}{\alpha_2^\star \theta_1} G_1(x_2^*)\right), \tag{S87}$$

where

$$\mathcal{F}(\rho) := G_2(\rho)\left(H_1(\rho) + \frac{\gamma_{c1}}{\theta_1}\right)$$

and $\mathcal{F}^{-1}$ refers to its inverse function, which, again, exists and is well-defined on the positive axis. Here, we consider four different cases, classified based on whether they involve saturation or dilution. We will explicitly derive ratio manifolds that describe the steady-state $x_1^*$-$x_2^*$ relationship by substituting the saturation terms with biologically meaningful Michaelis-Menten terms.

**Case I (with saturation, with constant controller degradation/dilution).** We deal with the case of saturation by setting $G_i(\rho) := \dfrac{\rho}{1 + K_{G_i}\rho}$ and $H_i(\rho) := \dfrac{\rho}{1 + K_{H_i}\rho}$ for every $i$. We do not impose conditions on $\gamma_{ci}$s. To find the solution(s) for

$\mathcal{F}^{-1}(\rho)$, we have to solve the zeros of a quadratic equation in the form $a(\mathcal{F}^{-1})^2 + b\mathcal{F}^{-1} + 1 = 0$, in which $a$ and $b$ take the forms $a = K_{G_2}K_{H_1} - (1 + K_{H_1}\gamma_{c1}/\theta_1)/\rho$ and $b = K_{G_2} + K_{H_1} - \gamma_{c1}/\theta_1\rho$. After some algebra, we arrive at the following as the only positive solution for $\mathcal{F}^{-1}(\rho)$

$$\mathcal{F}^{-1}(\rho) = \frac{\dfrac{\gamma_{c1}/\theta_1\rho - (K_{G_2} + K_{H_1})}{2} - \sqrt{\left(\dfrac{\gamma_{c1}/\theta_1\rho - (K_{G_2} + K_{H_1})}{2}\right)^2 - K_{G_2}K_{H_1} + \dfrac{1 + K_{H_1}\gamma_{c1}/\theta_1}{\rho}}}{K_{G_2}K_{H_1} - \dfrac{1 + K_{H_1}\gamma_{c1}/\theta_1}{\rho}}, \tag{S88}$$

provided the condition

$$\rho\left((\frac{K_{G_2} - K_{H_1}}{2})^2 + (\frac{\gamma_{c1}}{2\theta_1\rho})^2 + \frac{\gamma_{c1}}{2\theta_1\rho}K_{H_1}\right) > \frac{\gamma_{c1}}{2\theta_1\rho}K_{G_2} - 1 \tag{C1}$$

is satisfied. Additionally, in the cases where

$$K_{G_2} + K_{H_1} > \gamma_{c1}/\theta_1\rho,$$

the condition

$$K_{G_2}K_{H_1} < (1 + K_{H_1}\gamma_{c1}/\theta_1)/\rho \tag{C2}$$

has to be satisfied too. Note that the solution is unique as the inequality $K_{H_1}b > a$ always holds. On substituting

$$\frac{\alpha_1^\star}{\alpha_2^\star\theta_1}\frac{x_2^*}{1 + K_{G_1}x_2^*}\left(\frac{\theta_2 x_2^*}{1 + K_{H_2}x_2^*} + \gamma_{c2}\right) \tag{S89}$$

for $\rho$, one can easily compute the explicit form of the ratio manifold from (S87). In particular, the conditions in (C1)-(C2) always hold if $G_2(x_1^*)$ takes linear forms. Moreover, there always exists a domain of validity for these two conditions if the effect of controller degradation is taken as negligible. In which case, there always exist a sufficiently small scalar $x_2^{\max}$ such that for every $x_2^* \in (0, x_2^{\max}]$ the ratio manifold can be followed from (S88) by substituting $\dfrac{\alpha_1^\star\theta_2}{\alpha_2^\star\theta_1}{x_2^*}^2/(1 + K_{G_1}x_2^*)(1 + K_{H_2}x_2^*)$ for $\rho$. Such a domain exists as the equation in (S89) is monotonically increasing in $x_2^*$ and remains positive for every $x_2^* > 0$. Of note, if both of the two conditions (C1)-(C2) are violated, no positive ratio between $x_1^*$ and $x_2^*$ is conceivable.

**Case II (with saturation, no constant controller degradation/dilution).** We keep the saturation settings as with the previous case, $G_i(\rho) := \dfrac{\rho}{1 + K_{G_i}\rho}$ and $H_i(\rho) := \dfrac{\rho}{1 + K_{H_i}\rho}$ for every $i$. We set, however, $\gamma_{ci} = 0$. From (S88), we find the positive solution for $\mathcal{F}^{-1}$ as below

$$\mathcal{F}^{-1}(\rho) = \frac{\sqrt{(\dfrac{K_{G_2} - K_{H_1}}{2})^2 + \dfrac{1}{\rho}} + \dfrac{K_{G_2} + K_{H_1}}{2}}{1/\rho - K_{G_2}K_{H_1}}, \qquad \text{if } \rho < 1/K_{G_2}K_{H_1}.$$

This finally leads us to the relation

$$x_1^* = \frac{\sqrt{(\dfrac{K_{G_2} - K_{H_1}}{2})^2 + \dfrac{(1 + K_{G_1}x_2^*)(1 + K_{H_2}x_2^*)}{\alpha_1^\star\theta_2{x_2^*}^2/\alpha_2^\star\theta_1}} + \dfrac{K_{G_2} + K_{H_1}}{2}}{\dfrac{(1 + K_{G_1}x_2^*)(1 + K_{H_2}x_2^*)}{\alpha_1^\star\theta_2{x_2^*}^2/\alpha_2^\star\theta_1} - K_{G_2}K_{H_1}}, \quad \text{if } x_2^* < x_2^{\max}. \tag{S90}$$

The positive constant $x_2^{\max}$ is defined as below

$$
\begin{aligned}
x_2^{\max} &= \frac{\frac{K_{G_1}+K_{H_2}}{2} + \sqrt{(\frac{K_{G_1}+K_{H_2}}{2})^2 + K_{G_2}K_{H_1}\frac{\alpha_1^\star\theta_2}{\alpha_2^\star\theta_1} - K_{G_1}K_{H_2}}}{K_{G_2}K_{H_1}\frac{\alpha_1^\star\theta_2}{\alpha_2^\star\theta_1} - K_{G_1}K_{H_2}} && \text{if } K_{G_2}K_{H_1}\frac{\alpha_1^\star\theta_2}{\alpha_2^\star\theta_1} > K_{G_1}K_{H_2}, \\
x_2^{\max} &\equiv \infty && \text{elsewhere.}
\end{aligned}
$$

**Case III (no saturation, with constant controller degradation/dilution).** This case we find by setting $G_i(\rho) := \rho$ and $H_i(\rho) := \rho$ for every $i$, but leaving $\gamma_{ci} \neq 0$. Simple calculation finds $\mathcal{F}^{-1}(\rho) = \sqrt{(\frac{\gamma_{c1}}{2\theta_1})^2 + \rho} - \frac{\gamma_{c1}}{2\theta_1}$ and, hence,

$$
x_1^* = \sqrt{(\frac{\gamma_{c1}}{2\theta_1})^2 + \frac{\alpha_1^\star}{\alpha_2^\star\theta_1}(\theta_2 x_2^* + \gamma_{c2})x_2^*} - \frac{\gamma_{c1}}{2\theta_1}. \tag{S91}
$$

Note, the above expression reduces to the linear expression

$$
x_1^* = \frac{\alpha_1^\star\gamma_{c2}}{\alpha_2^\star\gamma_{c1}}x_2^*
$$

if the parametric condition

$$
\frac{\gamma_{c1}}{\gamma_{c2}} = \sqrt{\frac{\alpha_2^\star\theta_2}{\alpha_1^\star\theta_1}}
$$

is fulfilled. However, this linear formulation, which implies perfect ratiometric adaptation, is not structurally robust, as any small perturbation in the controller parameters could shift it to an imperfect ratiometric compensation. A robust structure for achieving perfect ratiometric adaptation, on the other hand, results from considering the controller dilution effects negligible, the next case.

**Case IV (no saturation, no constant controller degradation/dilution).** This case we find by setting $G_i(\rho) := \rho$, $H_i(\rho) := \rho$ and $\gamma_{ci} = 0$ for every $i$. From (S88), it follows that $\mathcal{F}^{-1}(\rho) = \sqrt{\rho}$. The equation in (S89) also simplifies to $\frac{\alpha_1^\star\theta_2}{\alpha_2^\star\theta_1}x_2^{*2}$. Subsequently, we obtain for the ratio manifold

$$
x_1^* = \sqrt{\frac{\alpha_1^\star\theta_2}{\alpha_2^\star\theta_1}}x_2^*. \tag{S92}
$$

This is the same equation we reported in (13) in the main text, Ratiometric Control using a Layered Autocatalytic Integral Feedback Strategy. Numerical illustrations of the ratio manifold for the above four cases are provided in Figure S9.

From Case III, one can use the results therein to recover the results previously obtained for the regulation tasks. For example, if we substitute the constant unity term for $x_2^*$ in the equation (S91), we would recover the dynamic structure in (6) from the section Multi-layer Autocatalytic Biomolecular Controllers: An Effective Solution of the main text and would also recover the steady-state relation form reported in (7) therein. Similarly, one can use the above results to consider the effect of saturation on the multicellular control circuits employed in the main text for regulation tasks. For example, consider the introduced motif for the case of a general plant as described by the equations (9), discussed in Multicellular Realization of the Layered Autocatalytic

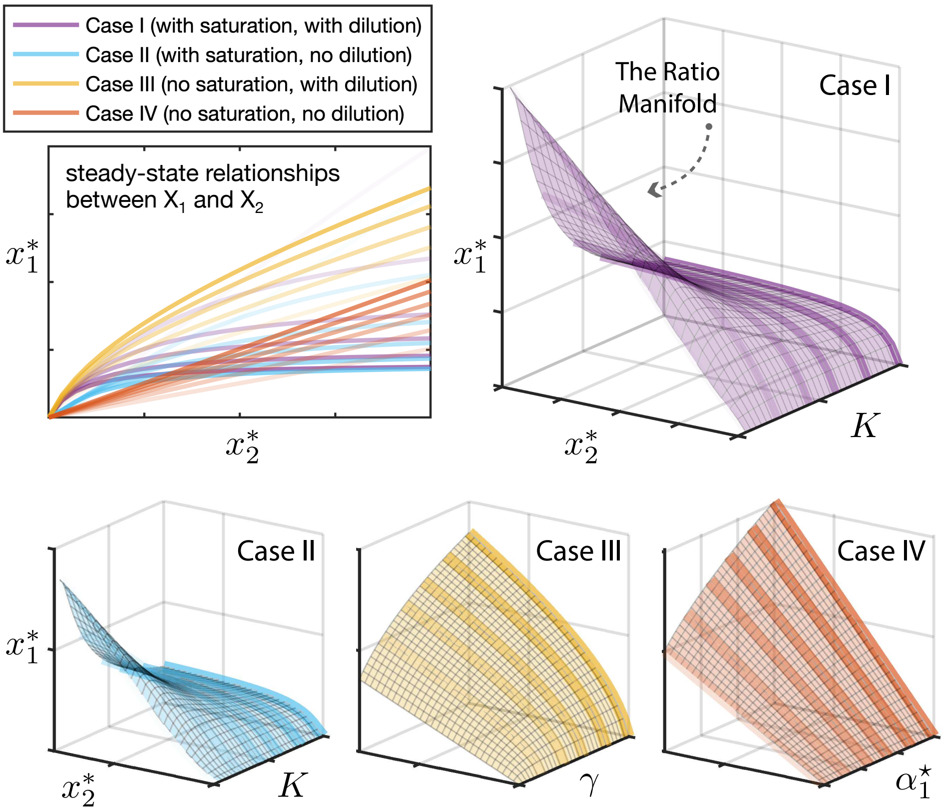

Figure S9: **Accounting for saturating dynamics and constant degradation/dilution of the controller species may lead to imperfect ratiometric adaptation results.** We numerically solve for the relationships between the steady-state concentrations of $\mathbf{X_1}$ and $\mathbf{X_2}$ obtained from the generalized dynamical system (S86) using a set of parameters. The resulting ratio manifolds are shown for the four cases discussed in Supplementary Note 5: Impact of Saturation and Constant Controller Degradation/Dilution on the Ratio Manifold. We set $K_{G_2}$ to zero to meet both of the conditions (C1)-(C2) reported therein. For the other saturation-related parameters, $K_{H_1}$, $K_{G_1}$, and $K_{H_2}$, we set them all three equal to the same value $K$. Setting $K = 0$ effectively disregards the saturation effects. For the controller's constant degradation/dilution effects, both $\gamma_{c1}$ and $\gamma_{c2}$ are set equal to the same value $\gamma$. In Case I, we vary $K$ while $\gamma$ is non-zero. In Case II, we conduct the same simulation with $\gamma = 0$. For Case III, we set $K = 0$ and vary $\gamma$ instead. For the last case, Case IV, associated with the regime capable of achieving robust perfect ratiometric adaptation, we set both $K$ and $\gamma$ to zero and vary another design parameter, $\alpha_1^\star$. The two-dimensional plot illustrates, for each case, six different sample trajectories in the $x_1^* - x_2^*$ space. Notably, only Case IV, which does not consider saturation and dilution, consistently results in pure linear forms within this 2D subspace, implying perfect ratiometric adaptation. In contrast, the other cases demonstrate imperfect ratiometric responses; however, the $x_1^* - x_2^*$ relationship is still solely determined by the controller parameters, thereby offering robustness with respect to stimuli influencing the process network.

Integrator Motif. Defining $r := 1 - (N_1 + N_2 + \sum_{i=3}^{m+2} c_{1i}N_i)/N_{\mathrm{m}}$, one can recast (9) as follows

$$\begin{cases} \dot{x} = f(x, N_1, N_2), & \text{(S93a)} \\ \dot{N}_1 = \mu_1 N_1 r - (\theta_1 x_n + \gamma_{c1}) N_1, & \text{(S93b)} \\ \dot{N}_2 = \rho_{x_n} G_2(x_n) N_2 r - \gamma_{c2} N_2. & \text{(S93c)} \end{cases}$$

It is easy to see that the above model follows the same structure as that of (S86), if we take linear forms for the three functions $G_1$, $H_1$, and $H_2$, if we leave $G_2$ to take the aforementioned Michaelis-Menten forms, and if we further set $\gamma_{c2} = 0$ and $x_2^* \equiv 1$. This way, we arrive at the following expression for $x_n^*$ by revoking (S88)

$$x_n^* = \mathcal{F}^{-1}\left(\frac{\mu_1 \gamma_{c2}}{\rho_{x_n} \theta_1}\right)$$

where

$$\mathcal{F}^{-1}(\rho) := \frac{\rho K_{G_2} - \gamma_{c1}/\theta_1}{2} + \sqrt{\left(\frac{\gamma_{c1}/\theta_1 - \rho K_{G_2}}{2}\right)^2 + \rho}.$$

Combing on the above finding leads to the simplified expression

$$x_n^* = \frac{\mu_1\gamma_{c2}K_{G_2} - \rho_{x_n}\gamma_{c1}}{2\theta_1\rho_{x_n}} + \sqrt{\left(\frac{\mu_1\gamma_{c2}K_{G_2} - \rho_{x_n}\gamma_{c1}}{2\theta_1\rho_{x_n}}\right)^2 + \frac{\mu_1\gamma_{c2}}{\rho_{x_n}\theta_1}}.$$

If we cancel the effect of saturation on $G_2$ by setting $K_{G_2} = 0$, it would result in the same set-point expression as reported in Multicellular Realization of the Layered Autocatalytic Integrator Motif. We would like to note that these derivations are valid as long as both of the conditions (C1)-(C2) are met. The second condition is always met as we have set $K_{H_1}$ to zero. For the first condition, however, one needs to check it for every given $K_{G_2}$. Speaking of sufficient conditions, it always holds provided that $K_{G_2}$ is chosen sufficiently small to meet $\frac{\gamma_{c1}}{2\theta_1\rho}K_{G_2} < 1$.

#### Supplementary Note 6: Further Details on Numerical Results and Examples

Here, we mainly provide the numerical values of the parameters used to obtain our simulation results in certain figures.

**Figure 3.** The initial conditions are $x_1^0 = 0$ and $z_1^0 = 0.1$, and the nominal parameters include $k^\star = 1, b_{p1}^\star = 0.1, \gamma_{p1} = 0.5, \alpha_1^\star = 20, \theta_1 = 0.1, \gamma_{c1} = 1, R_{tot} = 100, w_1 = 1$. The considered model for the simulation part (**D**) follows the dynamics of (5) with $f(x_1, r) = b_{p1}^\star r - \gamma_{p1}x_1$, in which the dynamics of the transcriptional parts are not explicitly accounted for, being assumed to have rapidly reached some quasi-steady states. Note that this choice of descriptor function for the process dynamics automatically sets both the competition gains $w_{p1}$ and $w_{p2}$ to zero. We remark that the expression for the normalized error $e_p$ follows $e_p = 1 + (k_2^\star - k_1^\star)/(k_1^\star + k_1^\star k_2^\star/\vartheta w_1)$ in the limit $R_{tot} \to \infty$, where $k_1^\star$ and $k_2^\star$ are the pre-perturbed and perturbed values of $k^\star$, respectively, and $\vartheta := \gamma_{p1}\alpha_1^\star/\theta_1 - b_{p1}^\star$.

**Figure 6.** The initial conditions are set to $x_1(0) = 0$ and $N_{1,2}(0) = 10^{-7}\,\mathrm{ml}^{-1}$, while $c_{1i} = 0$. Default parameters used to obtain the simulation results are $k = 5 \times 10^{-7}\,\mathrm{nM\,ml\,h}^{-1}, \gamma_{p1} = 0.3\,\mathrm{h}^{-1}, \gamma = 0.05\,\mathrm{h}^{-1}, \mu_1 = 0.8\,\mathrm{h}^{-1}, \rho_{x_1} = 10^{-3}\,\mathrm{nM}^{-1}\mathrm{h}^{-1}, \theta_1 = 10^{-4}\,\mathrm{nM}^{-1}\mathrm{h}^{-1}, N_\mathrm{m} = 10^9\,\mathrm{ml}^{-1}$.

**Figure 7.** The initial conditions: $a_{1,3}(0) = 0, N_{1,2,3}(0) = 10^{-7}\,\mathrm{ml}^{-1}$. Typical parameters are $\rho_1 = 0.1\,\mathrm{nM}^{-1}\mathrm{h}^{-1}, \rho_3 = 10^{-3}\,\mathrm{nM}^{-1}\mathrm{h}^{-1}, k_{a1} = 5 \times 10^{-7}\,\mathrm{nM\,ml\,h}^{-1}, k_{a3} = 5 \times 10^{-7}\,\mathrm{nM\,ml\,h}^{-1}, \gamma_{a1} = 0.3\,\mathrm{h}^{-1}, \gamma_{a3} = 0.3\,\mathrm{h}^{-1}$, the rest are kept consistent with those of Figure 6.

**Figure 8.** The associated closed-loop dynamic model can be described as

$$\begin{cases} \dot{m}_{x_1} = k^\star z_1 r_1 - \gamma_{m_{x_1}} m_{x_1}, & \text{(S94a)} \\ \dot{m}_\mathrm{P} = b_\mathrm{P}^\star r_1 - \gamma_{m_\mathrm{P}} m_\mathrm{P}, & \text{(S94b)} \\ \dot{m}_{z_1} = \kappa_1^\star z_1 x_2 r_1 - \gamma_{m_{z_1}} m_{z_1}, & \text{(S94c)} \\ \dot{m}_{z_2} = \kappa_2^\star z_2 x_1 r_1 - \gamma_{m_{z_2}} m_{z_2}, & \text{(S94d)} \end{cases}$$

for the transcriptional components,

$$\begin{cases} \dot{x}_1 = \eta_0^\star m_{x_1} r_2 + \eta_1^\star m_{\mathrm{P}} r_2 - \gamma_{x_1} x_1, & \text{(S94e)} \\ \dot{x}_2 = \eta_2^\star m_{\mathrm{P}} r_2 - \gamma_{x_2} x_2, & \text{(S94f)} \\ \dot{z}_1 = \lambda_1^\star m_{z_1} r_2 - \theta_1 z_1 x_1, & \text{(S94g)} \\ \dot{z}_2 = \lambda_2^\star m_{z_2} r_2 - \theta_2 z_2 x_2, & \text{(S94h)} \end{cases}$$

for the translational components, and

$$\begin{cases} r_1 = \dfrac{R_{tot_1}}{1 + w_{11} z_1 + w_{12} z_1 x_2 + w_{13} z_2 x_1}, & \text{(S94i)} \\ r_2 = \dfrac{R_{tot_2}}{1 + w_{21} m_{x_1} + w_{22} m_{\mathrm{P}} + w_{23} m_{z_1} + w_{24} m_{z_2}}, & \text{(S94j)} \end{cases}$$

for the resource terms that appear because of competitive production reactions. Refer to Figure 8B for the closed-loop circuit diagram and see A Mathematical Framework to Model Intracellular Resource Competition for detailed information regarding the resource-limited modelings. Define $\alpha_i^\star := \lambda_i^\star \kappa_i^\star / \gamma_{m_{z_i}}$ for $i \in \{1, 2\}$. Then, solving for $x_1^* / x_2^*$ results in (13) from Ratiometric Control using a Layered Autocatalytic Integral Feedback Strategy. From the above model, it is easy to compute that $r_1^* r_2^* = \theta_1 \Gamma / \alpha_1^\star$. Initial conditions are set to 0.1 for $z_1$ and $z_2$ and zero for the remaining state variables. Nominal values for the parameters: $\Gamma = 2, k^\star = 10, b_{\mathrm{P}}^\star = 0.2, \kappa_{1,2}^\star = 1, \eta_{0,1,2}^\star = 1, \lambda_1^\star = 2, \lambda_2^\star = 1, \theta_1 = 2, \theta_2 = 4, \gamma_{x_1} = 0.5, R_{tot_1} = 50, R_{tot_2} = 100$, with the mRNA degradation rates set to 0.1.

**Figure 9.** The subsequent equation is used for the closed-loop system dynamics

$$\begin{cases} \dot{a}_i = k_{ai} N_i - \gamma_{ai} a_i \qquad \text{for } i \in \{1, \cdots, 4\}, \\ \dot{N}_3 = \rho_1 a_1 N_3 (1 - \dfrac{c_{31}(N_1 + N_2) + N_3 + c_{34} N_4}{N_{\mathrm{m3}}}) - \gamma N_3, & \text{(S95a)} \\ \dot{N}_4 = \rho_2 a_2 N_4 (1 - \dfrac{c_{41}(N_1 + N_2) + c_{43} N_3 + N_4}{N_{\mathrm{m4}}}) - \gamma N_4, & \text{(S95b)} \\ \dot{N}_1 = \rho_4 a_4 N_1 (1 - \dfrac{N_1 + N_2 + c_{13} N_3 + c_{14} N_4}{N_{\mathrm{m}}}) - \theta_1 a_3 N_1, & \text{(S95c)} \\ \dot{N}_2 = \rho_3 a_3 N_2 (1 - \dfrac{N_1 + N_2 + c_{13} N_3 + c_{14} N_4}{N_{\mathrm{m}}}) - \theta_2 a_4 N_2. & \text{(S95d)} \end{cases}$$

For a set of parameters in which $N_{\mathrm{m3}} = N_{\mathrm{m}}$, $N_{\mathrm{m4}} = N_{\mathrm{m}}$, and all $c_{ij}$ are set to unity, we list the according steady-state relationships as below

$$N_1^* = \frac{\gamma_{a1} \gamma}{k_{a1} \rho_1} \sqrt{\frac{\rho_3 \rho_4}{\theta_1 \theta_2}},$$
$$N_2^* = \frac{\gamma_{a2} \gamma}{k_{a2} \rho_2} \sqrt{\frac{\rho_3 \rho_4}{\theta_1 \theta_2}},$$
$$N_3^* = \frac{\sqrt{\dfrac{\rho_4 \theta_2}{\rho_3 \theta_1} \dfrac{k_{a4} \gamma_{a3}}{k_{a3} \gamma_{a4}}}}{1 + \sqrt{\dfrac{\rho_4 \theta_2}{\rho_3 \theta_1} \dfrac{k_{a4} \gamma_{a3}}{k_{a3} \gamma_{a4}}}} \left( (1 - \sqrt{\frac{\theta_1 \theta_2}{\rho_3 \rho_4}}) N_{\mathrm{m}} - (\frac{\gamma_{a1} \gamma}{k_{a1} \rho_1} \sqrt{\frac{\rho_3 \rho_4}{\theta_1 \theta_2}} + \frac{\gamma_{a2} \gamma}{k_{a2} \rho_2} \sqrt{\frac{\rho_3 \rho_4}{\theta_1 \theta_2}}) \right),$$

$$N_4^* = \frac{(1-\sqrt{\frac{\theta_1\theta_2}{\rho_3\rho_4}})N_{\rm m} - (\frac{\gamma_{a1}\gamma}{k_{a1}\rho_1}\sqrt{\frac{\rho_3\rho_4}{\theta_1\theta_2}} + \frac{\gamma_{a2}\gamma}{k_{a2}\rho_2}\sqrt{\frac{\rho_3\rho_4}{\theta_1\theta_2}})}{1+\sqrt{\frac{\rho_4\theta_2}{\rho_3\theta_1}\frac{k_{a4}\gamma_{a3}}{k_{a3}\gamma_{a4}}}},$$

$$a_i^* = k_{ai}N_i^*/\gamma_{ai}.$$

From the above expressions, it is easy to check that the reference ratio (the induced set-point) is admissible if the two conditions

$$\rho_3\rho_4 > \theta_1\theta_2 \tag{S96a}$$

and

$$\sqrt{\theta_1\theta_2/\rho_3\rho_4} - \theta_1\theta_2/\rho_3\rho_4 > \gamma(\gamma_{a1}/k_{a1}\rho_1 + \gamma_{a2}/k_{a2}\rho_2)/N_{\rm m} \tag{S96b}$$

are satisfied. Nominal parameters used to obtain the simulation results are $\rho_1 = 0.005\,{\rm nM^{-1}h^{-1}}$, $\rho_2 = 0.0025\,{\rm nM^{-1}h^{-1}}$, $\rho_3 = 10^{-4}\,{\rm nM^{-1}h^{-1}}$, $\rho_4 = 0.0064\,{\rm nM^{-1}h^{-1}}$, $k_{a1} = 5\times10^{-7}\,{\rm nM\,ml\,h^{-1}}$, $k_{a2} = 6\times10^{-7}\,{\rm nM\,ml\,h^{-1}}$, $k_{a3} = 4\times10^{-7}\,{\rm nM\,ml\,h^{-1}}$, $k_{a4} = 1\times10^{-7}\,{\rm nM\,ml\,h^{-1}}$, $\gamma_{ai} = 0.3\,{\rm h^{-1}}$, $\gamma = 0.03\,{\rm h^{-1}}$, $\theta_1 = 10^{-4}\,{\rm nM^{-1}h^{-1}}$, $\theta_2 = 10^{-4}\,{\rm nM^{-1}h^{-1}}$, $N_{\rm m} = 10^9\,{\rm ml^{-1}}$.
